# Phospho-signaling couples polar asymmetry and proteolysis within a membraneless microdomain in *C. crescentus*

**DOI:** 10.1101/2023.08.19.553945

**Authors:** Yasin M Ahmed, Grant R Bowman

## Abstract

Asymmetric cell division in bacteria is achieved through cell polarization, where regulatory proteins are directed to specific cell poles. Curiously, both poles contain a membraneless microdomain, established by the polar assembly hub PopZ, through most of the cell cycle, yet many PopZ clients are unipolar and transiently localized. We find that PopZ’s interaction with the response regulator CpdR is controlled by phosphorylation, via the histidine kinase CckA. Phosphorylated CpdR does not interact with PopZ and is not localized to cell poles. At poles where CckA acts as a phosphatase, de-phosphorylated CpdR binds directly with PopZ and subsequently recruits ClpX, substrates, and other members of a protease complex to the cell pole. We also find that co-recruitment of protease components and substrates to polar microdomains enhances their coordinated activity. This study connects phosphosignaling with polar assembly and the activity of a protease that triggers cell cycle progression and cell differentiation.

## Introduction

Many bacteria divide asymmetrically, generating two distinct cell types that can have vast differences in gene expression, morphology, and other behaviors^1–3^. This basic form of multicellularity was very likely to have arisen more than a billion of years before the emergence of eukaryotic life^4^. In rod-shaped bacteria, asymmetry is generally achieved through cell polarization, where distinct sets of regulatory proteins are directed to opposite cell poles, and subsequent cell division generates daughter cells that inherit different sets of regulatory factors and therefore different cell fates^5,6^. These systems depend on mechanisms that distinguish one pole from the other, and in many cases the associated molecular processes that drive this organization are not clear.

In Alphaproteobacteria, one of the key polarization factors is the polar organizing protein PopZ, which is required for the polar localization of many cell fate regulators^7^. *In vitro*, PopZ self-assemblies into oligomeric forms that undergo higher-order assembly into fibrils and larger structures^8,9^. Under some conditions, these assemblages behave as liquid-liquid phase separated protein condensates and have the ability to interact with client proteins ^10,11^. PopZ exhibits similar assembly and recruitment activities when expressed in *E. coli*, where it accumulates as a single cytoplasmic focus ^12–14^. In its natural context PopZ and its associated proteins form three-dimensional structures that abut the membrane at cell poles^15^. *C. crescentus* PopZ is localized to both poles through most of the cell cycle, yet several of the regulatory proteins that depend on it for polar localization exhibit transient, unipolar localization^16–18^.

The N-terminal domain of PopZ is an interaction hub that binds to at least eleven different client proteins and recruits them from the cytoplasm to polar microdomains ^12,19^. Including indirect interactions, PopZ serves as localization determinant for large interconnected protein networks^12,16–18^. For example, the protease ClpP and its partner ClpX are indirectly recruited to polar microdomains through interaction with the regulatory factor CpdR, which binds directly to PopZ^12,17^. CpdR is a cell cycle-regulated adaptor protein that delivers substrate proteins to ClpXP for timely degradation^20^. Some ClpXP substrates require one or more additional adaptor proteins, namely RcdA and PopA, and these are recruited to cell poles through interaction with PopZ at the same stage of the cell cycle as CpdR, which is known as the swarmer to stalked cell transition^21^. Thus, ClpP, ClpX, up to three adaptors, and the substrate proteins themselves^22,23^ are all co-recruited by PopZ into polar microdomains, where they become concentrated relative to bulk cytoplasm.

The question of how co-localization and concentration in PopZ microdomains affects the recruited proteins’ activities has not been conclusively resolved. Owing to the limited size of PopZ structures and the relative abundance of available clients, many PopZ-associated proteins are only partially localized in polar microdomains^24,25^. This raises the question of whether the levels of concentration and co-localization at cell poles are large enough to influence cell physiology. Surprisingly, one study found that three different pole-localized ClpXP substrates were degraded more quickly in a Δ*popZ* strain than in wildtype^26^, a phenotype that could occur if polar microdomains have an inhibitory effect on the activity of protease complexes. However, this is not easily reconciled with the observation that both ClpXP-mediated proteolysis and the polar accumulation of ClpXP complexes occur at the same stage of the cell cycle.

Two other unresolved questions concern the regulation of transient localization to cell poles and the mechanisms by which one PopZ microdomain is differentiated from another. These questions are exemplified by the members of the ClpXP proteolysis complex, which are localized to only one of the two poles, and only during the swarmer to stalked cell transition. Notably, both the CpdR and RcdA adaptor proteins are direct binding partners of PopZ^12,19^, which suggests that their interaction with the polar hub is subject to some form of regulation.

CpdR is a two-component response regulator whose phosphorylation state is regulated by the histidine kinase CckA, via an intermediary phosphotransfer protein ChpT. CckA exhibits kinase activity through the majority of the cell cycle, when CpdR and ClpXP protease complexes are neither pole-localized nor active in degrading substrates^17,20,27^. CckA works as a phosphatase during the swarmer to stalked cell transition, when CpdR is localized to one pole and ClpXP is active. CckA binds directly to PopZ and is localized to both cell poles through most of the cell cycle, where its activity is controlled through the influence of asymmetrically localized upstream regulators^28^.

Taking these observations together, we hypothesize that CckA kinase/phosphatase activity is a switch that regulates PopZ’s interaction with CpdR and the consequent recruitment of other members of ClpXP proteolysis complexes. In this work, we tested that hypothesis by modulating CpdR phosphorylation levels in *C. crescentus*, in a reconstituted *E. coli* system, and *in vitro*. We also asked if the concentration of ClpXP complex members at the cell pole could influence the overall rate of proteolysis for the entire cell, using a combination of cell imaging and computational modeling. The results explain how CpdR phosphorylation and de-phosphorylation influences polar asymmetry and cell differentiation during the *C. crescentus* cell cycle.

## Results

### CpdR-YFP localization is correlated with its phosphorylation state during the cell cycle

In wildtype *C. crescentus*, polar localization of CpdR-YFP temporally coincides with the time that it is dephospohrylatred by CckA-ChpT phosphotransfer during the swarmer-to-stalked transition^17,27^(Fig. 1a). We used cell cycle synchronization to quantify the relationship between CpdR-YFP localization and phosphorylation. After isolating cells at the swarmer cell (G0) stage, we counted the percentage of cells exhibiting polar CpdR-YFP localization and quantified the relative levels of phosphorylated versus unphosphorylated CpdR-YFP over the course of the cell cycle (Fig. 1b). The highest and lowest degrees of polar localization temporally corresponded with the highest and lowest fractions of unphosprylated CpdR-YFP, respectively. After cell division, CpdR-YFP remained diffuse in swarmer progeny but exhibited polar localization in stalked progeny, and the level of CpdR-YFP phosphorylation in this heterogenous population was balanced (Fig. 1c). We conclude that there is a strong positive correlation between CpdR-YFP de-phosphorylation and polar localization during the *C. crescentus* cell cycle.

**Fig. 1.**
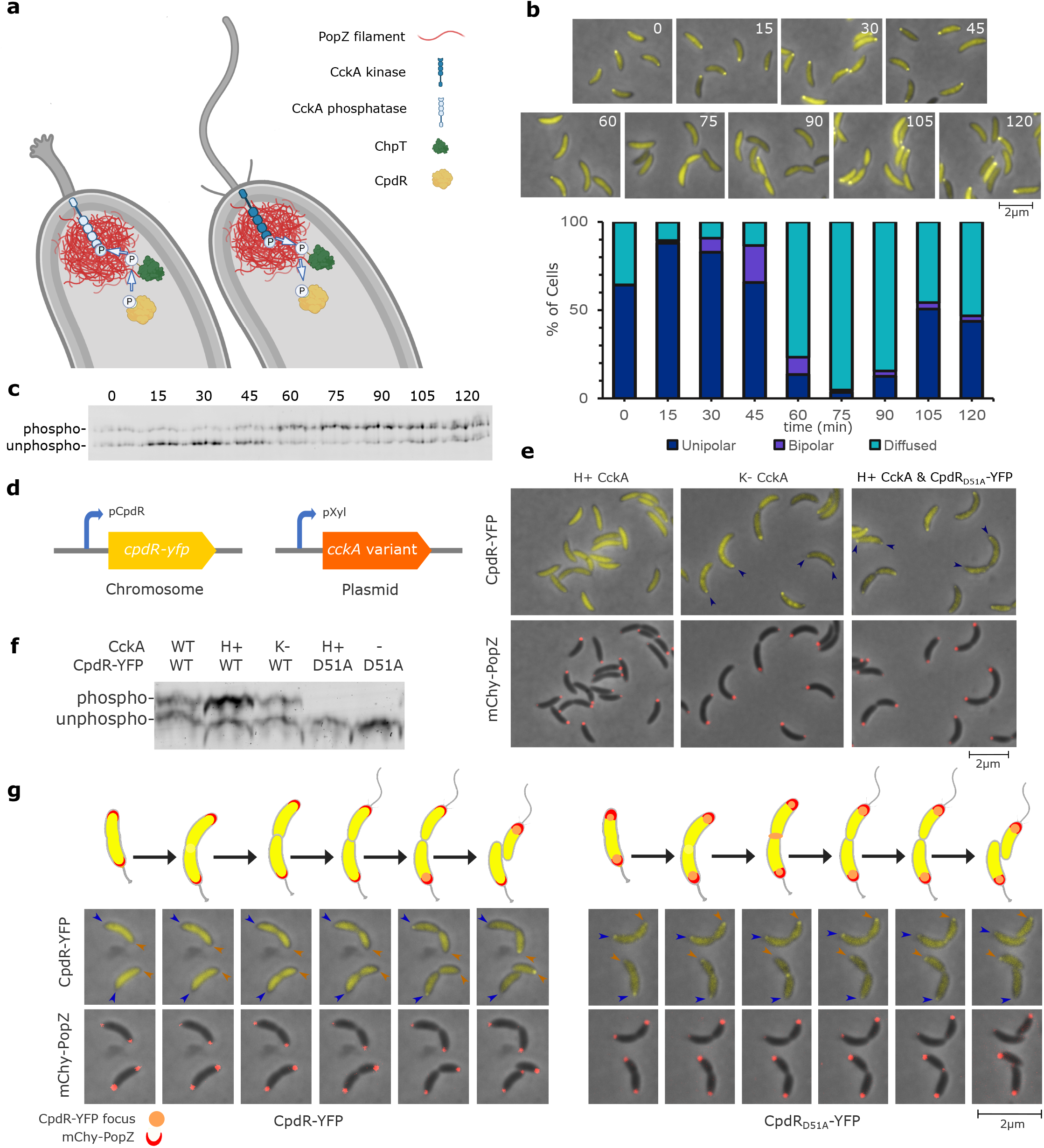
CpdR Phosphorylation State Influences its Co-localization with PopZ. **a**, CpdR phospho-signaling at stalked (left) and swarmer (right) cell poles, where CckA acts as a phosphatase or kinase, respectively. **b**, *C. crescentus* cells expressing CpdR-YFP were synchronized and, at indicated time points over a cell cycle time course, aliquots were removed for observation. The frequency of polar foci was plotted (n>300/time point). **c**, CpdR-YFP phosphorylation levels in lysates from b were observed by Phos-tag gel electrophoresis. **d**, Genetic modifications for controlling CpdR-YFP phosphorylation. Single-copy *cpdR-yfp* is expressed from the native promoter, multicopy *cckA* variants are expressed from P_xyl_ without xylose induction. **e**, Localization of CpdR-YFP or CpdR_D51A_-YFP and mChy-PopZ in different CckA signaling contexts. Arrowheads mark polar localization. **f**, CpdR-YFP phosphorylation levels in lysates from e, observed using Phos-tag gel electrophoresis. For CckA variants, WT=wildtype; H+=hyperactive kinase; K-=kinase deficient. To equalize loading of unstable non-phosphorylated CpdR-YFP, 3-fold or 5-fold more material was loaded in K- and D51A lanes, respectively. **g**, Cells expressing CpdR-YFP or CpdR_D51A_-YFP and mChy-PopZ were observed during cell division, using time-lapse microscopy at 15 minute intervals. Here, the fluorescence levels of individual panels were adjusted to aid visualization.

### CckA kinase influences CpdR-YFP phosphorylation and its co-localization with PopZ

We asked if we could influence CpdR-YFP phosphorylation and localization by expressing mutant variants of CckA with differing levels of kinase/phosphatase activity^29^. When we expressed a hyperactive-kinase (H+) form of CckA (G319E), the ratio of phosphorylated to unphosphorylated CpdR-YFP was relatively high, and the protein did not localize to cell poles (Fig. 1d-g). When we expressed a kinase-deficient (K-) form of CckA (H322A), the ratio was substantially lower, and cells exhibited robust polar foci. We also observed transient localization of CpdR-YFP at the division plane, which reflects CpdR’s role as an adaptor protein for FtsZ proteolysis^30^. To determine if polar localization is indirectly controlled by CckA signaling, we expressed a mutant form of CpdR-YFP (CpdR_D51A_-YFP) that cannot be phosphorylated^17^. Whether or not the hyperactive-kinase form of CckA was expressed in this context, CpdR_D51A_-YFP exhibited polar localization, and moreover, was usually localized to both of the cell poles in cells that also had a bi-polar distribution of PopZ (Fig. 1e, 1g). These results show that the kinase/phosphatase activity of CckA is closely correlated with the phosphorylation state of CpdR and its localization at *C. crescentus* cell poles.

### CpdR Phosphorylation affects interaction with PopZ in *E. coli*

We reconstituted CpdR phosphorylation and studied polar localization in *E. coli* to determine whether CpdR’s phosphorylation state affects its interaction with the polar organizing protein PopZ. To do this, we co-expressed CpdR-GFP and mChy-PopZ together with the phosphotransfer protein ChpT and either a wildtype, kinase-deficient, or hyperactive kinase variants of CckA (Fig. 2a). Experiments were performed in a Δ*clpXP* mutant background^31^ to eliminate the possibility of interference from interactions with endogenous protease^20^. We observed limited co-localization between CpdR-GFP and mChy-PopZ foci in cells expressing a wildtype variant of CckA, and no co-localization when the hyperactive kinase variant was expressed (Fig. 2b, Supplementary Fig. 1a). Conversely, strains expressing kinase-deficient CckA exhibited increased co-localization, and we found similarly elevated levels of co-localization when we expressed non-phosphorylatable CpdR_D51A_-GFP with hyperactive kinase CckA. We used Phos-tag gel electrophoresis to assess the relative levels of CpdR phosphorylation in these strains (Fig. 2c) and found that the ratio of unphosphorylated to phosphorylated CpdR is positively correlated with the degree of co-localization with mChy-PopZ foci. An N-terminal truncation mutant of PopZ failed to recruit CpdR-GFP into polar foci (Fig. 2a), indicating that the hub domain of PopZ is responsible for the interaction^19^.

**Fig. 2.**
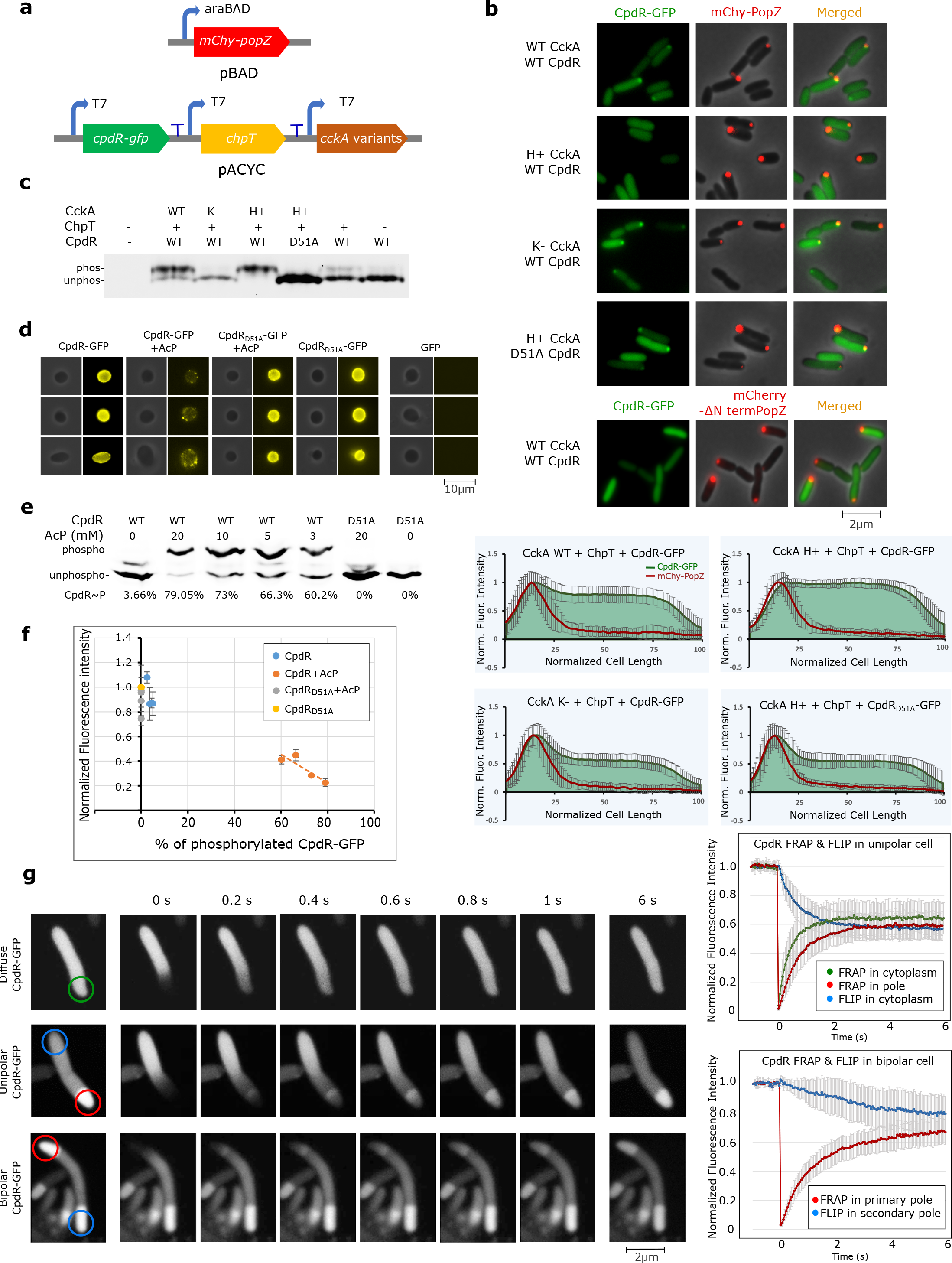
CpdR Phosphorylation State regulates CpdR-PopZ interactions. **a**, Reconstitution of CpdR phosphorylation and PopZ interaction in *E. coli*. **b**, mChy-PopZ and CpdR-GFP were observed by microscopy and normalized fluorescence intensities were plotted against cell length (graphs). For CckA variants, WT=wildtype; H+=hyperactive kinase; K-=kinase deficient. **c**, CpdR-GFP phosphorylation levels in lysates from c, observed using Phos-tag gel electrophoresis. **d**, Images of CpdR-GFP interactions with PopZ condensates, with phase contrast and YFP fluorescence channels shown. GFP-only is included as a negative control. +AcP: after incubation with acetyl phosphate. **e**, CpdR-GFP phosphorylation levels in samples from d, observed using Phos-tag gel electrophoresis. Data shows composite images from two different gels. **f**, Fluorescence values from d were normalized to the maximum signal intensity and plotted against percentage of CpdR-YFP∼P in 2e. **g**, FRAP and FLIP assay for CpdR-GFP in *E. coli* cells expressing PopZ. Recovery and loss of fluorescence were plotted against time in seconds.

### CpdR Phosphorylation affects interaction with PopZ *in vitro*

We use an *in vitro* assay to determine whether CpdR phosphorylation directly affects PopZ interaction. First, we produced macromolecular condensates of purified PopZ^10^ that specifically concentrated CpdR-GFP from solution. However, GFP alone was not recruited by these condensates (Fig. 2d). Next, we modulated CpdR-GFP’s phosphorylation state by pre-incubating it with varying concentrations of acetyl phosphate (AcP) (Fig. 2e). The degree of CpdR phosphorylation was negatively correlated with its partitioning in PopZ condensates (Fig. 2f). We controlled the presence of acetyl phosphate by applying it to non-phosphorylatable CpdR_D51A_-GFP and found that this did not affect protein mobility in Phos-tag gel electrophoresis or block partitioning within PopZ condensates. Taken together with the *E. coli* reconstitution experiments, these results suggest that CpdR phosphorylation inhibits physical interaction with PopZ.

### CpdR-PopZ interaction is highly dynamic

A large fraction of CpdR is diffuse in cell body, even when it almost entirely dephosphorylated (Fig. 1b, 2b). We hypothesized that this is reflective of a weak and therefore highly transient interaction with PopZ and tested this in FRAP experiments on *E. coli* cells co-expressing CpdR-GFP and mChy-PopZ (Fig. 2g). We found that PopZ-associated CpdR-GFP was rapidly replenished from the cytoplasmic pool after photobleaching, with a half-time of 0.61 seconds. This was slightly slower than the rate of recovery of CpdR-GFP diffusing through normal cytoplasm, at 0.36 seconds (Fig. 2g). During recovery, a wavefront of CpdR-GFP advanced through PopZ foci (Supplementary Video 1), which could occur if CpdR-GFP molecules are transiently held by interactions with PopZ. To obtain information on the off-rate, we created *E. coli* cells with mChy-PopZ foci at both cell poles. Bleaching CpdR-GFP at one pole resulted in rapid fluorescence recovery, with concomitant rapid fluorescence loss at the opposite pole (Fig. 2g). By contrast, the recovery rate of mChy-PopZ had a half-life of over 3 minutes (Supplementary Fig. 1b), indicating that associations among the scaffold molecules are more stable.

### CpdR phosphorylation influences the localization of RcdA and ClpX

We hypothesized that we could control the localization of ClpXP complexes (Fig. 3a) by modifying the phosphorylation state of CpdR. To test this, we created strains in which either the ClpX-associated adaptor RcdA ^18^ or ClpX itself were tagged with GFP and expressed from their endogenous promoters, and in which the native copy of *popZ* had been exchanged with a functional mChy-tagged version^7^ (Fig. 3b). In an otherwise wildtype genetic background, RcdA-GFP and ClpX-GFP both exhibited the expected pattern of transient co-localization with polar mChy-PopZ during the swarmer-to-slaked cell transition^18,30^ (Fig. 3c). In a Δ*cpdR* background, RcdA-GFP was diffuse, which adds to an earlier report that CpdR is required for ClpX localization^17^ (Fig. 3c).

**Fig. 3.**
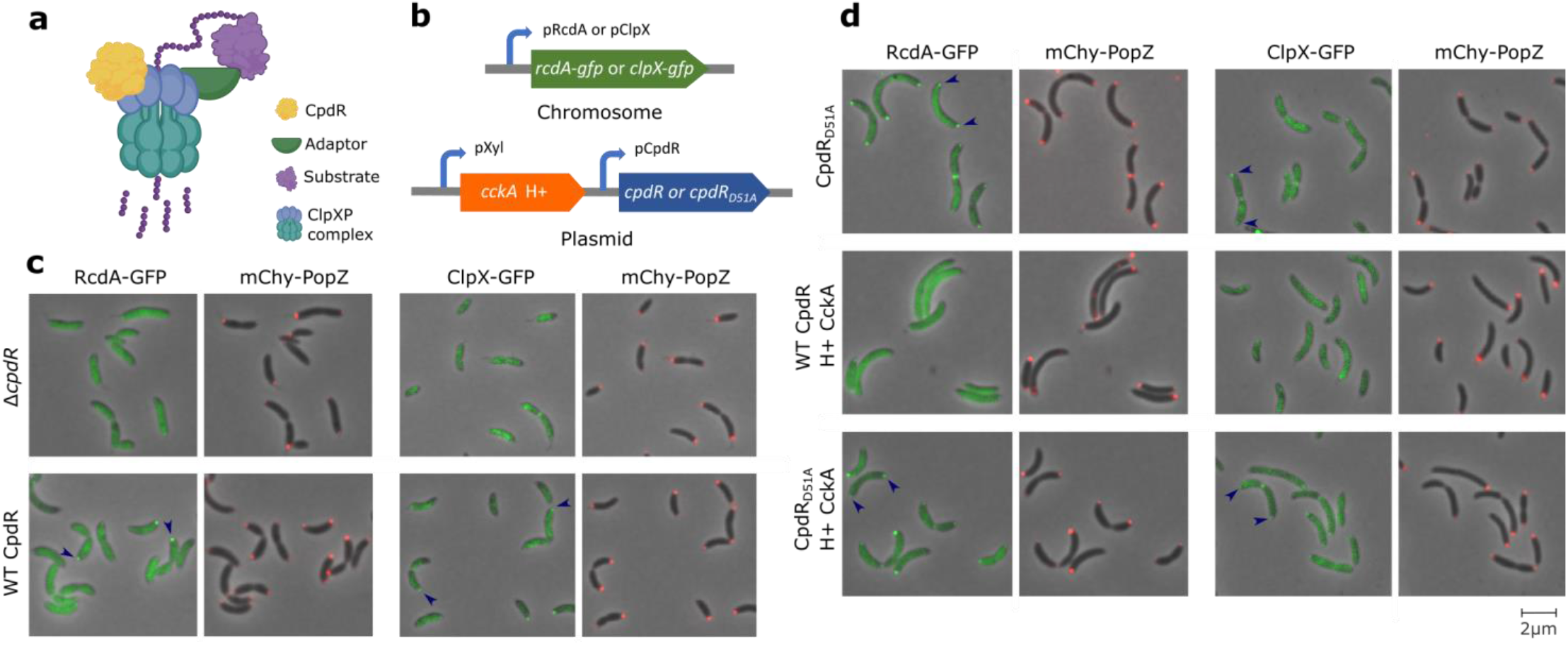
CpdR Phosphorylation State Drives the Localization of Protease Components. **a**, Diagram of CpdR-mediated substrate degradation by ClpXP. Depending on the substrate, an additional adaptor (green) could be RcdA and/or PopA. **b**, Genetic modifications for observing RcdA -GFP or ClpXP -GFP in different CpdR phosphorylation contexts, built in a Δ*cpdR* ; *popZ::mChy-popZ C. crescentus* strain background. **c-d**, Localization of RcdA-GFP or ClpX-GFP and mChy-PopZ in different CpdR phosphorylation contexts. Arrowheads mark polar localization.

To assess protein localization in cells where CpdR is always dephosphorylated, we replaced endogenous *cpdR* with the *cpdR*_*D15A*_ variant. Strikingly, RcdA-GFP was localized to both poles in cells that also had bi-polar mChy-PopZ foci, and ClpX-GFP showed a similar, though less intensely localized pattern (Fig. 3d). To assess protein localization in cells where CpdR is always phosphorylated, we expressed the hyperactive variant of CckA. In these cells, RcdA-GFP and ClpX-GFP were diffuse. We also expressed hyperactive CckA in the *cpdR*_*D51A*_ background. Here, RcdA-GFP and ClpX-GFP exhibited bi-polar localization (Fig. 3d), indicating that the phosphorylation state of CpdR is the controlling factor. The differences in these strains’ localization patterns suggest that CpdR’s phosphorylation state is normally under strict temporal and spatial control, limiting ClpXP complexes to one pole at the swarmer-to-slaked cell transition. Since RcdA-GFP requires ClpX for polar localization^18^, CpdR may influence its localization indirectly. We propose the following order of protein recruitment at *C. crescentus* cell poles: PopZ -> dephosphorylated CpdR -> ClpX -> RcdA.

### Substrate Proteolysis coincides with Polar Localization

To assess the localization of proteolysis substrates, we expressed YFP-tagged versions of PdeA, TacA, and CtrA RD+15^32^ in wildtype, Δ*cpdR*, and Δ*popZ* genetic backgrounds. These three substrates were expressed from chromosomally integrated plasmids that drove gene expression from xylose-inducible promoters, and were chosen as representatives of CpdR-dependent, CpdR/RcdA-dependent, and CpdR/RcdA/PopA-dependent classes, respectively. Their localization patterns are consistent with an earlier report on PdeA localization^33^, and add to the literature by showing that YFP-TacA localization depends on RcdA (Supplementary Fig. 2b), and that all three classes of substrates exhibit transient, unipolar localization in wildtype cells and diffuse localization in both Δ*popZ* and Δ*cpdR* backgrounds (Fig. 4a).

**Fig. 4.**
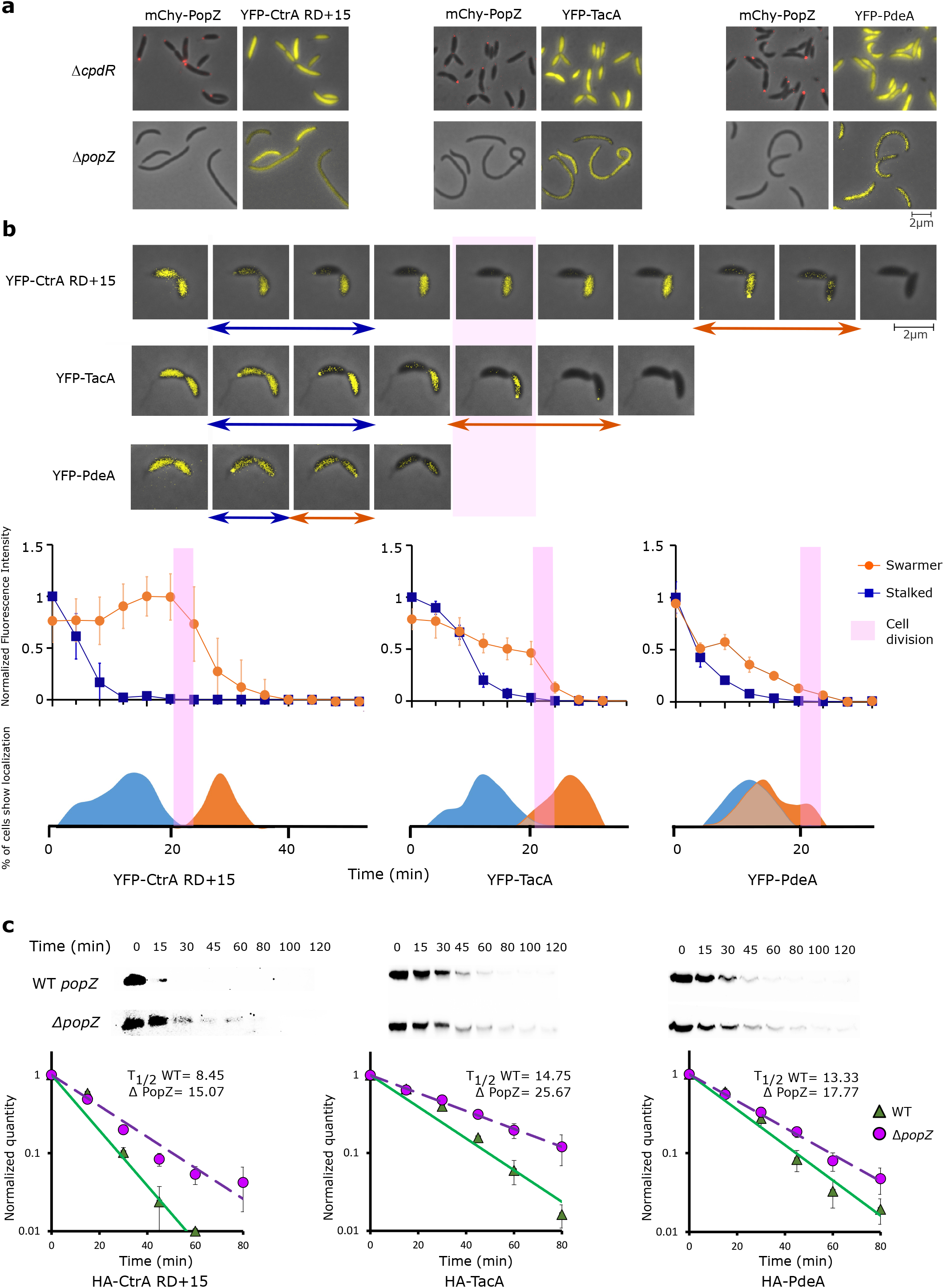
Polar localization of CpdR substrates influences their rate of degradation. **a**, Localization of YFP-tagged substrates in Δ*cpdR*; *popZ::mChy-popZ* and Δ*popZ C. crescentus* strain backgrounds. **b**, Time-lapse images of YFP-tagged substrate localization in a WT *C. crescentus* background, at 4 minute intervals. Blue arrows mark frames with foci in stalked cell, orange arrows mark frames with foci in swarmer cell. Pink bar idicates the time of cell separation. After accounting for photobleaching and temporally alignging the cells with respect to the time of cell separation (n=20), average fluorescence intensities for stalked and swarmer cell bodies, normalized to maximum fluorescence intensity, were plotted against time (line graphs). The percentage of cell bodies ehxhibiting polar localization, normalized to the highest percentage, were plotted on the same time axis (solid curves). **c**, Degradation of HA-tagged proteolysis substrates following inducer wash-out, observed by western blotting with α-HA antibody. Average band intensities from three separate experiments were plotted against time (graphs, bar= standard deviation).

We used time-lapse fluorescence microscopy to observe the spatial-temporal relationships between localization and proteolysis for the three substrate proteins. In stalked cell progeny, all three proteolysis substrates were cleared from the cell between before cell separation, and the highest frequency of polar localization was observed during this time period (Fig. 4b). Since cell separation lags the separation of progeny cells’ cytoplasm via inner membrane fusion by several minutes^34^, it is likely that proteolysis and polar localization occurred soon after compartmentalization. In swarmer cell progeny, YFP-PdeA was often cleared concomitantly or within four minutes of its clearance from stalked cells, which coincided with the highest frequency of polar localization. The majority of YFP-TacA and YFP-CtrA RD+15 were cleared approximately 12 minutes later, when their peak frequency in polar localization occurred (Fig. 4b). From these observations, we conclude that the polar localization of all three proteolysis substrates is closely correlated with their time of proteolysis.

### PopZ facilitates rapid substrate degradation

To better understand the functional relationship between substrate protein degradation and polar localization, we performed two different types of experiments to assess the degradation rates in wildtype versus Δ*popZ* cells. In the first type of experiment, we blocked new protein synthesis by treating cells with chloramphenicol. Over subsequent timepoints, a subpopulation of cells in wildtype cultures retained large quantities of substrates for more than 60 minutes, but this was not observed in Δ*popZ* cultures (Supplementary Fig. 3a). We propose that chloramphenicol treatment interferes with the measurement of degradation rates by preventing cell cycle progression in swarmer cells, locking them into a stage of high substrate stability. Δ*popZ* strains, whose cell division is often uncoupled from the cell cycle ^7^, do not appear to produce significant numbers of this cell type.

In the second type of experiment, we used inducer wash-out to block new substrate protein synthesis, allowing general protein synthesis and the cell cycle to continue (Fig. 4c). Under these conditions, ClpXP substrates were degraded up to twice the rate in wildtype cells compared to Δ*popZ*. We therefore conclude that ClpXP substrates are degraded more rapidly when PopZ microdomains are present. Taking this together with the observation that the protease, adaptors, and substrates are concentrated in PopZ microdomains during proteolysis (Fig. 1g, 3d, 4b), we propose that one of the functions of PopZ microdomains is to enhance the assembly of ClpXP-adaptor-substrate complexes.

### A conceptual model for enhanced assembly of proteolysis complexes in PopZ microdomains

PopZ microdomains occupy approximately 0.5% of the cytoplasm ^15^, and only a fraction of proteolysis substrates and protease components are found in this relatively small compartment. A question is whether PopZ microdomains could enhance proteolysis complex assembly and activity on a scale that is sufficient to affect the overall rate of degradation for the entire cell. Using the biochemical simulation program Smoldyn^35^, we developed a computational model to determine whether the physiological characteristics of the system, in terms of compartment sizes, protein localization, diffusion rates, rate of proteolysis, and number of substrate molecules, are compatible with the idea that PopZ microdomains enhance proteolysis.

Because a complete model of adaptor mediated ClpXP proteolysis would involve a large number of unknown parameters, we simplified the system by modeling an interaction between two reactants, A and B, which were eliminated after colliding. Using single molecule tracking information on CtrA and other pole-localized proteins in *C. crescentus* as a guide^25^, A and B were allowed to exhibit Brownian motion in PopZ microdomains, but with a diffusion coefficient that was 10 to 100-fold lower than bulk cytoplasm. This had the effect of increasing local protein concentration, mimicking the effect of weak interactions with PopZ.

In our simulations of cells with physiologically sized polar microdomains, 50% of reactants were degraded within 1.5 minutes, compared to more than 5 min in cells that lacked a polar microdomain (Fig. 5a). Increasing the number of substrate molecules in polar compartments, by either increasing compartment volume or lowering the rate of diffusion at the cell pole, had the effect of increasing reaction rate (Fig. 5a, 5b; Supplementary Video 2, 3). Gains in reaction efficiency were subject to rational limits on microdomain volume and particle behavior (Supplementary Fig. 4). For example, at extremely low diffusion coefficients, particles became “stuck” in polar microdomains and although highly concentrated, collided less often. In conclusion, the simulations provide support for the idea that protein-protein interactions are enhanced when they are concentrated within PopZ microdomains, and that this can occur on a size and time scale that is relevant to proteolysis *in vivo*.

**Fig. 5.**
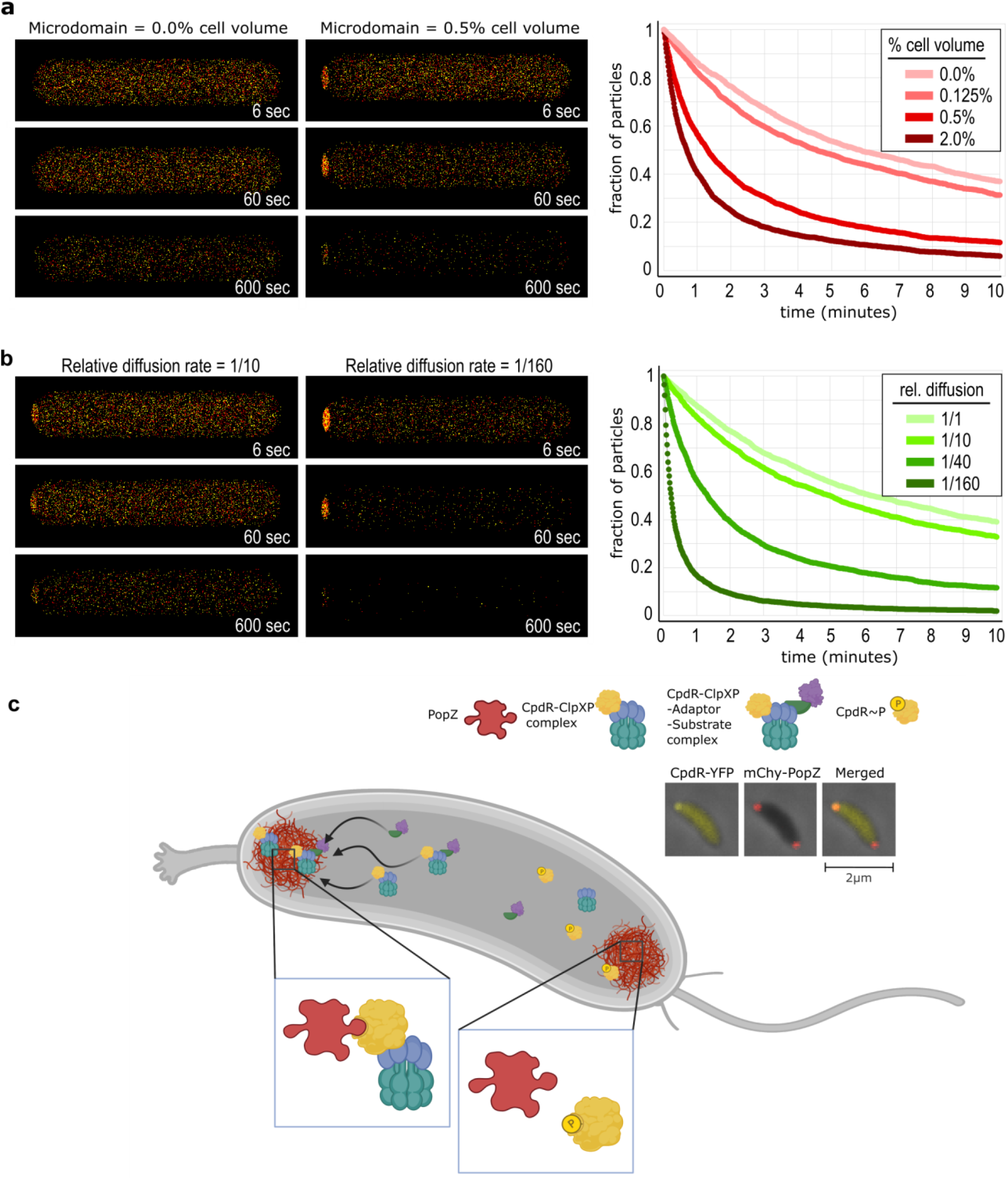
Conceptual models of substrate proteolysis in membraneless polar microdomains. **a-b**, Three-dimensional reaction-diffusion simulations with two types of particles, colored red and yellow, that disappear after colliding. Parameters for cell size, particle diffusion rate in bulk cytoplasm, and the number of particles are based on physoligical values for *C. crescentus*. **a**, Cells with different sizes of membraneless polar microdomains, where particles become concentrated because they diffuse at 1/40^th^ the rate in bulk cytoplasm. PopZ microdomains occupy 0.5% of cell volume *in C. crescentus*. **b**, Cells with different concentrations of particles in polar microdomians. Slower diffusion rates in polar microdomains result in higher concentration. **c**, Asymmetric localizatin of CpdR and associated ClpXP complexes as a consequence of asymmetric CckA signaling activity. Inset panels show fluorescence images of a *C. crescentus* stalked cell, where mChy-PopZ is localized to both poles and CpdR-YFP is localized to only the stalked pole.

## Discussion

In this work, we show that CpdR’s interaction with pole-localized PopZ is closely correlated with its phosphorylation state (Fig. 1 and 2), that changing CpdR phosphorylation level through CckA signaling or the expression of the CpdR_D51A_ variant is sufficient to alter the localization of CpdR and associated ClpXP complexes between diffuse and bi-polar (Fig. 3d), and that the polar accumulation of ClpXP substrates is positively correlated with proteolysis (Fig. 4, 5, Supplementary Fig. 4). Together, these results suggest that CpdR-mediated recruitment of ClpXP to polar PopZ subdomains stimulates proteolytic activity (Fig. 5b).

This is the first study to show that phosphorylation directly regulates a client protein’s interaction with the PopZ hub. An earlier study showed that a chromosome segregation protein, ParA, loses affinity to PopZ after hydrolyzing ATP^36^. Other studies have found that a ClpX adaptor, PopA, interacts with PopZ when it is bound to cyclic di-GMP^37,38^. CpdR has a more basal position than PopA in the adaptor hierarchy that determines ClpXP substrate specificity, meaning that it targets a substantially larger number of substrates, including those that are targeted by PopA^23^. A third adaptor, RcdA, interacts directly with PopZ in *E. coli* co-expression experiments but requires de-phosphorylated CpdR (Fig. 3c) or c-di-GMP bound PopA^39^ for polar localization in *C. crescentus*, where protein expression levels are lower. We hypothesize that RcdA itself has a relatively low affinity to PopZ that is increased by associating with other PopZ-interacting proteins, forming multiprotein complexes with high avidity. All three ClpX adaptors’ interactions with PopZ are post-translationally regulated, suggesting that this is a key leverage point in the regulation of proteolysis and cell cycle progression.

The mechanisms that control whether CckA works as a kinase or phosphatase are driving forces behind the localization of CpdR and associated members of ClpXP protease complexes. In stalked cells, where PopZ exhibits bipolar localization and CpdR/ClpXP are exclusively at the stalked pole (Fig. 1g, 3c), the differential activity of CckA at opposite poles^29^ may be sufficient to support this aspect of polar asymmetry. However, CckA control may not be sufficient to explain all aspects of protease localization and activity. Whereas three different substrates were localized and degraded simultaneously in stalked cell progeny, PdeA was localized and degraded several minutes earlier than TacA and CtrA in swarmer cell progeny (Fig. 4b). We hypothesize that the timing differences are related to the expression of additional regulatory factors during pole remodeling at the swarmer to stalk transition. CtrA proteolysis requires a polar di-guanylate cyclase to activate PopA^33^, and TacA proteolysis may require the phosphatase stimulating activity of SpmY, which is also recruited to the transitioning pole^40^.

The term “polar localization” is used here and in most other reports to communicate the idea that a protein is visibly concentrated at the cell poles relative to the bulk cytoplasm. But this term is highly imprecise and potentially misleading, as a protein may exhibit “polar localization” when a small fraction of the total is at a pole. For example, since both cell poles occupy approximately 0.1% of the total cytoplasm volume, only 1% of a protein that is 10-fold concentrated at cell poles would be pole-localized. Indeed, fluorescence images of many different proteins in *C. crescentus*, including CpdR, substrates, and other members of the ClpXP complex, suggest low levels of actual polar localization. How could a system benefit from having such a localization pattern?

An answer to this question may lie in the fact that many pole-localized proteins receive regulatory cues at a cell pole and carry out their activity in bulk cytoplasm. CpdR is de-phosphorylated by CckA at the stalked cell pole, yet its targets are transcriptional regulators, chemotaxis regulators, and other proteins that work outside of PopZ microdomains. Thus, dynamic exchange between pole and cytoplasm is required for translating polar signaling into cytoplasmic activity. This is consistent with the CpdR-PopZ interaction dynamics we observed in *E. coli* (Fig. 2g) and the behavior of other polar regulators in *C. crescentus*^25,41^.

Further, low levels of polar localization coupled with highly dynamic exchange can have a substantial influence on entire populations of molecules on the whole-cell scale (Fig. 5). Applying this to adaptors, substrates, and ClpXP complexes, we propose that their concentration in PopZ microdomains increases the frequency of intermolecular collisions and therefore enhances multiprotein complex assembly and overall proteolysis rate. Faster ClpXP-dependent proteolysis could make the cell cycle more robust by sharpening phase changes at the swarmer-to-stalked transition.

## Methods

### Bacterial Plasmid and strain construction

All the bacterial plasmids were constructed by Gibson cloning method. Descriptions of all plasmids used in this study are given in Supplementary Table 1. Descriptions of all the bacterial strains used in this study are listed in Supplementary Table 2. Proteins were fluorescently labeled with mCherry (mChy), msfGFP or eYFP (see FPbase.org).

### *E. coli* vectors

Among the two multiple cloning sites of pACYC-duet (Invitrogen); from site 1 we used NcoI and NotI restriction sites for cloning *cpdR-gfp* and from site 2 we used NdeI and XhoI restriction sites for cloning *chpT*. For both CpdR-GFP and ChpT, as start codon GTG was used to ensure moderate protein expression. In pCDF-duet (Invitrogen) plasmid, *cckA* variants were cloned at site 1 bearing NcoI and NotI restriction site. These *cckA* variants were PCR amplified, including the T7 promoter, and cloned at NheI restriction site upstream of p15A ori in pACYC-duet. In a separate pACYC-duet plasmid, *cpdR-gfp-6xhis* was cloned at site 1.

#### *C. crescentus* vectors

Plasmid pBXMCS2 was used for cloning *cckA* variants in between NdeI and EcoRI restriction sites. pMCS5 vector was used to clone *cpdR* or *cpdR-yfp* variants. pXMCS5 was used to clone *yfp-ctrA RD+15, yfp-tacA, yfp-pdeA, HA-ctrA RD+15, HA-tacA or HA-pdeA*. pVMCS6 was used to clone *rcdA-gfp* or *clpX-gfp*. NdeI and KpnI restriction sites were used in pMCS5, pXMCS5 and pVMCS6 vectors. All of these vectors were described by Thanbichler et al.^42^.

### Bacterial Cell culture

*E. coli* cells were grown at 30 °C overnight in Luria-Bertani (LB) liquid media or at 37 °C in LB agar plate supplemented with 1.5% agar. Liquid cultures were grown on rotor if not mentioned otherwise and all the growth of bacterial culture was measured by absorption OD_600_. For *E. coli* strains, antibiotics were used at following concentrations: 50 μg ml^-1^ ampicillin, 20 μg ml^-1^ chloramphenicol, 50 μg ml^-1^ spectinomycin, 12 μg ml^-1^ oxytetracycline, 30 μg ml^-1^ streptomycin, 30 μg ml^-1^ kanamycin.

*C. crescentus* was grown at 28 °C in PYE liquid media or PYE agar plates supplemented with 1.5% agar. For antibiotic-resistant strains, antibiotics were used at the following concentrations: 2 μg ml^-1^ chloramphenicol, 25 μg ml^-1^ spectinomycin, 1 μg ml^-1^ oxytetracycline, 5 μg ml^-1^ streptomycin, 5 μg ml^-1^ kanamycin. *C. crescentus* plasmids were transformed via electroporation or genes with associated antibiotic markers were transduced using phage *ϕCr30*. Prior to analysis, stationary phase cells were diluted 50X in fresh PYE and grown until they reached an OD_600_ = 0.3. Xylose-inducible genes were induced by 0.2% final concentration of D-xylose unless mentioned otherwise. *C. crescentus* synchronies were performed according to Toro et al.^43^ but in PYE media at 28 °C.

### Wide-field Microscopy

Cells were immobilized on a 1% agarose gel pad and viewed with a Zeiss Axio Imager Z2 epifluorescence microscope equipped with a Hamamatsu Orca-Flash 4.0 sCMOS camera and a Plan-Apochromat 100x/1.46 Oil Ph3 objective. Zen Blue software was used for image capture and quantification. For fluorescence imaging, mChy was observed by excitation at 587 nm and emission at 610 nm, GFP by excitation at 488 nm and emission at 509 nm, and YFP by excitation at 508 nm and emission at 524 nm. Exposure times for imaging mChy, GFP and YFP tagged proteins were 500 ms, 500 ms and 1000 ms respectively. All microscopy was performed at 1000X magnification.

### Phos-tag gel assays

#### Lysis buffer composition

Buffer A: 0.5 M Tris HCl 0.4% SDS pH 6.8 1.25 ml, IgePal CA-630 100 μl, Glycerol 1.5 ml, H20 up to 9.5 ml Buffer B: 50 μl 2-mercaptoethanol, 10 μl EDTA free protease-phosphatase inhibitor, 20 μl lysozyme (10 mg ml^-1^), 0.2 μl Nuclease/Benzonase, 920 μl Buffer A Sample buffer 10X: 0.5 M Tris HCl 0.4% SDS pH 6.8 1.25 ml, Glycerol 5 ml, 10 mg Bromophenol Blue, H20 up to 10ml

#### Bacterial Cell Lysis Protocol

1ml cells of OD_600_ 0.6-1 were taken, centrifuged at 13000 rpm 1 min, and pellets were resuspended in.100 μl of freshly prepared Buffer B by mixing with a pipettor. After Incubating on a roller/rotor for 10 min at room temperature, the lysed samples were centrifuged at 13000 rpm for 5 min at 4 °C and the cleared supernatant was carefully collected in a new tube and kept on ice. 10 μl of 10X sample buffer was added and 30 μl sample was loaded into each well of a prepared Phos-tag gel in a cold room at 4 °C.

#### Phos-tag gel composition

10-12% gels were prepared according to manufacturer’s instructions (Wako) with the exception of adding 40 μM MnCl_2_ and 15 μM Phos-tag reagent as final concentration. The gel and the running buffer were kept at 4 °C for 30 min prior to loading the sample. The samples were run at 100 volts until the dye front eluted from the gel. As the desired proteins were tagged with GFP or YFP, the gels were imaged in a gel doc (Biorad Chemidoc™ MP) at 488 nm wavelength.

### *E. coli* co-expression assay and fluorescence quantitation

Stationary phase *E. coli* cells bearing pACYC-duet and pBAD plasmids were diluted 100X in fresh LB and grown on rotor at 30 °C until they reach mid exponential phase with an OD_600_ = 0.3 (strain GB#1971-1977, Supplementary Table 1). Cells were induced with 0.2% arabinose for 3 hours and with 10 μM IPTG for 2 hours to express mChy-PopZ and CpdR-GFP+ChpT+CckA, respectively. Microscopic imaging was performed on 3 different replicative samples and 60 representative cells from each cell type (20 from each replicate) were chosen for fluorescence quantification. Cells with zero mChy-PopZ foci or foci at both poles were excluded from the analysis. As described by Nordyke et al.^19^, background-subtracted pixel intensities for each channel were measured along a straight line drawn lengthwise through mid-cell. Cubic spline interpolation was used to generate fluorescence intensity values for 100 equally spaced points along each line, then all points were normalized to 1 prior to averaging.

### Protein Purification

PopZ, CpdR-GFP, CpdR_D51A_-GFP proteins were expressed using strains GB#169, GB#1969 and GB#1970.

*E. coli* cells were grown overnight up to the stationary phase, which was diluted 100X in 1L LB broth. Eppendorf innova S44i shaker incubator was used to shake at 200 rpm and culture was grown back to the OD_600_=0.6. Then the culture was induced using final concentration of 1 mM IPTG. Cells were grown for 6 hours and then harvested by centrifuging at 4000 rpm for 30 min in model centrifuge. Cells were resuspended in 20 ml HMK buffer (20 mM Hepes, 2 mM MgCl_2_, 100 mM KCl pH 7.5) and lysed using french pressure cell press (Sim-aminco) with 1000 psi pressure. Proteins were purified using HisPur cobalt resin (Thermo Scientific) according to the product protocol in batch purification method. Purity was checked using SDS-PAGE followed by Coomassie blue staining. Protein concentration was quantified by BCA protein assay kit (Thermo Scientific) according to the user manual. Purified proteins were preserved in 100 μl aliquots at -80 °C after flash freezing in liquid nitrogen.

## PopZ condensates and *in vitro* phosphorylation

*In vitro* phosphorylation was performed with the indicated concentration of Acetyl phosphate (AcP) in HMK buffer with 20 mM MgCl_2_ and at a final concentration of 100 μM CpdR-GFP. Samples were incubated at 37 °C for 6 hours. To form PopZ condensates, 20 mM MgCl_2_ was added to 100 μM purified PopZ for 10 min at room temperature. Next, 20 μM PopZ condensates were mixed with 5 μM CpdR-GFP or GFP, and 7μl of this mixture was transferred to a glass slide and covered with a coverslip. Imaging was performed immediately thereafter using phase contrast and the YFP fluorescence channel at 1000X magnification. 10 condensates of similar size from each type of mixture were chosen for fluorescence quantification using FIJI plugin MicrobeJ.

### ClpXP substrate degradation assay and Western blot

WT and *ΔpopZ C. crescentus* strains, expressing HA tagged CtrA RD+ 15 (GB#1989, GB#1992), TacA (GB#1990, GB#1993), PdeA (GB#1991, GB#1994) were induced at OD_600_=0.3 with D-xylose in 3 ml volume each for 2 hours. Cells were washed 3 times by centrifugation and resuspension in fresh PYE, with final adjustment to approximately 3 ml at the same OD_600._ Over the subsequent 2 hours of growth on a rotary shaker at 28 °C Temperature, 300 μl sample aliquots were harvested and centrifuged at 9000 rpm for 2 min. The cell pellets were resuspended in 30 μl SDS-PAGE sample buffer, flash frozen in liquid nitrogen, and preserved at -80 °C. Whole sample were loaded in a 10% SDS-PAGE gel and resolved at 100 volts. After transferring to a PVDF membrane (0.45 μm pore size for Ha-TacA and HA-PdeA samples, 0.22 μm pore size for HA-CtrA RD+15 samples), Western blots were developed using an anti-HA mouse monoclonal primary antibody (Invitrogen).

Similar analyses were performed on strains expressing YFP tagged CtrA RD+ 15 (GB#1983, GB#1986), TacA (GB#1984, GB#1987), PdeA (GB#1985, GB#1988). After washing away the inducer, chloramphenicol was added at 30 μg ml^-1^ where appropriate and incubated for 5 min at 28 °C. From that point forward, fluorescence imaging was performed at 4 min time intervals and sample collection for western blot was performed as described above. Fluorescence quantification was performed using FIJI plugin MicrobeJ. We normalized the fluorescence values in *C. crescentus* time-lapse experiments to account for photobleaching. To obtain bacterial cells with constant levels of YFP, we induced YFP expression in *E. coli* cells from the pBAD promoter to a YFP fluorescence intensity level that was no more than 2-fold different than the *C. crescentus* cells we wished to analyze. Next, we imaged the *E. coli* cells under the same time-lapse conditions that were used to observe *C. crescentus*. The total fluorescence of 10 *E. coli* cells were observed over 10 consecutive time points. Following background subtraction, the total cell fluorescence signal from time point T_1_ was divided by the total cell fluorescence signal from T_n_ to obtain a YFP photobleaching coefficient for T_n_ time point. To obtain fluorescence values for *C. crescentus* observations, YFP photobleaching coefficients corresponding to the appropriate time intervals were multiplied by the whole cell *C. crescentus* fluorescence values.

### Photobleaching assay

Olympus IX-81 inverted confocal microscope equipped with a Yokogawa spinning disk (CSU X1) and a sCMOS camera (Orca-FLASH 4.0; Hamamatsu) was used for confocal microscopy. Excitation wavelengths were controlled using an acousto-optical tunable filter (ILE4; Spectral Applied Research) and 405 FRAP laser was used with 100 mW nominal power. MetaMorph 7.7 software (MetaMorph Inc.) was used for image acquisition. z-Stack images were acquired for selection of appropriate cells using a 100×, 1.40 N.A. oil objective; FRAP time-lapse images were acquired using a 60×, 1.35 N.A. oil objective at 50 ms time intervals. To create bipolar mChy-PopZ containing cells, *E. coli* liquid culture was supplemented with 30 μg ml^-1^ cephalexin along with 0.1 mM IPTG and 0.2% arabinose at OD_600_ = 0.3. Cells were incubated for 4 hours at 30 °C. To plot Fluorescence Recovery After Photobleaching (FRAP) and Fluorescence Loss in Photobleaching (FLIP), time-lapse images of 10 different cells from each type were acquired. Fluorescence intensity of the photobleached area, unbleached area and background area were quantified using FIJI software for all the time points. Normalization of fluorescence was performed by subtracting the background signal and multiplying by the photobleaching coefficient. Photobleaching coefficients were obtained using a neighboring unbleached cell as a reference, by the method described in the previous section.

### Computational Simulation

Smoldyn files are provided in Supplementary Data 1 and a summary of the simulation parameters is provided in Supplementary Table 3. The following information was used to determine simulation parameters:

Cell dimensions, compartment sizes, and number of proteins: We chose 1 size unit in Smoldyn to equal 10nm. Basing our models on a measured cytoplasm diameter of, we created rod-shaped cells with length and volume that correspond to approximately 2.84 μm and 0.61 femtoliters, respectively, which are consistent with measurements of stalked cells^44^. PopZ compartment size has been measured by 3D super-resolution fluorescence imaging^45^. We chose a mid-range value of 3.4 × 106 nm^3^ for our standard cell models, which corresponds to approximately 0.56% of the total cytoplasmic volume. Information from global measurements of mRNA translation rates^24^ and fluorescence intensity measurements of FP-tagged proteins have been combined to provide the following estimates of protein abundance in *C. crescentus* stalked cells: PopZ 8,200; CtrA 25,402; CpdR 6,044; RcdA 2,583^46^. Using these numbers as a guide, we set the total number of A and B proteins in our model at 2,500 each.

Diffusion coefficients and time steps: The parameters for diffusion coefficients are based on single-particle tracking studies in *C. crescentus* cells^25^. ChpT and CtrA were found to move sub-diffusively in bulk cytoplasm, with diffusion coefficients of 1.8 μm^2^/sec, which translates to 18 units^2^/msec in our scaled simulations. These proteins continue to exhibit Brownian diffusion while co-localized with PopZ at cell poles, but at substantially slower rates of 0.1 μm^2^/sec and 0.01 μm^2^/sec, respectively (corresponding to 1 units^2^/msec and 0.1 units^2^/msec). We used these values as approximations for the diffusion of molecules A and B in our standard cell models, setting their diffusion coefficient at 20 units^2^/msec in bulk cytoplasm and 0.5 units^2^/msec in polar subdomains. We set the time step to 0.1 msec, which corresponds to a resolution of approximately 20nm^35^.

Reaction probabilities and binding radii: Smoldyn represents proteins as point-like particles, and updates each particle’s position over iterative time steps, assigning a random direction and a distance calculated from its diffusion coefficient. The rate of a second order reaction, as occurs for A + B -> 0 in our simulation, is determined by particle proximities. When two reacting particles are positioned within a defined binding radius at the end of a time step, those particles react. Reactions rates are determined by the combined influences of binding radii, particle concentrations, and diffusion rates. We used an empirical process to arrive at a reasonable binding radius for A and B. We estimated that approximately 90% of proteolysis substrates are cleared from *C. crescentus* cells within 10 minutes of the swarmer to stalked cell transition (Fig. 4b). After entering a range of binding radii into a base model with the other parameters described in the preceding paragraphs, we found that a binding radius of 0.025 for A and B provided a degradation rate that is comparable to those experimental observations. This binding radius was used in all of the simulations shown in this work. Simulations were run on the University of Wyoming Beartooth High Performance Computing (HPC) environment.

## Acknowledgements

The authors would like to thank Steve Andrews for comments and advice. This work was supported by the National Institutes of Health under award numbers 2P20GM103432 and R01GM118792, and by the National Science Foundation under award number 1518171.

