## Supplementary Figures for "Phospho-signaling couples polar asymmetry and proteolysis within a membraneless microdomain in *C. crescentus*"

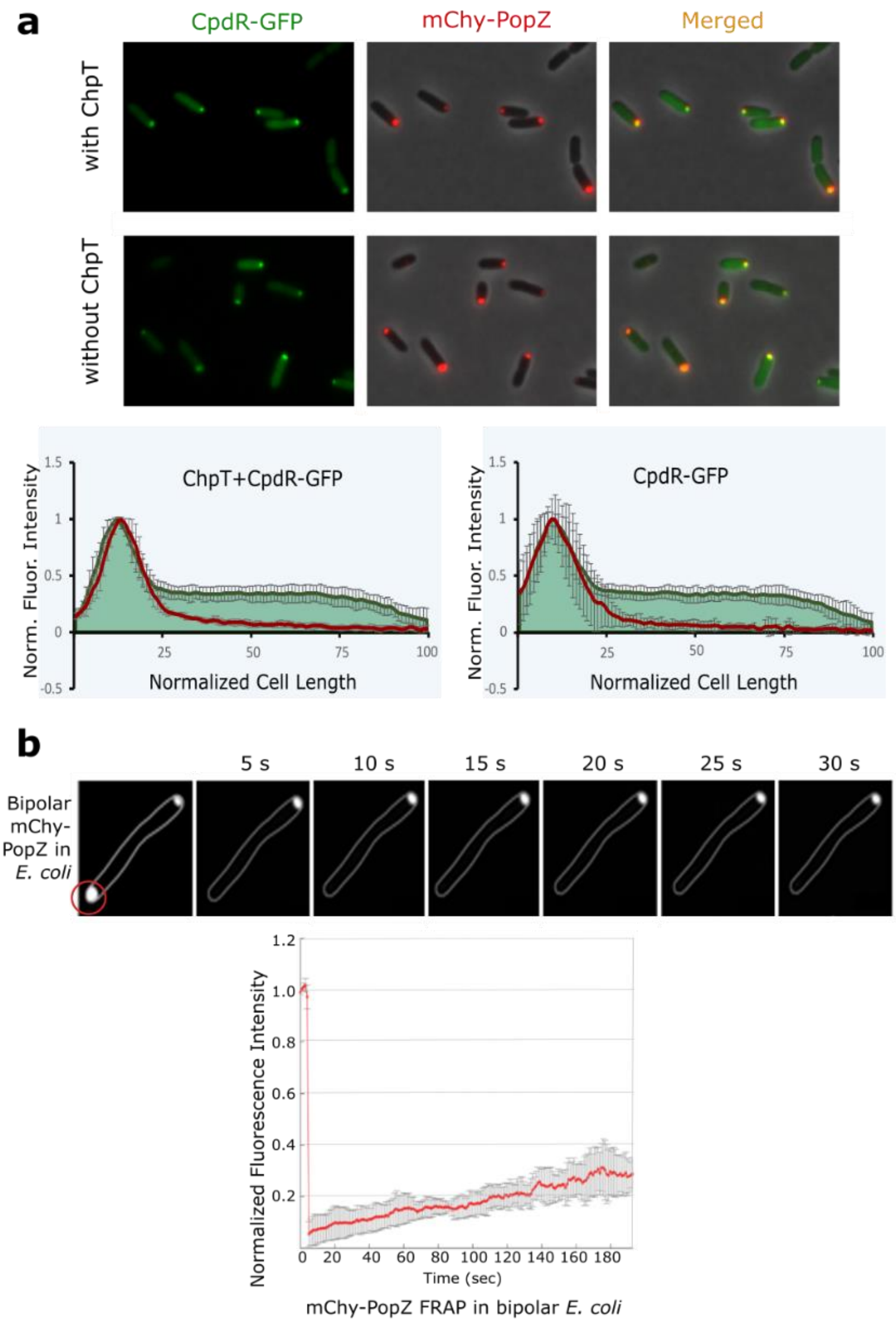


**Supplementary Fig. 1| Protein localization and dynamics in *E. coli*.**

**a,** *E. coli* co-expression assay, with mChy-PopZ and CpdR-GFP expressed in the presence or absence of the intermediary phosphotransferase ChpT. mChy-PopZ and CpdR-GFP were observed by microscopy and normalized fluorescence intensities were plotted against cell length (graphs). **b,** FRAP analysis of mChy-PopZ in *E. coli* cells with bipolar foci. Recovery of fluorescence was plotted against time in seconds.

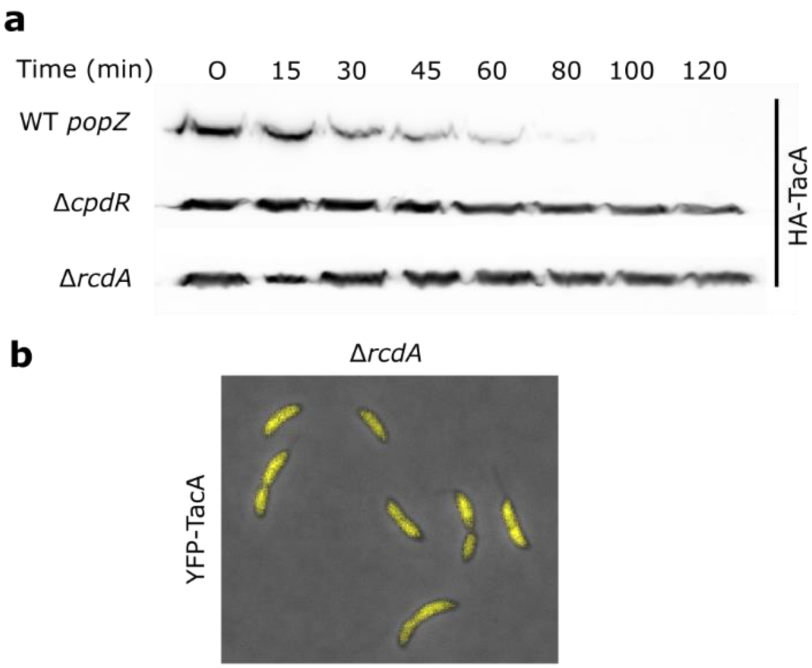


**Supplementary Fig. 2| TacA requires CpdR and RcdA adaptors for proteolysis and polar localization.**
**a,** Degradation of HA-TacA following inducer wash-out in Δ*popZ*, Δ*cpdR*, and Δ*rcdA C. crescentus* strain backgrounds, observed by western blotting with α-HA antibody. **b,** YFP-TacA localization in Δ*rcdA C. crescentus* strain background.

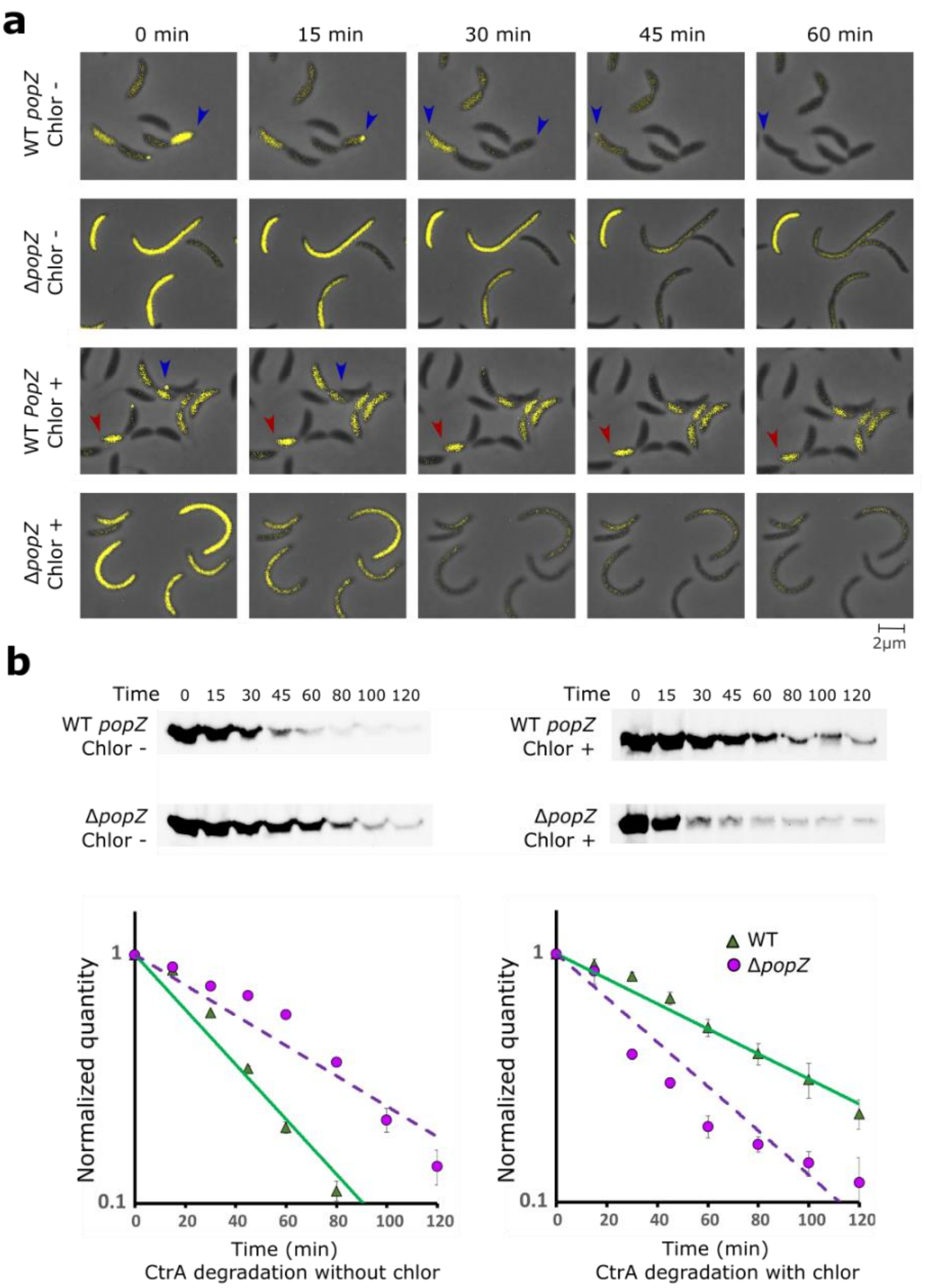


**Supplementary Fig. 3| Substrate degradation in the presence and absence of chloramphenicol.**
**a,** YFP-CtrA RD+15 distribution in WT and Δ*popZ C. crescentus* stain backgrounds with and without chloramphenicol (chlor) treatment. Blue arrowheads mark cells that exhibit YFP-CtrA RD+15 polar foci and degradation. Red arrows mark cells that do not exhibit YFP-CtrA RD+15 polar foci or degradation. **b,** Degradation of YFP-CtrA RD+15 following inducer wash-out in wildtype and Δ*popZ* strain backgrounds in the presence or absence of chloamphenicol treatment. Western blotting was performed with α-YFP antibody. Average band intensities from three separate experiments were plotted against time (graphs, bar = standard deviation).

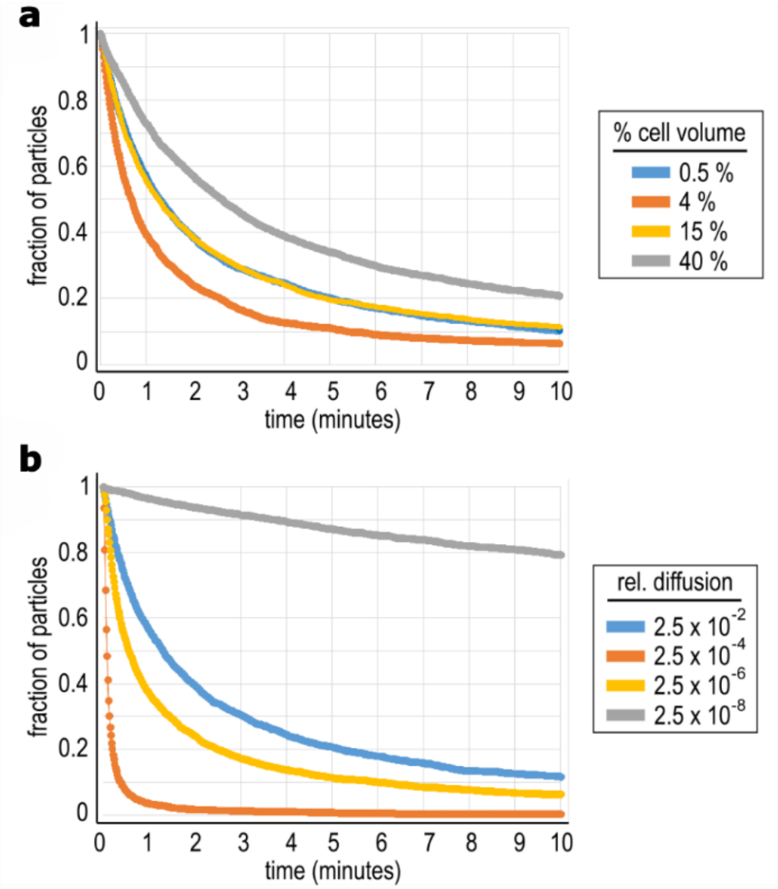


**Supplementary Fig. 4| Supplementary Fig. 4|Conceptual models of substrate proteolysis in membraneless polar microdomains, using extreme parameter values.** a-b, Three-dimensional reaction-diffusion simulations, as shown in Figure 5 a-b, except using different parameters. **a**, Cells with different sizes of membraneless polar microdomains, expressed as a percentage of total cell volume. 0.5% approximates the physiological size of PopZ microdomains *in C. crescentus*. At large microdomain volumes, the reaction slows because particles are more diffuse in a large compartment with a relatively slow particle diffusion rate. **b**, Cells with polar microdomains that have different particle diffusion rates. 2.5 x $10^{-2}$ (or 1/40) approximates the fold difference in diffusion rate between PopZ microdomains and bulk cytoplasm in *C. crescentus*. Up to a point, slower diffuion rates lead to higher particle concentration and faster reaction rates. However, when particle diffusion in polar microdomains is extremely slow, some particles become stuck just below the microdomain surface and their collision frequency is low.

**Supplementary Video 1|** Time-lapse video of FRAP. Fluorescence Recovery After Photobleaching in *E. coli* cell pole, showing a wavefront of CpdR-GFP advances through PopZ microdomain.

**Supplementary Video 2|** Time-lapse video of model #18. The top cell has a polar microdomain while the lower one does not. Particles in the polar microdomain diffuse at 1/40th their rate in bulk cytoplasm. Total simulation time was 600,000 msec. Sixty images, produced at intervals of 10,000 msec, were assembled for the video.

**Supplementary Video 3|** Time-lapse video of model #19. Particles in the top cell's polar microdomain diffuse at 1/10th their rate in bulk cytoplasm. Particles in the bottom cell's polar microdomain diffuse at 1/160th their rate in bulk cytoplasm. Total simulation time was 600,000 msec. Sixty images, produced at intervals of 10,000 msec, were assembled for the video.
