## Supplementary Tables for "Phospho-signaling couples polar asymmetry and proteolysis within a membraneless microdomain in *C. crescentus*"

**Supplementary Table 1**

| Strain name | Cell Background | Plasmid description | Source |
|---|---|---|---|
| **GB#1194** | *Caulobacter* wildtype synchronizable strain NA1000 CB15N | | [1] |
| **GB#1081** | *popZ*::*specR* | | [2] |
| **GB#1007** | *popZ::mChy-popZ* | | [3] |
| **GB#264** | *Caulobacter* WT; GB#1194 | pMR10 YFP-CtrA RD+15 | [4] |
| **GB#944** | *E. coli* BL21 DE3 | pBad-mChy PopZ | [5] |
| **GB#169** | *E. coli* Rosetta | pET28a-6XHis-PopZ | [2] |
| **JH#66** | *E. coli* DH5α | pBAD-6XHis-GFP | [6] |
| **GB#1969** | | pACYC-CpdR-TEV-msfGFP-His6X | this study |
| **GB#1970** | | pACYC-CpdRD51A-TEV-msfGFP-His6X | this study |
| **GB#2022** | *E. coli* MG1655 DE3 *clpXP*::*kanR* | | this study |
| **GB#1971** | *E. coli* MG1655 DE3 *clpXP*::*kanR* | pACYC-CpdRGFP-ChpT-CckA WT + pBAD-mCherry PopZ | this study |
| **GB#1972** | *E. coli* MG1655 DE3 *clpXP*::*kanR* | pACYC-CpdRGFP-ChpT-CckA H+ + pBAD-mCherry PopZ | this study |
| **GB#1973** | *E. coli* MG1655 DE3 *clpXP*::*kanR* | pACYC-CpdRGFP-ChpT-CckA K- + pBAD-mCherry PopZ | this study |
| **GB#1974** | *E. coli* MG1655 DE3 *clpXP*::*kanR* | pACYC-CpdR$_{D51A}$ GFP-ChpT-CckA H+ + pBAD-mCherry PopZ | this study |
| **GB#1975** | *E. coli* MG1655 DE3 *clpXP*::*kanR* | pACYC-CpdRGFP-ChpT + pBAD-mCherry PopZ | this study |
| **GB#1976** | *E. coli* MG1655 DE3 *clpXP*::*kanR* | pACYC-CpdRGFP + pBAD-mCherry PopZ | this study |
| **GB#1977** | *E. coli* MG1655 DE3 *clpXP*::*kanR* | pACYC-CpdRGFP + pBAD-mCherry ΔN term PopZ | this study |
| **GB#1978** | *popZ::mChy-popZ*; GB#1007 | PMCS5-pCpdR CpdR-YFP | this study |

| | | | |
|---|---|---|---|
| **GB#1979** | *popZ::mChy-popZ*; GB#1007 | PMCS5-pCpdR CpdR$_{D51A}$-YFP | this study |
| **GB#1980** | *popZ::mChy-popZ*; GB#1007 | PMCS5-pCpdR CpdR-YFP + pBXMCS2-CckA K+ | this study |
| **GB#1981** | *popZ::mChy-popZ*; GB#1007 | PMCS5-pCpdR CpdR-YFP + pBXMCS2-CckA H- | this study |
| **GB#1982** | *popZ::mChy-popZ*; GB#1007 | PMCS5-pCpdR CpdR$_{D51A}$-YFP + pBXMCS2-CckA K+ | this study |
| **GB#1983** | *Caulobacter* WT, GB#1194 | pXMCS5-YFP-CtrA RD+15 | this study |
| **GB#1984** | *Caulobacter* WT, GB#1194 | pXMCS5-YFP-TacA | this study |
| **GB#1985** | *Caulobacter* WT, GB#1194 | pXMCS5-YFP-PdeA | this study |
| **GB#1986** | *Caulobacter popZ::specR*, GB#1081 | pXMCS5-YFP-CtrA RD+15 | this study |
| **GB#1987** | *Caulobacter popZ::specR*, GB#1081 | pXMCS5-YFP-TacA | this study |
| **GB#1988** | *Caulobacter popZ::specR*, GB#1081 | pXMCS5-YFP-PdeA | this study |
| **GB#1989** | *Caulobacter* WT, GB#1194 | pXMCS5-HA-CtrA RD+15 | this study |
| **GB#1990** | *Caulobacter* WT, GB#1194 | pXMCS5-HA-TacA | this study |
| **GB#1991** | *Caulobacter* WT, GB#1194 | pXMCS5-HA-PdeA | this study |
| **GB#1992** | *Caulobacter popZ::specR*, GB#1081 | pXMCS5-HA-CtrA RD+15 | this study |
| **GB#1993** | *Caulobacter popZ::specR*, GB#1081 | pXMCS5-HA-TacA | this study |

| | | | |
|---|---|---|---|
| **GB#1994** | *Caulobacter popZ::specR*, GB#1081 | pXMCS5-HA-PdeA | this study |
| **GB#1995** | *cpdR::specR; popZ::mChy-popZ* | | this study |
| **GB#1996** | *ΔrcdA, GB#1106*[7] | pMCS5 pCpdR-YFP | this study |
| **GB#1997** | *cpdR::specR, popZ::mChy-popZ;* GB#1995 | pVMCS6-ClpX-msfGFP | this study |
| **GB#2002** | *cpdR::specR, popZ::mChy-popZ;* GB#1995 | pVMCS6-RcdA-msfGFP | this study |
| **GB#2007** | *cpdR::specR, popZ::mChy-popZ;* GB#1995 | pXMCS5-YFP-CtrA RD+15 | this study |
| **GB#2008** | *cpdR::specR, popZ::mChy-popZ;* GB#1995 | pXMCS5-YFP-TacA | this study |
| **GB#2009** | *cpdR::specR, popZ::mChy-popZ;* GB#1995 | pXMCS5-YFP-PdeA | this study |
| **GB#2014** | *cpdR::specR, popZ::mChy-popZ;* GB#1995 | pVMCS6-ClpX-GFP + pBXMCS2-pCpdR CpdR | this study |
| **GB#2015** | *cpdR::specR, popZ::mChy-popZ;* GB#1995 | pVMCS6-ClpX-GFP + pBXMCS2-pCpdR $CpdR_{D51A}$ | this study |
| **GB#2016** | *cpdR::specR, popZ::mChy-popZ;* GB#1995 | pVMCS6-ClpX-GFP + pBXMCS2-pXyl-CckA K+-pCpdR CpdR | this study |
| **GB#2017** | *cpdR::specR, popZ::mChy-popZ;* GB#1995 | pVMCS6-ClpX-GFP + pBXMCS2-pXyl-CckA K+-pCpdR $CpdR_{D51A}$ | this study |

| | | | |
|---|---|---|---|
| **GB#2018** | *cpdR*::*specR, popZ::mChy-popZ;* GB#1995 | pVMCS6-RcdA-GFP + pBXMCS2-pCpdR CpdR | this study |
| **GB#2019** | *cpdR*::*specR, popZ::mChy-popZ;* GB#1995 | pVMCS6-RcdA-GFP + pBXMCS2-pCpdR $CpdR_{D51A}$ | this study |
| **GB#2020** | *cpdR*::*specR, popZ::mChy-popZ;* GB#1995 | pVMCS6-RcdA-GFP + pBXMCS2-pXyl-CckA K+-pCpdR CpdR | this study |
| **GB#2021** | *cpdR*::*specR, popZ::mChy-popZ;* GB#1995 | pVMCS6-RcdA-GFP + pBXMCS2-pXyl-CckA K+-pCpdR $CpdR_{D51A}$ | this study |


1. Evinger, M. & Agabian, N. Envelope-associated nucleoid from Caulobacter crescentus stalked and swarmer cells. *J Bacteriol* **132**, 294–301 (1977).

2. Bowman, G. R. *et al.* A Polymeric Protein Anchors the Chromosomal Origin/ParB Complex at a Bacterial Cell Pole. *Cell* **134**, 945–955 (2008).

3. Bowman, G. R. *et al.* Oligomerization and higher-order assembly contribute to sub-cellular localization of a bacterial scaffold. *Molecular Microbiology* **90**, 776–795 (2013).

4. Ryan, K. R., Judd, E. M. & Shapiro, L. The CtrA Response Regulator Essential for Caulobacter crescentus Cell-cycle Progression Requires a Bipartite Degradation Signal for Temporally Controlled Proteolysis. *Journal of Molecular Biology* **324**, 443–455 (2002).

5. Perez, A. M. Assembly of a Sub-Cellular Protein Complex Generates Asymmetry in the Bacterium Caulobacter crescentus. (Stanford University, 2015).

6. Holmes, J. A. *et al.* Caulobacter PopZ forms an intrinsically disordered hub in organizing bacterial cell poles. *Proc Natl Acad Sci U S A* **113**, 12490–12495 (2016).

7. McGrath, P. T., Iniesta, A. A., Ryan, K. R., Shapiro, L. & McAdams, H. H. A Dynamically Localized Protease Complex and a Polar Specificity Factor Control a Cell Cycle Master Regulator. *Cell* **124**, 535–547 (2006).

**Supplementary Table 2**

| Plasmid name | Vector backbone | Description | Sequence<br>*where present, ellipsis correspond to vector backbone sequence |
|---|---|---|---|
| **pGB1945** | pCDF-duet | pCDF-CckA WT | gagatataccATGGCCGACTTGCAGCTCCAGGACAAGGTTTCGACCGGCGCCCCGCGTCGGCGGTTT<br>GATCCATGGCTGGTCGGCGCGGCGGTGTTTTTCGTGGCTGCGGCGGCCCTCTCGGCGGCGCCGG<br>CGCTCAAGGCCGGACCGACCACCCTGGCCGGCCTGCTGCTGCTTCTGGGCGTCGCGGGCGTGGC<br>TGTGTTGGGCCTTGTCGCCATTCGCGGCTCAGCGCTTTCCGGCGGCGACGCCGACCAGGCTGAG<br>GGGTTCATCGAGGCGCTGGCCGAGCCGGCCGCCCTGGCCGCCGCCGACGGTCGCGTCCTGGCCG<br>CCAACGGGCCCTGGCGCGAAGTCATGGGCGAACAGCGCCGCCTGCCCAAGGGCGTGGCGGGCT<br>CCAGCCTGTTTGCGGCCCTGGTCCAGGCGCGCCAGGGGCAGATGGCCGAGGGCATGCTGAGCGC<br>TGGAGGAACCGACTATACCGCCAAGGTCTCGCGCCTTGCGGGCGGACGGCTGATGATCCGGCTTG<br>CGCCCATCGTTGTCGCTGAGCCGGTTGTGGAAGACGCGTCGCCGGCCCCGGTCGCCGAGCGCGC<br>CGCCCCGCCACCCAGCTCGCTGGACGCCTTCGCCGGAGCCTCGCCGTTCGGCGCGGCCCTGCTGG<br>AAGGCCTGGAGCCGTTCACCTCGCGGGTGCTGGAGACCAATCCGGCGCTGACCACGATGACCGG<br>CGCCAAGGCCGGTGTGCTGTTCGGCGATCTGATCGACGCCGCCTCGCGCGCCGAGGCCGAGACG<br>CGCCTGAACGAAGGCCGCGCCGGTCCCTACGAGGTGCGGTTGGCGCGCGATCCGTCGCGCATCG<br>CTCACCTTTACCTCTATCGCGCCGAAGGCCGCCTGGTGGCCTACATGATCGACGTGTCCGAGCAGA<br>AGCAGATCGAGCTGCAGCTGTCCCAGGCCCAGAAGATGCAGGCCATCGGCCAGCTGGCCGGCGG<br>CGTCGCGCACGACTTCAACAACCTCTTGACCGCCATCCAGCTGCGTCTGGACGAGTTGCTGCATCG<br>CCATCCCGTCGGTGATCCGTCGTACGAGGGTCTCAACGAGATCCGTCAGACGGGCGTGCGCGCCG<br>CCGACCTCGTGCGCAAGCTCTTGGCTTTCTCGCGCAAGCAGACCGTGCAGCGCGAGGTGCTGGAT<br>CTGGGCGAGCTGATCAGCGAGTTCGAGGTCTTGCTGCGCCGCCTGCTGCGCGAAGACGTCAAGC<br>TGATCACCGACTATGGCCGCGACCTGCCGCAGGTGCGCGCCGACAAGAGCCAGCTCGAGACGGC<br>GGTCATGAACCTGGCCGTCAACGCCCGCGACGCCGTGCGCGCGGCCAAGGGCGGCGGCGTCGT<br>GCGCATCCGCACCGCGCGCCTGACCCGCGACGAGGCGATCCAGCTGGGTTTCCCGGCCGCCGAC<br>GGCGACACGGCCTTCATTGAGGTCAGTGACGATGGTCCGGGCATTCCGCCCGACGTCATGGGCAA<br>GATCTTCGACCCGTTCTTCACCACCAAGCCGGTGGGCGAGGGTACGGGCCTAGGCCTAGCCACGG<br>TCTATGGCATCGTTAAGCAGAGCGACGGCTGGATTCACGTCCACAGCCGTCCGAACGAAGGCGCG<br>GCCTTCCGCATCTTCCTGCCGGTCTATGAAGCGCCCGCCGGCGCGGTCGCCGTCCAGGCCGTCGC<br>CGAGCCCGCCAAGCCGCGCGCCGCTCGCGACCTGTCGGGCGCCGGCCGCATCCTGTTCGTCGAG<br>GACGAGGACGCCGTGCGCAGCGTCGCCGCCCGCCTGCTGCGCGCCCGTGGCTACGAGGTGCTTG<br>AGGCGGCCGACGGCGAAGAGGCCCTGATCATCGCCGAGGAAAACGCCGGCACGATCGACCTCTT<br>GATCAGCGACGTGATCATGCCCGGCATCGACGGCCCGACCTTGCTGAAGAAGGCGCGTGGCTATC<br>TGGGGACCGCGCCGGTGATGTTCATCTCCGGCTATGCCGAGGCCGAGTTCAGCGACCTCTTGGAA<br>GGCGAGACGGGCGTGACCTTCCTCCCCAAGCCGATCGACATCAAGACCCTGGCCGAGCGCGTCA<br>AGCAGCAGCTGCAGGCGGCGTAGgcggccgcat |
| **pGB1946** | pCDF-duet | pCDF-CckA H+ | gagatataccATGGCCGACTTGCAGCTCCAGGACAAGGTTTCGACCGGCGCCCCGCGTCGGCGGTTT<br>GATCCATGGCTGGTCGGCGCGGCGGTGTTTTTCGTGGCTGCGGCGGCCCTCTCGGCGGCGCCGG<br>CGCTCAAGGCCGGACCGACCACCCTGGCCGGCCTGCTGCTGCTTCTGGGCGTCGCGGGCGTGGC<br>TGTGTTGGGCCTTGTCGCCATTCGCGGCTCAGCGCTTTCCGGCGGCGACGCCGACCAGGCTGAG<br>GGGTTCATCGAGGCGCTGGCCGAGCCGGCCGCCCTGGCCGCCGCCGACGGTCGCGTCCTGGCCG<br>CCAACGGGCCCTGGCGCGAAGTCATGGGCGAACAGCGCCGCCTGCCCAAGGGCGTGGCGGGCT<br>CCAGCCTGTTTGCGGCCCTGGTCCAGGCGCGCCAGGGGCAGATGGCCGAGGGCATGCTGAGCGC<br>TGGAGGAACCGACTATACCGCCAAGGTCTCGCGCCTTGCGGGCGGACGGCTGATGATCCGGCTTG<br>CGCCCATCGTTGTCGCTGAGCCGGTTGTGGAAGACGCGTCGCCGGCCCCGGTCGCCGAGCGCGC<br>CGCCCCGCCACCCAGCTCGCTGGACGCCTTCGCCGGAGCCTCGCCGTTCGGCGCGGCCCTGCTGG<br>AAGGCCTGGAGCCGTTCACCTCGCGGGTGCTGGAGACCAATCCGGCGCTGACCACGATGACCGG<br>CGCCAAGGCCGGTGTGCTGTTCGGCGATCTGATCGACGCCGCCTCGCGCGCCGAGGCCGAGACG<br>CGCCTGAACGAAGGCCGCGCCGGTCCCTACGAGGTGCGGTTGGCGCGCGATCCGTCGCGCATCG<br>CTCACCTTTACCTCTATCGCGCCGAAGGCCGCCTGGTGGCCTACATGATCGACGTGTCCGAGCAGA<br>AGCAGATCGAGCTGCAGCTGTCCCAGGCCCAGAAGATGCAGGCCATCGGCCAGCTGGCCGGCGA |

| | | | |
|---|---|---|---|
| | | | AGTCGCGCACGACTTCAACAACCTCTTGACCGCCATCCAGCTGCGTCTGGACGAGTTGCTGCATCG<br>CCATCCCGTCGGTGATCCGTCGTACGAGGGTCTCAACGAGATCCGTCAGACGGGCGTGCGCGCCG<br>CCGACCTCGTGCGCAAGCTCTTGGCTTTCTCGCGCAAGCAGACCGTGCAGCGCGAGGTGCTGGAT<br>CTGGGCGAGCTGATCAGCGAGTTCGAGGTCTTGCTGCGCCGCCTGCTGCGCGAAGACGTCAAGC<br>TGATCACCGACTATGGCCGCGACCTGCCGCAGGTGCGCGCCGACAAGAGCCAGCTCGAGACGGC<br>GGTCATGAACCTGGCCGTCAACGCCCGCGACGCCGTGCGCGCGGCCAAGGGCGGCGGCGTCGT<br>GCGCATCCGCACCGCGCGCCTGACCCGCGACGAGGCGATCCAGCTGGGTTTCCCGGCCGCCGAC<br>GGCGACACGGCCTTCATTGAGGTCAGTGACGATGGTCCGGGCATTCCGCCCGACGTCATGGGCAA<br>GATCTTCGACCCGTTCTTCACCACCAAGCCGGTGGGCGAGGGTACGGGCCTAGGCCTAGCCACGG<br>TCTATGGCATCGTTAAGCAGAGCGACGGCTGGATTCACGTCCACAGCCGTCCGAACGAAGGCGCG<br>GCCTTCCGCATCTTCCTGCCGGTCTATGAAGCGCCCGCCGGCGCGGTCGCCGTCCAGGCCGTCGC<br>CGAGCCCGCCAAGCCGCGCGCCGCTCGCGACCTGTCGGGCGCCGGCCGCATCCTGTTCGTCGAG<br>GACGAGGACGCCGTGCGCAGCGTCGCCGCCCGCCTGCTGCGCGCCCGTGGCTACGAGGTGCTTG<br>AGGCGGCCGACGGCGAAGAGGCCCTGATCATCGCCGAGGAAAACGCCGGCACGATCGACCTCTT<br>GATCAGCGACGTGATCATGCCCGGCATCGACGGCCCGACCTTGCTGAAGAAGGCGCGTGGCTATC<br>TGGGGACCGCGCCGGTGATGTTCATCTCCGGCTATGCCGAGGCCGAGTTCAGCGACCTCTTGGAA<br>GGCGAGACGGGCGTGACCTTCCTCCCCAAGCCGATCGACATCAAGACCCTGGCCGAGCGCGTCA<br>AGCAGCAGCTGCAGGCGGCGTAGgcggccgcat |
| **pGB1947** | pCDF-duet | pCDF-CckAK- | gagatataccATGGCCGACTTGCAGCTCCAGGACAAGGTTTCGACCGGCGCCCCGCGTCGGCGGTTT<br>GATCCATGGCTGGTCGGCGCGGCGGTGTTTTTCGTGGCTGCGGCGGCCCTCTCGGCGGCGCCGG<br>CGCTCAAGGCCGGACCGACCACCCTGGCCGGCCTGCTGCTGCTTCTGGGCGTCGCGGGCGTGGC<br>TGTGTTGGGCCTTGTCGCCATTCGCGGCTCAGCGCTTTCCGGCGGCGACGCCGACCAGGCTGAG<br>GGGTTCATCGAGGCGCTGGCCGAGCCGGCCGCCCTGGCCGCCGCCGACGGTCGCGTCCTGGCCG<br>CCAACGGGCCCTGGCGCGAAGTCATGGGCGAACAGCGCCGCCTGCCCAAGGGCGTGGCGGGCT<br>CCAGCCTGTTTGCGGCCCTGGTCCAGGCGCGCCAGGGGCAGATGGCCGAGGGCATGCTGAGCGC<br>TGGAGGAACCGACTATACCGCCAAGGTCTCGCGCCTTGCGGGCGGACGGCTGATGATCCGGCTTG<br>CGCCCATCGTTGTCGCTGAGCCGGTTGTGGAAGACGCGTCGCCGGCCCCGGTCGCCGAGCGCGC<br>CGCCCCGCCACCCAGCTCGCTGGACGCCTTCGCCGGAGCCTCGCCGTTCGGCGCGGCCCTGCTGG<br>AAGGCCTGGAGCCGTTCACCTCGCGGGTGCTGGAGACCAATCCGGCGCTGACCACGATGACCGG<br>CGCCAAGGCCGGTGTGCTGTTCGGCGATCTGATCGACGCCGCCTCGCGCGCCGAGGCCGAGACG<br>CGCCTGAACGAAGGCCGCGCCGGTCCCTACGAGGTGCGGTTGGCGCGCGATCCGTCGCGCATCG<br>CTCACCTTTACCTCTATCGCGCCGAAGGCCGCCTGGTGGCCTACATGATCGACGTGTCCGAGCAGA<br>AGCAGATCGAGCTGCAGCTGTCCCAGGCCCAGAAGATGCAGGCCATCGGCCAGCTGGCCGGCGG<br>CGTCGCGCACGACTTCAACAACCTCTTGACCGCCATCCAGCTGCGTCTGGACGAGTTGCTGCATCG<br>CCATCCCGTCGGTGATCCGTCGTACGAGGGTCTCAACGAGATCCGTCAGACGGGCGTGCGCGCCG<br>CCGACCTCGTGCGCAAGCTCTTGGCTTTCTCGCGCAAGCAGACCGTGCAGCGCGAGGTGCTGGAT<br>CTGGGCGAGCTGATCAGCGAGTTCGAGGTCTTGCTGCGCCGCCTGCTGCGCGAAGACGTCAAGC<br>TGATCACCGACTATGGCCGCGACCTGCCGCAGGTGCGCGCCGACAAGAGCCAGCTCGAGACGGC<br>GGTCATGAACCTGGCCGTCAACGCCCGCGACGCCGTGCGCGCGGCCAAGGGCGGCGGCGTCGT<br>GCGCATCCGCACCGCGCGCCTGACCCGCGACGAGGCGATCCAGCTGGGTTTCCCGGCCGCCGAC<br>GGCGACACGGCCTTCATTGAGGTCAGTGACGATGGTCCGGGCATTCCGCCCGACGTCATGGGCAA<br>GATCTTCGACCCGTTCTTCACCACCAAGCCGGTGGGCGAGGGTACGGGCCTAGGCCTAGCCACGG<br>TCTATGGCATCGTTAAGCAGAGCGACGGCTGGATTCACGTCCACAGCCGTCCGAACGAAGGCGCG<br>GCCTTCCGCATCTTCCTGCCGGTCTATGAAGCGCCCGCCGGCGCGGTCGCCGTCCAGGCCGTCGC<br>CGAGCCCGCCAAGCCGCGCGCCGCTCGCGACCTGTCGGGCGCCGGCCGCATCCTGTTCGTCGAG<br>GACGAGGACGCCGTGCGCAGCGTCGCCGCCCGCCTGCTGCGCGCCCGTGGCTACGAGGTGCTTG<br>AGGCGGCCGACGGCGAAGAGGCCCTGATCATCGCCGAGGAAAACGCCGGCACGATCGACCTCTT<br>GATCAGCGACGTGATCATGCCCGGCATCGACGGCCCGACCTTGCTGAAGAAGGCGCGTGGCTATC<br>TGGGGACCGCGCCGGTGATGTTCATCTCCGGCTATGCCGAGGCCGAGTTCAGCGACCTCTTGGAA<br>GGCGAGACGGGCGTGACCTTCCTCCCCAAGCCGATCGACATCAAGACCCTGGCCGAGCGCGTCA<br>AGCAGCAGCTGCAGGCGGCGTAGgcggccgcat |

| | | | |
|---|---|---|---|
| **pGB1948** | pACYC-duet | pACYC-CpdRGFP-ChpT-CckAWT | gagatataccGTGGCCCGCATCCTCCTCGCCGAAGACGATGATTCCCTGCGCGGCTTCCTGGCCCGCG<br>CGCTGGAACGCGCCGGCTTCGAAGTCCAGGCCTGCGCCGACGGCGAAGAGGCCGTCCAGCACCT<br>GGACCATCCCTGGGACCTGCTGCTGACCGACATCGTCATGCCCGGCATGGACGGCATCGAGGTGG<br>CCCGCCAGGCCGCCGCCCGCGACCCGTCCCTGCGCATCATGTTCATCACCGGCTTCGCCGCCGTGG<br>CCCTCTCGGCCCAGGACCGCGCGCCCGCCGGCGCCAAGGTGCTGTCCAAGCCCGTGCACCTGCG<br>CGACCTCGTCGCCGAGGTCGAAAAGATGATGGCGGCCCTCGAGAGCGGCGGCGGCGGCAGTAA<br>AGGTGAAGAACTGTTCACCGGTGTTGTTCCGATCCTGGTTGAACTGGATGGTGATGTTAACGGCC<br>ACAAATTCTCTGTTCGTGGTGAAGGTGAAGGTGATGCAACCAACGGTAAACTGACCCTGAAATTC<br>ATCTGCACTACCGGTAAACTGCCGGTTCCATGGCCGACTCTGGTGACTACCCTGACCTATGGTGTTC<br>AGTGTTTTTCTCGTTACCCGGATCACATGAAGCAGCATGATTTCTTCAAATCTGCAATGCCGGAAGG<br>TTATGTACAGGAGCGCACCATTTCTTTCAAAGACGATGGCACCTACAAAACCCGTGCAGAGGTTAA<br>ATTTGAAGGTGATACTCTGGTGAACCGTATTGAACTGAAAGGCATTGATTTCAAAGAGGACGGCA<br>ACATCCTGGGCCACAAACTGGAATATAACTTCAACTCCCATAACGTTTACATCACCGCAGACAAACA<br>GAAGAACGGTATCAAAGCTAACTTCAAAATTCGCCATAACGTTGAAGACGGTAGCGTACAGCTGG<br>CGGACCACTACCAGCAGAACACTCCGATCGGTGATGGTCCGGTTCTGCTGCCGGATAACCACTACC<br>TGTCCACCCAGTCTAAACTGTCCAAAGACCCGAACGAAAAGCGCGACCACATGGTGCTGCTGGAG<br>TTCGTTACTGCAGCAGGTATCACGCACGGCATGGATGAACTCTACAAATGAgcggccgcat.........agat<br>atacatGTGACCGAGACCGTCACCGAGACCACCGCCCCCGCGTCCCCCGAAGCCGACGTCCAGGGT<br>CCCGATTTCGCCGCCATGCTGGCCGCGCGCCTGTGTCACGACTTCATCAGTCCGGCCAGCGCCATC<br>GTCTCGGGCCTGGATCTGCTGGAAGACCCCTCGGCCCAGGACATGCGCGACGACGCCATGAACCT<br>GATCGCCTCCTCGGCCCGCAAGCTGGCGGACCTCTTGCAGTTCACCCGAGTGGCCTTCGGCGCCT<br>CGGCCTCGGCCGAGAACTTCGACTCGCGCGAACTGGAAAAGCTGGCCCAGGGCGTCTTTGCCCA<br>TGTCCGCCCGACCCTGGACTGGCAGATCGAGCCGCAGGCGATGAACAAGCCCTCGTCGCGCGCG<br>GTGCTGAACATCGCCCAGATCGCCGCCAGCGCCCTGCCGGCCGGCGGCGTGGCCACCGTCAAGG<br>GCGTGGCCGCCGACGGGCGCTTCTCGATCATCGCCGACGCCAAGGGCCCGCGCGCGCGCCTGCG<br>TCCGGAGGTGCTGGCGGGCCTAAAGGGCGAGCCGCTGGCCGAGGGCCTGGGCGGTCCGTGGGT<br>GCAGGCGGCCTATCTGAACGCCCTGGTGCGCGCGGCCGGCGGCCAGATCGCCGTCGAGATCGGC<br>GAGGACCGCGCCTCGATCGCCGCCTGGGTCCCGGCGTAActcgagtctg.........tcagcgctagttatgcgac<br>tcctgcattaggaaattaatacgactcactataggggaattgtgagcggataacaattcccctgtagaaataattttgtttaacttt<br>aataaggagatataccATGGCCGACTTGCAGCTCCAGGACAAGGTTTCGACCGGCGCCCCGCGTCGGC<br>GGTTTGATCCATGGCTGGTCGGCGCGGCGGTGTTTTTCGTGGCTGCGGCGGCCCTCTCGGCGGCG<br>CCGGCGCTCAAGGCCGGACCGACCACCCTGGCCGGCCTGCTGCTGCTTCTGGGCGTCGCGGGCG<br>TGGCTGTGTTGGGCCTTGTCGCCATTCGCGGCTCAGCGCTTTCCGGCGGCGACGCCGACCAGGCT<br>GAGGGGTTCATCGAGGCGCTGGCCGAGCCGGCCGCCCTGGCCGCCGCCGACGGTCGCGTCCTGG<br>CCGCCAACGGGCCCTGGCGCGAAGTCATGGGCGAACAGCGCCGCCTGCCCAAGGGCGTGGCGG<br>GCTCCAGCCTGTTTGCGGCCCTGGTCCAGGCGCGCCAGGGGCAGATGGCCGAGGGCATGCTGAG<br>CGCTGGAGGAACCGACTATACCGCCAAGGTCTCGCGCCTTGCGGGCGGACGGCTGATGATCCGG<br>CTTGCGCCCATCGTTGTCGCTGAGCCGGTTGTGGAAGACGCGTCGCCGGCCCCGGTCGCCGAGC<br>GCGCCGCCCCGCCACCCAGCTCGCTGGACGCCTTCGCCGGAGCCTCGCCGTTCGGCGCGGCCCT<br>GCTGGAAGGCCTGGAGCCGTTCACCTCGCGGGTGCTGGAGACCAATCCGGCGCTGACCACGATG<br>ACCGGCGCCAAGGCCGGTGTGCTGTTCGGCGATCTGATCGACGCCGCCTCGCGCGCCGAGGCCG<br>AGACGCGCCTGAACGAAGGCCGCGCCGGTCCCTACGAGGTGCGGTTGGCGCGCGATCCGTCGCG<br>CATCGCTCACCTTTACCTCTATCGCGCCGAAGGCCGCCTGGTGGCCTACATGATCGACGTGTCCGA<br>GCAGAAGCAGATCGAGCTGCAGCTGTCCCAGGCCCAGAAGATGCAGGCCATCGGCCAGCTGGCC<br>GGCGGCGTCGCGCACGACTTCAACAACCTCTTGACCGCCATCCAGCTGCGTCTGGACGAGTTGCT<br>GCATCGCCATCCCGTCGGTGATCCGTCGTACGAGGGTCTCAACGAGATCCGTCAGACGGGCGTGC<br>GCGCCGCCGACCTCGTGCGCAAGCTCTTGGCTTTCTCGCGCAAGCAGACCGTGCAGCGCGAGGT<br>GCTGGATCTGGGCGAGCTGATCAGCGAGTTCGAGGTCTTGCTGCGCCGCCTGCTGCGCGAAGAC<br>GTCAAGCTGATCACCGACTATGGCCGCGACCTGCCGCAGGTGCGCGCCGACAAGAGCCAGCTCG<br>AGACGGCGGTCATGAACCTGGCCGTCAACGCCCGCGACGCCGTGCGCGCGGCCAAGGGCGGCG<br>GCGTCGTGCGCATCCGCACCGCGCGCCTGACCCGCGACGAGGCGATCCAGCTGGGTTTCCCGGCC<br>GCCGACGGCGACACGGCCTTCATTGAGGTCAGTGACGATGGTCCGGGCATTCCGCCCGACGTCAT<br>GGGCAAGATCTTCGACCCGTTCTTCACCACCAAGCCGGTGGGCGAGGGTACGGGCCTAGGCCTA<br>GCCACGGTCTATGGCATCGTTAAGCAGAGCGACGGCTGGATTCACGTCCACAGCCGTCCGAACGA |

| | | | |
|---|---|---|---|
| | | | AGGCGCGGCCTTCCGCATCTTCCTGCCGGTCTATGAAGCGCCCGCCGGCGCGGTCGCCGTCCAGG<br>CCGTCGCCGAGCCCGCCAAGCCGCGCGCCGCTCGCGACCTGTCGGGCGCCGGCCGCATCCTGTTC<br>GTCGAGGACGAGGACGCCGTGCGCAGCGTCGCCGCCCGCCTGCTGCGCGCCCGTGGCTACGAGG<br>TGCTTGAGGCGGCCGACGGCGAAGAGGCCCTGATCATCGCCGAGGAAAACGCCGGCACGATCGA<br>CCTCTTGATCAGCGACGTGATCATGCCCGGCATCGACGGCCCGACCTTGCTGAAGAAGGCGCGTG<br>GCTATCTGGGGACCGCGCCGGTGATGTTCATCTCCGGCTATGCCGAGGCCGAGTTCAGCGACCTCT<br>TGGAAGGCGAGACGGGCGTGACCTTCCTCCCCAAGCCGATCGACATCAAGACCCTGGCCGAGCG<br>CGTCAAGCAGCAGCTGCAGGCGGCGTAGcggagtgtat |
| **pGB1949** | pACYC-duet | pACYC-CpdRGFP-ChpT-CckA H+ | gagatataccGTGGCCCGCATCCTCCTCGCCGAAGACGATGATTCCCTGCGCGGCTTCCTGGCCCGCG<br>CGCTGGAACGCGCCGGCTTCGAAGTCCAGGCCTGCGCCGACGGCGAAGAGGCCGTCCAGCACCT<br>GGACCATCCCTGGGACCTGCTGCTGACCGACATCGTCATGCCCGGCATGGACGGCATCGAGGTGG<br>CCCGCCAGGCCGCCGCCCGCGACCCGTCCCTGCGCATCATGTTCATCACCGGCTTCGCCGCCGTGG<br>CCCTCTCGGCCCAGGACCGCGCGCCCGCCGGCGCCAAGGTGCTGTCCAAGCCCGTGCACCTGCG<br>CGACCTCGTCGCCGAGGTCGAAAAGATGATGGCGGCCCTCGAGAGCGGCGGCGGCGGCAGTAA<br>AGGTGAAGAACTGTTCACCGGTGTTGTTCCGATCCTGGTTGAACTGGATGGTGATGTTAACGGCC<br>ACAAATTCTCTGTTCGTGGTGAAGGTGAAGGTGATGCAACCAACGGTAAACTGACCCTGAAATTC<br>ATCTGCACTACCGGTAAACTGCCGGTTCCATGGCCGACTCTGGTGACTACCCTGACCTATGGTGTTC<br>AGTGTTTTTCTCGTTACCCGGATCACATGAAGCAGCATGATTTCTTCAAATCTGCAATGCCGGAAGG<br>TTATGTACAGGAGCGCACCATTTCTTTCAAAGACGATGGCACCTACAAAACCCGTGCAGAGGTTAA<br>ATTTGAAGGTGATACTCTGGTGAACCGTATTGAACTGAAAGGCATTGATTTCAAAGAGGACGGCA<br>ACATCCTGGGCCACAAACTGGAATATAACTTCAACTCCCATAACGTTTACATCACCGCAGACAAACA<br>GAAGAACGGTATCAAAGCTAACTTCAAAATTCGCCATAACGTTGAAGACGGTAGCGTACAGCTGG<br>CGGACCACTACCAGCAGAACACTCCGATCGGTGATGGTCCGGTTCTGCTGCCGGATAACCACTACC<br>TGTCCACCCAGTCTAAACTGTCCAAAGACCCGAACGAAAAGCGCGACCACATGGTGCTGCTGGAG<br>TTCGTTACTGCAGCAGGTATCACGCACGGCATGGATGAACTCTACAAATGAgcggccgcat.........agat<br>atacatGTGACCGAGACCGTCACCGAGACCACCGCCCCCGCGTCCCCCGAAGCCGACGTCCAGGGT<br>CCCGATTTCGCCGCCATGCTGGCCGCGCGCCTGTGTCACGACTTCATCAGTCCGGCCAGCGCCATC<br>GTCTCGGGCCTGGATCTGCTGGAAGACCCCTCGGCCCAGGACATGCGCGACGACGCCATGAACCT<br>GATCGCCTCCTCGGCCCGCAAGCTGGCGGACCTCTTGCAGTTCACCCGAGTGGCCTTCGGCGCCT<br>CGGCCTCGGCCGAGAACTTCGACTCGCGCGAACTGGAAAAGCTGGCCCAGGGCGTCTTTGCCCA<br>TGTCCGCCCGACCCTGGACTGGCAGATCGAGCCGCAGGCGATGAACAAGCCCTCGTCGCGCGCG<br>GTGCTGAACATCGCCCAGATCGCCGCCAGCGCCCTGCCGGCCGGCGGCGTGGCCACCGTCAAGG<br>GCGTGGCCGCCGACGGGCGCTTCTCGATCATCGCCGACGCCAAGGGCCCGCGCGCGCGCCTGCG<br>TCCGGAGGTGCTGGCGGGCCTAAAGGGCGAGCCGCTGGCCGAGGGCCTGGGCGGTCCGTGGGT<br>GCAGGCGGCCTATCTGAACGCCCTGGTGCGCGCGGCCGGCGGCCAGATCGCCGTCGAGATCGGC<br>GAGGACCGCGCCTCGATCGCCGCCTGGGTCCCGGCGTAActcgagtctg.........tcagcgctagttatgcgac<br>tcctgcattaggaaattaatacgactcactataggggaattgtgagcggataacaattcccctgtagaaataattttgtttaacttt<br>aataaggagatataccATGGCCGACTTGCAGCTCCAGGACAAGGTTTCGACCGGCGCCCCGCGTCGGC<br>GGTTTGATCCATGGCTGGTCGGCGCGGCGGTGTTTTTCGTGGCTGCGGCGGCCCTCTCGGCGGCG<br>CCGGCGCTCAAGGCCGGACCGACCACCCTGGCCGGCCTGCTGCTGCTTCTGGGCGTCGCGGGCG<br>TGGCTGTGTTGGGCCTTGTCGCCATTCGCGGCTCAGCGCTTTCCGGCGGCGACGCCGACCAGGCT<br>GAGGGGTTCATCGAGGCGCTGGCCGAGCCGGCCGCCCTGGCCGCCGCCGACGGTCGCGTCCTGG<br>CCGCCAACGGGCCCTGGCGCGAAGTCATGGGCGAACAGCGCCGCCTGCCCAAGGGCGTGGCGG<br>GCTCCAGCCTGTTTGCGGCCCTGGTCCAGGCGCGCCAGGGGCAGATGGCCGAGGGCATGCTGAG<br>CGCTGGAGGAACCGACTATACCGCCAAGGTCTCGCGCCTTGCGGGCGGACGGCTGATGATCCGG<br>CTTGCGCCCATCGTTGTCGCTGAGCCGGTTGTGGAAGACGCGTCGCCGGCCCCGGTCGCCGAGC<br>GCGCCGCCCCGCCACCCAGCTCGCTGGACGCCTTCGCCGGAGCCTCGCCGTTCGGCGCGGCCCT<br>GCTGGAAGGCCTGGAGCCGTTCACCTCGCGGGTGCTGGAGACCAATCCGGCGCTGACCACGATG<br>ACCGGCGCCAAGGCCGGTGTGCTGTTCGGCGATCTGATCGACGCCGCCTCGCGCGCCGAGGCCG<br>AGACGCGCCTGAACGAAGGCCGCGCCGGTCCCTACGAGGTGCGGTTGGCGCGCGATCCGTCGCG<br>CATCGCTCACCTTTACCTCTATCGCGCCGAAGGCCGCCTGGTGGCCTACATGATCGACGTGTCCGA |

| | | | |
|---|---|---|---|
| | | | GCAGAAGCAGATCGAGCTGCAGCTGTCCCAGGCCCAGAAGATGCAGGCCATCGGCCAGCTGGCC<br>GGCGAAGTCGCGCACGACTTCAACAACCTCTTGACCGCCATCCAGCTGCGTCTGGACGAGTTGCT<br>GCATCGCCATCCCGTCGGTGATCCGTCGTACGAGGGTCTCAACGAGATCCGTCAGACGGGCGTGC<br>GCGCCGCCGACCTCGTGCGCAAGCTCTTGGCTTTCTCGCGCAAGCAGACCGTGCAGCGCGAGGT<br>GCTGGATCTGGGCGAGCTGATCAGCGAGTTCGAGGTCTTGCTGCGCCGCCTGCTGCGCGAAGAC<br>GTCAAGCTGATCACCGACTATGGCCGCGACCTGCCGCAGGTGCGCGCCGACAAGAGCCAGCTCG<br>AGACGGCGGTCATGAACCTGGCCGTCAACGCCCGCGACGCCGTGCGCGCGGCCAAGGGCGGCG<br>GCGTCGTGCGCATCCGCACCGCGCGCCTGACCCGCGACGAGGCGATCCAGCTGGGTTTCCCGGCC<br>GCCGACGGCGACACGGCCTTCATTGAGGTCAGTGACGATGGTCCGGGCATTCCGCCCGACGTCAT<br>GGGCAAGATCTTCGACCCGTTCTTCACCACCAAGCCGGTGGGCGAGGGTACGGGCCTAGGCCTA<br>GCCACGGTCTATGGCATCGTTAAGCAGAGCGACGGCTGGATTCACGTCCACAGCCGTCCGAACGA<br>AGGCGCGGCCTTCCGCATCTTCCTGCCGGTCTATGAAGCGCCCGCCGGCGCGGTCGCCGTCCAGG<br>CCGTCGCCGAGCCCGCCAAGCCGCGCGCCGCTCGCGACCTGTCGGGCGCCGGCCGCATCCTGTTC<br>GTCGAGGACGAGGACGCCGTGCGCAGCGTCGCCGCCCGCCTGCTGCGCGCCCGTGGCTACGAGG<br>TGCTTGAGGCGGCCGACGGCGAAGAGGCCCTGATCATCGCCGAGGAAAACGCCGGCACGATCGA<br>CCTCTTGATCAGCGACGTGATCATGCCCGGCATCGACGGCCCGACCTTGCTGAAGAAGGCGCGTG<br>GCTATCTGGGGACCGCGCCGGTGATGTTCATCTCCGGCTATGCCGAGGCCGAGTTCAGCGACCTCT<br>TGGAAGGCGAGACGGGCGTGACCTTCCTCCCCAAGCCGATCGACATCAAGACCCTGGCCGAGCG<br>CGTCAAGCAGCAGCTGCAGGCGGCGTAGcggagtgtat |
| **pGB1950** | pACYC-duet | pACYC-CpdRGFP-ChpT-CckAK- | gagatataccGTGGCCCGCATCCTCCTCGCCGAAGACGATGATTCCCTGCGCGGCTTCCTGGCCCGCG<br>CGCTGGAACGCGCCGGCTTCGAAGTCCAGGCCTGCGCCGACGGCGAAGAGGCCGTCCAGCACCT<br>GGACCATCCCTGGGACCTGCTGCTGACCGACATCGTCATGCCCGGCATGGACGGCATCGAGGTGG<br>CCCGCCAGGCCGCCGCCCGCGACCCGTCCCTGCGCATCATGTTCATCACCGGCTTCGCCGCCGTGG<br>CCCTCTCGGCCCAGGACCGCGCGCCCGCCGGCGCCAAGGTGCTGTCCAAGCCCGTGCACCTGCG<br>CGACCTCGTCGCCGAGGTCGAAAAGATGATGGCGGCCCTCGAGAGCGGCGGCGGCGGCAGTAA<br>AGGTGAAGAACTGTTCACCGGTGTTGTTCCGATCCTGGTTGAACTGGATGGTGATGTTAACGGCC<br>ACAAATTCTCTGTTCGTGGTGAAGGTGAAGGTGATGCAACCAACGGTAAACTGACCCTGAAATTC<br>ATCTGCACTACCGGTAAACTGCCGGTTCCATGGCCGACTCTGGTGACTACCCTGACCTATGGTGTTC<br>AGTGTTTTTCTCGTTACCCGGATCACATGAAGCAGCATGATTTCTTCAAATCTGCAATGCCGGAAGG<br>TTATGTACAGGAGCGCACCATTTCTTTCAAAGACGATGGCACCTACAAAACCCGTGCAGAGGTTAA<br>ATTTGAAGGTGATACTCTGGTGAACCGTATTGAACTGAAAGGCATTGATTTCAAAGAGGACGGCA<br>ACATCCTGGGCCACAAACTGGAATATAACTTCAACTCCCATAACGTTTACATCACCGCAGACAAACA<br>GAAGAACGGTATCAAAGCTAACTTCAAAATTCGCCATAACGTTGAAGACGGTAGCGTACAGCTGG<br>CGGACCACTACCAGCAGAACACTCCGATCGGTGATGGTCCGGTTCTGCTGCCGGATAACCACTACC<br>TGTCCACCCAGTCTAAACTGTCCAAAGACCCGAACGAAAAGCGCGACCACATGGTGCTGCTGGAG<br>TTCGTTACTGCAGCAGGTATCACGCACGGCATGGATGAACTCTACAAATGAgcggccgcat.........agat<br>atacatGTGACCGAGACCGTCACCGAGACCACCGCCCCCGCGTCCCCCGAAGCCGACGTCCAGGGT<br>CCCGATTTCGCCGCCATGCTGGCCGCGCGCCTGTGTCACGACTTCATCAGTCCGGCCAGCGCCATC<br>GTCTCGGGCCTGGATCTGCTGGAAGACCCCTCGGCCCAGGACATGCGCGACGACGCCATGAACCT<br>GATCGCCTCCTCGGCCCGCAAGCTGGCGGACCTCTTGCAGTTCACCCGAGTGGCCTTCGGCGCCT<br>CGGCCTCGGCCGAGAACTTCGACTCGCGCGAACTGGAAAAGCTGGCCCAGGGCGTCTTTGCCCA<br>TGTCCGCCCGACCCTGGACTGGCAGATCGAGCCGCAGGCGATGAACAAGCCCTCGTCGCGCGCG<br>GTGCTGAACATCGCCCAGATCGCCGCCAGCGCCCTGCCGGCCGGCGGCGTGGCCACCGTCAAGG<br>GCGTGGCCGCCGACGGGCGCTTCTCGATCATCGCCGACGCCAAGGGCCCGCGCGCGCGCCTGCG<br>TCCGGAGGTGCTGGCGGGCCTAAAGGGCGAGCCGCTGGCCGAGGGCCTGGGCGGTCCGTGGGT<br>GCAGGCGGCCTATCTGAACGCCCTGGTGCGCGCGGCCGGCGGCCAGATCGCCGTCGAGATCGGC<br>GAGGACCGCGCCTCGATCGCCGCCTGGGTCCCGGCGTAActcgagtctg.........tcagcgctagttatgcgac<br>tcctgcattaggaaattaatacgactcactataggggaattgtgagcggataacaattcccctgtagaaataattttgtttaacttt<br>aataaggagatataccATGGCCGACTTGCAGCTCCAGGACAAGGTTTCGACCGGCGCCCCGCGTCGGC<br>GGTTTGATCCATGGCTGGTCGGCGCGGCGGTGTTTTTCGTGGCTGCGGCGGCCCTCTCGGCGGCG |

| | | | |
|---|---|---|---|
| | | | CCGGCGCTCAAGGCCGGACCGACCACCCTGGCCGGCCTGCTGCTGCTTCTGGGCGTCGCGGGCG<br>TGGCTGTGTTGGGCCTTGTCGCCATTCGCGGCTCAGCGCTTTCCGGCGGCGACGCCGACCAGGCT<br>GAGGGGTTCATCGAGGCGCTGGCCGAGCCGGCCGCCCTGGCCGCCGCCGACGGTCGCGTCCTGG<br>CCGCCAACGGGCCCTGGCGCGAAGTCATGGGCGAACAGCGCCGCCTGCCCAAGGGCGTGGCGG<br>GCTCCAGCCTGTTTGCGGCCCTGGTCCAGGCGCGCCAGGGGCAGATGGCCGAGGGCATGCTGAG<br>CGCTGGAGGAACCGACTATACCGCCAAGGTCTCGCGCCTTGCGGGCGGACGGCTGATGATCCGG<br>CTTGCGCCCATCGTTGTCGCTGAGCCGGTTGTGGAAGACGCGTCGCCGGCCCCGGTCGCCGAGC<br>GCGCCGCCCCGCCACCCAGCTCGCTGGACGCCTTCGCCGGAGCCTCGCCGTTCGGCGCGGCCCT<br>GCTGGAAGGCCTGGAGCCGTTCACCTCGCGGGTGCTGGAGACCAATCCGGCGCTGACCACGATG<br>ACCGGCGCCAAGGCCGGTGTGCTGTTCGGCGATCTGATCGACGCCGCCTCGCGCGCCGAGGCCG<br>AGACGCGCCTGAACGAAGGCCGCGCCGGTCCCTACGAGGTGCGGTTGGCGCGCGATCCGTCGCG<br>CATCGCTCACCTTTACCTCTATCGCGCCGAAGGCCGCCTGGTGGCCTACATGATCGACGTGTCCGA<br>GCAGAAGCAGATCGAGCTGCAGCTGTCCCAGGCCCAGAAGATGCAGGCCATCGGCCAGCTGGCC<br>GGCGGCGTCGCGGCGGACTTCAACAACCTCTTGACCGCCATCCAGCTGCGTCTGGACGAGTTGCT<br>GCATCGCCATCCCGTCGGTGATCCGTCGTACGAGGGTCTCAACGAGATCCGTCAGACGGGCGTGC<br>GCGCCGCCGACCTCGTGCGCAAGCTCTTGGCTTTCTCGCGCAAGCAGACCGTGCAGCGCGAGGT<br>GCTGGATCTGGGCGAGCTGATCAGCGAGTTCGAGGTCTTGCTGCGCCGCCTGCTGCGCGAAGAC<br>GTCAAGCTGATCACCGACTATGGCCGCGACCTGCCGCAGGTGCGCGCCGACAAGAGCCAGCTCG<br>AGACGGCGGTCATGAACCTGGCCGTCAACGCCCGCGACGCCGTGCGCGCGGCCAAGGGCGGCG<br>GCGTCGTGCGCATCCGCACCGCGCGCCTGACCCGCGACGAGGCGATCCAGCTGGGTTTCCCGGCC<br>GCCGACGGCGACACGGCCTTCATTGAGGTCAGTGACGATGGTCCGGGCATTCCGCCCGACGTCAT<br>GGGCAAGATCTTCGACCCGTTCTTCACCACCAAGCCGGTGGGCGAGGGTACGGGCCTAGGCCTA<br>GCCACGGTCTATGGCATCGTTAAGCAGAGCGACGGCTGGATTCACGTCCACAGCCGTCCGAACGA<br>AGGCGCGGCCTTCCGCATCTTCCTGCCGGTCTATGAAGCGCCCGCCGGCGCGGTCGCCGTCCAGG<br>CCGTCGCCGAGCCCGCCAAGCCGCGCGCCGCTCGCGACCTGTCGGGCGCCGGCCGCATCCTGTTC<br>GTCGAGGACGAGGACGCCGTGCGCAGCGTCGCCGCCCGCCTGCTGCGCGCCCGTGGCTACGAGG<br>TGCTTGAGGCGGCCGACGGCGAAGAGGCCCTGATCATCGCCGAGGAAAACGCCGGCACGATCGA<br>CCTCTTGATCAGCGACGTGATCATGCCCGGCATCGACGGCCCGACCTTGCTGAAGAAGGCGCGTG<br>GCTATCTGGGGACCGCGCCGGTGATGTTCATCTCCGGCTATGCCGAGGCCGAGTTCAGCGACCTCT<br>TGGAAGGCGAGACGGGCGTGACCTTCCTCCCCAAGCCGATCGACATCAAGACCCTGGCCGAGCG<br>CGTCAAGCAGCAGCTGCAGGCGGCGTAGcggagtgtat |
| **pGB1951** | pACYC-duet | pACYC-CpdR$_{D51A}$ GFP-ChpT-CckA H+ | gagatataccATGGCCCGCATCCTCCTCGCCGAAGACGATGATTCCCTGCGCGGCTTCCTGGCCCGCG<br>CGCTGGAACGCGCCGGCTTCGAAGTCCAGGCCTGCGCCGACGGCGAAGAGGCCGTCCAGCACCT<br>GGACCATCCCTGGGACCTGCTGCTGACCGCCATCGTCATGCCCGGCATGGACGGCATCGAGGTGG<br>CCCGCCAGGCCGCCGCCCGCGACCCGTCCCTGCGCATCATGTTCATCACCGGCTTCGCCGCCGTGG<br>CCCTCTCGGCCCAGGACCGCGCGCCCGCCGGCGCCAAGGTGCTGTCCAAGCCCGTGCACCTGCG<br>CGACCTCGTCGCCGAGGTCGAAAAGATGATGGCGGCCCTCGAGAGCGGCGGCGGCGGCAGTAA<br>AGGTGAAGAACTGTTCACCGGTGTTGTTCCGATCCTGGTTGAACTGGATGGTGATGTTAACGGCC<br>ACAAATTCTCTGTTCGTGGTGAAGGTGAAGGTGATGCAACCAACGGTAAACTGACCCTGAAATTC<br>ATCTGCACTACCGGTAAACTGCCGGTTCCATGGCCGACTCTGGTGACTACCCTGACCTATGGTGTTC<br>AGTGTTTTTCTCGTTACCCGGATCACATGAAGCAGCATGATTTCTTCAAATCTGCAATGCCGGAAGG<br>TTATGTACAGGAGCGCACCATTTCTTTCAAAGACGATGGCACCTACAAAACCCGTGCAGAGGTTAA<br>ATTTGAAGGTGATACTCTGGTGAACCGTATTGAACTGAAAGGCATTGATTTCAAAGAGGACGGCA<br>ACATCCTGGGCCACAAACTGGAATATAACTTCAACTCCCATAACGTTTACATCACCGCAGACAAACA<br>GAAGAACGGTATCAAAGCTAACTTCAAAATTCGCCATAACGTTGAAGACGGTAGCGTACAGCTGG<br>CGGACCACTACCAGCAGAACACTCCGATCGGTGATGGTCCGGTTCTGCTGCCGGATAACCACTACC<br>TGTCCACCCAGTCTAAACTGTCCAAAGACCCGAACGAAAAGCGCGACCACATGGTGCTGCTGGAG<br>TTCGTTACTGCAGCAGGTATCACGCACGGCATGGATGAACTCTACAAATGAgcggccgcat.........agat<br>atacatGTGACCGAGACCGTCACCGAGACCACCGCCCCCGCGTCCCCCGAAGCCGACGTCCAGGGT<br>CCCGATTTCGCCGCCATGCTGGCCGCGCGCCTGTGTCACGACTTCATCAGTCCGGCCAGCGCCATC<br>GTCTCGGGCCTGGATCTGCTGGAAGACCCCTCGGCCCAGGACATGCGCGACGACGCCATGAACCT<br>GATCGCCTCCTCGGCCCGCAAGCTGGCGGACCTCTTGCAGTTCACCCGAGTGGCCTTCGGCGCCT<br>CGGCCTCGGCCGAGAACTTCGACTCGCGCGAACTGGAAAAGCTGGCCCAGGGCGTCTTTGCCCA<br>TGTCCGCCCGACCCTGGACTGGCAGATCGAGCCGCAGGCGATGAACAAGCCCTCGTCGCGCGCG |

| | | | |
|---|---|---|---|
| | | | GTGCTGAACATCGCCCAGATCGCCGCCAGCGCCCTGCCGGCCGGCGGCGTGGCCACCGTCAAGG<br>GCGTGGCCGCCGACGGGCGCTTCTCGATCATCGCCGACGCCAAGGGCCCGCGCGCGCGCCTGCG<br>TCCGGAGGTGCTGGCGGGCCTAAAGGGCGAGCCGCTGGCCGAGGGCCTGGGCGGTCCGTGGGT<br>GCAGGCGGCCTATCTGAACGCCCTGGTGCGCGCGGCCGGCGGCCAGATCGCCGTCGAGATCGGC<br>GAGGACCGCGCCTCGATCGCCGCCTGGGTCCCGGCGTAActcgagtctg.........tcagcgctagttatgcgac<br>tcctgcattaggaaattaatacgactcactataggggaattgtgagcggataacaattcccctgtagaaataattttgtttaacttt<br>aataaggagatataccATGGCCGACTTGCAGCTCCAGGACAAGGTTTCGACCGGCGCCCCGCGTCGGC<br>GGTTTGATCCATGGCTGGTCGGCGCGGCGGTGTTTTTCGTGGCTGCGGCGGCCCTCTCGGCGGCG<br>CCGGCGCTCAAGGCCGGACCGACCACCCTGGCCGGCCTGCTGCTGCTTCTGGGCGTCGCGGGCG<br>TGGCTGTGTTGGGCCTTGTCGCCATTCGCGGCTCAGCGCTTTCCGGCGGCGACGCCGACCAGGCT<br>GAGGGGTTCATCGAGGCGCTGGCCGAGCCGGCCGCCCTGGCCGCCGCCGACGGTCGCGTCCTGG<br>CCGCCAACGGGCCCTGGCGCGAAGTCATGGGCGAACAGCGCCGCCTGCCCAAGGGCGTGGCGG<br>GCTCCAGCCTGTTTGCGGCCCTGGTCCAGGCGCGCCAGGGGCAGATGGCCGAGGGCATGCTGAG<br>CGCTGGAGGAACCGACTATACCGCCAAGGTCTCGCGCCTTGCGGGCGGACGGCTGATGATCCGG<br>CTTGCGCCCATCGTTGTCGCTGAGCCGGTTGTGGAAGACGCGTCGCCGGCCCCGGTCGCCGAGC<br>GCGCCGCCCCGCCACCCAGCTCGCTGGACGCCTTCGCCGGAGCCTCGCCGTTCGGCGCGGCCCT<br>GCTGGAAGGCCTGGAGCCGTTCACCTCGCGGGTGCTGGAGACCAATCCGGCGCTGACCACGATG<br>ACCGGCGCCAAGGCCGGTGTGCTGTTCGGCGATCTGATCGACGCCGCCTCGCGCGCCGAGGCCG<br>AGACGCGCCTGAACGAAGGCCGCGCCGGTCCCTACGAGGTGCGGTTGGCGCGCGATCCGTCGCG<br>CATCGCTCACCTTTACCTCTATCGCGCCGAAGGCCGCCTGGTGGCCTACATGATCGACGTGTCCGA<br>GCAGAAGCAGATCGAGCTGCAGCTGTCCCAGGCCCAGAAGATGCAGGCCATCGGCCAGCTGGCC<br>GGCGAAGTCGCGCACGACTTCAACAACCTCTTGACCGCCATCCAGCTGCGTCTGGACGAGTTGCT<br>GCATCGCCATCCCGTCGGTGATCCGTCGTACGAGGGTCTCAACGAGATCCGTCAGACGGGCGTGC<br>GCGCCGCCGACCTCGTGCGCAAGCTCTTGGCTTTCTCGCGCAAGCAGACCGTGCAGCGCGAGGT<br>GCTGGATCTGGGCGAGCTGATCAGCGAGTTCGAGGTCTTGCTGCGCCGCCTGCTGCGCGAAGAC<br>GTCAAGCTGATCACCGACTATGGCCGCGACCTGCCGCAGGTGCGCGCCGACAAGAGCCAGCTCG<br>AGACGGCGGTCATGAACCTGGCCGTCAACGCCCGCGACGCCGTGCGCGCGGCCAAGGGCGGCG<br>GCGTCGTGCGCATCCGCACCGCGCGCCTGACCCGCGACGAGGCGATCCAGCTGGGTTTCCCGGCC<br>GCCGACGGCGACACGGCCTTCATTGAGGTCAGTGACGATGGTCCGGGCATTCCGCCCGACGTCAT<br>GGGCAAGATCTTCGACCCGTTCTTCACCACCAAGCCGGTGGGCGAGGGTACGGGCCTAGGCCTA<br>GCCACGGTCTATGGCATCGTTAAGCAGAGCGACGGCTGGATTCACGTCCACAGCCGTCCGAACGA<br>AGGCGCGGCCTTCCGCATCTTCCTGCCGGTCTATGAAGCGCCCGCCGGCGCGGTCGCCGTCCAGG<br>CCGTCGCCGAGCCCGCCAAGCCGCGCGCCGCTCGCGACCTGTCGGGCGCCGGCCGCATCCTGTTC<br>GTCGAGGACGAGGACGCCGTGCGCAGCGTCGCCGCCCGCCTGCTGCGCGCCCGTGGCTACGAGG<br>TGCTTGAGGCGGCCGACGGCGAAGAGGCCCTGATCATCGCCGAGGAAAACGCCGGCACGATCGA<br>CCTCTTGATCAGCGACGTGATCATGCCCGGCATCGACGGCCCGACCTTGCTGAAGAAGGCGCGTG<br>GCTATCTGGGGACCGCGCCGGTGATGTTCATCTCCGGCTATGCCGAGGCCGAGTTCAGCGACCTCT<br>TGGAAGGCGAGACGGGCGTGACCTTCCTCCCCAAGCCGATCGACATCAAGACCCTGGCCGAGCG<br>CGTCAAGCAGCAGCTGCAGGCGGCGTAGcggagtgtat |
| **pGB1952** | pACYC-duet | pACYC-CpdRGFP-ChpT | gagatataccGTGGCCCGCATCCTCCTCGCCGAAGACGATGATTCCCTGCGCGGCTTCCTGGCCCGCG<br>CGCTGGAACGCGCCGGCTTCGAAGTCCAGGCCTGCGCCGACGGCGAAGAGGCCGTCCAGCACCT<br>GGACCATCCCTGGGACCTGCTGCTGACCGACATCGTCATGCCCGGCATGGACGGCATCGAGGTGG<br>CCCGCCAGGCCGCCGCCCGCGACCCGTCCCTGCGCATCATGTTCATCACCGGCTTCGCCGCCGTGG<br>CCCTCTCGGCCCAGGACCGCGCGCCCGCCGGCGCCAAGGTGCTGTCCAAGCCCGTGCACCTGCG<br>CGACCTCGTCGCCGAGGTCGAAAAGATGATGGCGGCCCTCGAGAGCGGCGGCGGCGGCAGTAA<br>AGGTGAAGAACTGTTCACCGGTGTTGTTCCGATCCTGGTTGAACTGGATGGTGATGTTAACGGCC<br>ACAAATTCTCTGTTCGTGGTGAAGGTGAAGGTGATGCAACCAACGGTAAACTGACCCTGAAATTC<br>ATCTGCACTACCGGTAAACTGCCGGTTCCATGGCCGACTCTGGTGACTACCCTGACCTATGGTGTTC<br>AGTGTTTTTCTCGTTACCCGGATCACATGAAGCAGCATGATTTCTTCAAATCTGCAATGCCGGAAGG<br>TTATGTACAGGAGCGCACCATTTCTTTCAAAGACGATGGCACCTACAAAACCCGTGCAGAGGTTAA<br>ATTTGAAGGTGATACTCTGGTGAACCGTATTGAACTGAAAGGCATTGATTTCAAAGAGGACGGCA<br>ACATCCTGGGCCACAAACTGGAATATAACTTCAACTCCCATAACGTTTACATCACCGCAGACAAACA<br>GAAGAACGGTATCAAAGCTAACTTCAAAATTCGCCATAACGTTGAAGACGGTAGCGTACAGCTGG<br>CGGACCACTACCAGCAGAACACTCCGATCGGTGATGGTCCGGTTCTGCTGCCGGATAACCACTACC |

| | | | |
|---|---|---|---|
| | | | TGTCCACCCAGTCTAAACTGTCCAAAGACCCGAACGAAAAGCGCGACCACATGGTGCTGCTGGAG<br>TTCGTTACTGCAGCAGGTATCACGCACGGCATGGATGAACTCTACAAATGAgcggccgcat.........agat<br>atacatGTGACCGAGACCGTCACCGAGACCACCGCCCCCGCGTCCCCGAAGCCGACGTCCAGGGT<br>CCCGATTTCGCCGCCATGCTGGCCGCGCGCCTGTGTCACGACTTCATCAGTCCGGCCAGCGCCATC<br>GTCTCGGGCCTGGATCTGCTGGAAGACCCCTCGGCCCAGGACATGCGCGACGACGCCATGAACCT<br>GATCGCCTCCTCGGCCCGCAAGCTGGCGGACCTCTTGCAGTTCACCCGAGTGGCCTTCGGCGCCT<br>CGGCCTCGGCCGAGAACTTCGACTCGCGCGAACTGGAAAAGCTGGCCCAGGGCGTCTTTGCCCA<br>TGTCCGCCCGACCCTGGACTGGCAGATCGAGCCGCAGGCGATGAACAAGCCCTCGTCGCGCGCG<br>GTGCTGAACATCGCCCAGATCGCCGCCAGCGCCCTGCCGGCCGGCGGCGTGGCCACCGTCAAGG<br>GCGTGGCCGCCGACGGGCGCTTCTCGATCATCGCCGACGCCAAGGGCCCGCGCGCGCGCCTGCG<br>TCCGGAGGTGCTGGCGGGCCTAAAGGGCGAGCCGCTGGCCGAGGGCCTGGGCGGTCCGTGGGT<br>GCAGGCGGCCTATCTGAACGCCCTGGTGCGCGCGGCCGGCGGCCAGATCGCCGTCGAGATCGGC<br>GAGGACCGCGCCTCGATCGCCGCCTGGGTCCCGGCGTAActcgagtctg |
| **pGB1953** | pACYC-duet | pACYC-CpdRGFP | gagatataccGTGGCCCGCATCCTCCTCGCCGAAGACGATGATTCCCTGCGCGGCTTCCTGGCCCGCG<br>CGCTGGAACGCGCCGGCTTCGAAGTCCAGGCCTGCGCCGACGGCGAAGAGGCCGTCCAGCACCT<br>GGACCATCCCTGGGACCTGCTGCTGACCGACATCGTCATGCCCGGCATGGACGGCATCGAGGTGG<br>CCCGCCAGGCCGCCGCCCGCGACCCGTCCCTGCGCATCATGTTCATCACCGGCTTCGCCGCCGTGG<br>CCCTCTCGGCCCAGGACCGCGCGCCCGCCGGCGCCAAGGTGCTGTCCAAGCCCGTGCACCTGCG<br>CGACCTCGTCGCCGAGGTCGAAAAGATGATGGCGGCCCTCGAGAGCGGCGGCGGCGGCAGTAA<br>AGGTGAAGAACTGTTCACCGGTGTTGTTCCGATCCTGGTTGAACTGGATGGTGATGTTAACGGCC<br>ACAAATTCTCTGTTCGTGGTGAAGGTGAAGGTGATGCAACCAACGGTAAACTGACCCTGAAATTC<br>ATCTGCACTACCGGTAAACTGCCGGTTCCATGGCCGACTCTGGTGACTACCCTGACCTATGGTGTTC<br>AGTGTTTTTCTCGTTACCCGGATCACATGAAGCAGCATGATTTCTTCAAATCTGCAATGCCGGAAGG<br>TTATGTACAGGAGCGCACCATTTCTTTCAAAGACGATGGCACCTACAAAACCCGTGCAGAGGTTAA<br>ATTTGAAGGTGATACTCTGGTGAACCGTATTGAACTGAAAGGCATTGATTTCAAAGAGGACGGCA<br>ACATCCTGGGCCACAAACTGGAATATAACTTCAACTCCCATAACGTTTACATCACCGCAGACAAACA<br>GAAGAACGGTATCAAAGCTAACTTCAAAATTCGCCATAACGTTGAAGACGGTAGCGTACAGCTGG<br>CGGACCACTACCAGCAGAACACTCCGATCGGTGATGGTCCGGTTCTGCTGCCGGATAACCACTACC<br>TGTCCACCCAGTCTAAACTGTCCAAAGACCCGAACGAAAAGCGCGACCACATGGTGCTGCTGGAG<br>TTCGTTACTGCAGCAGGTATCACGCACGGCATGGATGAACTCTACAAATGAgcggccgcat |
| **pGB1954** | pBAD | pBAD-mCherry ΔN term PopZ | ggaattaaccATGGTGAGCAAGGGCGAGGAGGATAACATGGCCATCATCAAGGAGTTCATGCGCTTC<br>AAGGTGCACATGGAGGGCTCCGTGAACGGCCACGAGTTCGAGATCGAGGGCGAGGGCGAGGGC<br>CGCCCCTACGAGGGCACCCAGACCGCCAAGCTGAAGGTGACCAAGGGTGGCCCCCTGCCCTTCG<br>CCTGGGACATCCTGTCCCCTCAGTTCATGTACGGCTCCAAGGCCTACGTGAAGCACCCCGCCGACA<br>TCCCCGACTACTTGAAGCTGTCCTTCCCCGAGGGCTTCAAGTGGGAGCGCGTGATGAACTTCGAG<br>GACGGCGGCGTGGTGACCGTGACCCAGGACTCCTCCCTGCAGGACGGCGAGTTCATCTACAAGG<br>TGAAGCTGCGCGGCACCAACTTCCCCTCCGACGGCCCCGTAATGCAGAAGAAGACCATGGGCTG<br>GGAGGCCTCCTCCGAGCGGATGTACCCCGAGGACGGCGCCCTGAAGGGCGAGATCAAGCAGAG<br>GCTGAAGCTGAAGGACGGCGGCCACTACGACGCTGAGGTCAAGACCACCTACAAGGCCAAGAA<br>GCCCGTGCAGCTGCCCGGCGCCTACAACGTCAACATCAAGTTGGACATCACCTCCCACAACGAGG<br>ACTACACCATCGTGGAACAGTACGAACGCGCCGAGGGCCGCCACTCCACCGGCGGCATGGACGA<br>GCTGTACAAGcctgcaggcgccttaattaatatgcatggtaccGATGACGCGCCGGCGGAGCCTGCGGCCGA<br>AGCGGCGCCCCCGCCGCCGCCGGAACCCGAACCTGAACCGGTGTCGTTCGACGACGAGGTTCTG<br>GAATTGACGGATCCGATCGCGCCCGAGCCCGAGCTGCCGCCGCTGGAGACTGTCGGCGACATCG<br>ACGTCTATTCGCCGCCGGAACCTGAGTCGGAACCGGCCTACACGCCGCCGCCGGCGGCTCCGGTG<br>TTTGATCGCGACGAAGTCGCCGAGCAGCTGGTCGGCGTTTCGGCCGCTTCGGCCGCGGCGAGCG<br>CCTTCGGCAGCCTGAGCTCGGCCCTGCTGATGCCCAAGGACGGTCGGACGCTGGAAGACGTCGT<br>ACGCGAGCTGCTGCGCCCGCTGCTCAAGGAGTGGCTGGACCAGAACCTGCCGCGCATCGTCGAG<br>ACCAAGGTTGAGGAAGAAGTGCAGCGTATCTCTCGGGGACGCGGCGCCTAActcgagatct |

| | | | |
|---|---|---|---|
| **pGB1955** | pMCS5 (NdeI, KpnI) | PMCS5- pCpdR CpdR YFP | ccaattgcatGGTCGCCGTCGAACATCGCCGGGTCCAGCTCCCAGGGCTCACGGTTCAGATCCACATA<br>GGCGCGGGCGAAGCGGGCGCGCACCACGGCGGCCCCCAGGGCGGGCGCGCCGCCGATGATGC<br>GGTCGACAAACGCGTCCTCGGACGCGCGCAGGGTCTCGAGCGGCAGGCGGACCGCGGCGACCA<br>TGTCCTCCGGATAGAGGTCGCCCGAGTGCGGCGAGGCGAACACCAAGGCGGTCGGCGGCGGCG<br>CCCCGGCCGGGACCGCGCGCAGCACCTCGAACGCTGGCCCGCCAAAGGTCTCCAGCGGCGGGG<br>ACGAAAGCTCTTCGCTCGAAAGGGGCCGCCACGCGCTCATGCCGCATCATCGAAGCGTCGCGGCG<br>CGGGGTCAAATCTCAATCGCCGCTGACGAAACATCCCCAGCCGCGACGTTTAGGTTCATCCCCGAT<br>TTACGGACGGGGCGATAAGGTGGATCCTCTATCGACGATCTTAATCGGACACGTGACCCCATGGCC<br>CGCATCCTCCTCGCCGAAGACGATGATTCCCTGCGCGGCTTCCTGGCCCGCGCGCTGGAACGCGC<br>CGGCTTCGAAGTCCAGGCCTGCGCCGACGGCGAAGAGGCCGTCCAGCACCTGGACCATCCCTGG<br>GACCTGCTGCTGACCGACATCGTCATGCCCGGCATGGACGGCATCGAGGTGGCCCGCCAGGCCGC<br>CGCCCGCGACCCGTCCCTGCGCATCATGTTCATCACCGGCTTCGCCGCCGTGGCCCTCTCGGCCCA<br>GGACCGCGCGCCCGCCGGCGCCAAGGTGCTGTCCAAGCCCGTGCACCTGCGCGACCTCGTCGCC<br>GAGGTCGAAAAGATGATGGCGGCCCGGTACCGCGGGCCCGGGATCCACATGGTGAGCAAGGGC<br>GAGGAGCTGTTCACCGGGGTGGTGCCCATCCTGGTCGAGCTGGACGGCGACGTAAACGGCCACA<br>AGTTCAGCGTGTCCGGCGAGGGCGAGGGCGATGCCACCTACGGCAAGCTGACCCTGAAGTTCAT<br>CTGCACCACCGGCAAGCTGCCCGTGCCCTGGCCCACCCTCGTGACCACCTTCGGCTACGGCCTGC<br>AGTGCTTCGCCCGCTACCCCGACCACATGAAGCAGCACGACTTCTTCAAGTCCGCCATGCCCGAAG<br>GCTACGTCCAGGAGCGCACCATCTTCTTCAAGGACGACGGCAACTACAAGACCCGCGCCGAGGT<br>GAAGTTCGAGGGCGACACCCTGGTGAACCGCATCGAGCTGAAGGGCATCGACTTCAAGGAGGAC<br>GGCAACATCCTGGGGCACAAGCTGGAGTACAACTACAACAGCCACAACGTCTATATCATGGCCGA<br>CAAGCAGAAGAACGGCATCAAGGTGAACTTCAAGATCCGCCACAACATCGAGGACGGCAGCGTG<br>CAGCTCGCCGACCACTACCAGCAGAACACCCCCATCGGCGACGGCCCCGTGCTGCTGCCCGACAA<br>CCACTACCTGAGCTACCAGTCCGCCCTGAGCAAAGACCCCAACGAGAAGCGCGATCACATGGTCC<br>TGCTGGAGTTCGTGACCGCCGCCGGGATCACTCTCGGCATGGACGAGCTGTACAAGTAGggtacctt<br>aa |
| **pGB1956** | pMCS5 | PMCS5- pCpdR CpdR$_{D51A}$ YFP | ccaattgcatGGTCGCCGTCGAACATCGCCGGGTCCAGCTCCCAGGGCTCACGGTTCAGATCCACATA<br>GGCGCGGGCGAAGCGGGCGCGCACCACGGCGGCCCCCAGGGCGGGCGCGCCGCCGATGATGC<br>GGTCGACAAACGCGTCCTCGGACGCGCGCAGGGTCTCGAGCGGCAGGCGGACCGCGGCGACCA<br>TGTCCTCCGGATAGAGGTCGCCCGAGTGCGGCGAGGCGAACACCAAGGCGGTCGGCGGCGGCG<br>CCCCGGCCGGGACCGCGCGCAGCACCTCGAACGCTGGCCCGCCAAAGGTCTCCAGCGGCGGGG<br>ACGAAAGCTCTTCGCTCGAAAGGGGCCGCCACGCGCTCATGCCGCATCATCGAAGCGTCGCGGCG<br>CGGGGTCAAATCTCAATCGCCGCTGACGAAACATCCCCAGCCGCGACGTTTAGGTTCATCCCCGAT<br>TTACGGACGGGGCGATAAGGTGGATCCTCTATCGACGATCTTAATCGGACACGTGACCCCATGGCC<br>CGCATCCTCCTCGCCGAAGACGATGATTCCCTGCGCGGCTTCCTGGCCCGCGCGCTGGAACGCGC<br>CGGCTTCGAAGTCCAGGCCTGCGCCGACGGCGAAGAGGCCGTCCAGCACCTGGACCATCCCTGG<br>GACCTGCTGCTGACCGCCATCGTCATGCCCGGCATGGACGGCATCGAGGTGGCCCGCCAGGCCGC<br>CGCCCGCGACCCGTCCCTGCGCATCATGTTCATCACCGGCTTCGCCGCCGTGGCCCTCTCGGCCCA<br>GGACCGCGCGCCCGCCGGCGCCAAGGTGCTGTCCAAGCCCGTGCACCTGCGCGACCTCGTCGCC<br>GAGGTCGAAAAGATGATGGCGGCCCGGTACCGCGGGCCCGGGATCCACATGGTGAGCAAGGGC<br>GAGGAGCTGTTCACCGGGGTGGTGCCCATCCTGGTCGAGCTGGACGGCGACGTAAACGGCCACA<br>AGTTCAGCGTGTCCGGCGAGGGCGAGGGCGATGCCACCTACGGCAAGCTGACCCTGAAGTTCAT<br>CTGCACCACCGGCAAGCTGCCCGTGCCCTGGCCCACCCTCGTGACCACCTTCGGCTACGGCCTGC<br>AGTGCTTCGCCCGCTACCCCGACCACATGAAGCAGCACGACTTCTTCAAGTCCGCCATGCCCGAAG<br>GCTACGTCCAGGAGCGCACCATCTTCTTCAAGGACGACGGCAACTACAAGACCCGCGCCGAGGT<br>GAAGTTCGAGGGCGACACCCTGGTGAACCGCATCGAGCTGAAGGGCATCGACTTCAAGGAGGAC<br>GGCAACATCCTGGGGCACAAGCTGGAGTACAACTACAACAGCCACAACGTCTATATCATGGCCGA<br>CAAGCAGAAGAACGGCATCAAGGTGAACTTCAAGATCCGCCACAACATCGAGGACGGCAGCGTG<br>CAGCTCGCCGACCACTACCAGCAGAACACCCCCATCGGCGACGGCCCCGTGCTGCTGCCCGACAA<br>CCACTACCTGAGCTACCAGTCCGCCCTGAGCAAAGACCCCAACGAGAAGCGCGATCACATGGTCC<br>TGCTGGAGTTCGTGACCGCCGCCGGGATCACTCTCGGCATGGACGAGCTGTACAAGTAGggtacctt<br>aa |

| | | | |
|---|---|---|---|
| **pGB1959** | pBXMCS2 (NdeI, EcoRI) | pBXMCS2-CckA K+ | agacgaccatATGGCCGACTTGCAGCTCCAGGACAAGGTTTCGACCGGCGCCCCGCGTCGGCGGTTT<br>GATCCATGGCTGGTCGGCGCGGCGGTGTTTTTCGTGGCTGCGGCGGCCCTCTCGGCGGCGCCGG<br>CGCTCAAGGCCGGACCGACCACCCTGGCCGGCCTGCTGCTGCTTCTGGGCGTCGCGGGCGTGGC<br>TGTGTTGGGCCTTGTCGCCATTCGCGGCTCAGCGCTTTCCGGCGGCGACGCCGACCAGGCTGAG<br>GGGTTCATCGAGGCGCTGGCCGAGCCGGCCGCCCTGGCCGCCGCCGACGGTCGCGTCCTGGCCG<br>CCAACGGGCCCTGGCGCGAAGTCATGGGCGAACAGCGCCGCCTGCCCAAGGGCGTGGCGGGCT<br>CCAGCCTGTTTGCGGCCCTGGTCCAGGCGCGCCAGGGGCAGATGGCCGAGGGCATGCTGAGCGC<br>TGGAGGAACCGACTATACCGCCAAGGTCTCGCGCCTTGCGGGCGGACGGCTGATGATCCGGCTTG<br>CGCCCATCGTTGTCGCTGAGCCGGTTGTGGAAGACGCGTCGCCGGCCCCGGTCGCCGAGCGCGC<br>CGCCCCGCCACCCAGCTCGCTGGACGCCTTCGCCGGAGCCTCGCCGTTCGGCGCGGCCCTGCTGG<br>AAGGCCTGGAGCCGTTCACCTCGCGGGTGCTGGAGACCAATCCGGCGCTGACCACGATGACCGG<br>CGCCAAGGCCGGTGTGCTGTTCGGCGATCTGATCGACGCCGCCTCGCGCGCCGAGGCCGAGACG<br>CGCCTGAACGAAGGCCGCGCCGGTCCCTACGAGGTGCGGTTGGCGCGCGATCCGTCGCGCATCG<br>CTCACCTTTACCTCTATCGCGCCGAAGGCCGCCTGGTGGCCTACATGATCGACGTGTCCGAGCAGA<br>AGCAGATCGAGCTGCAGCTGTCCCAGGCCCAGAAGATGCAGGCCATCGGCCAGCTGGCCGGCGA<br>AGTCGCGCACGACTTCAACAACCTCTTGACCGCCATCCAGCTGCGTCTGGACGAGTTGCTGCATCG<br>CCATCCCGTCGGTGATCCGTCGTACGAGGGTCTCAACGAGATCCGTCAGACGGGCGTGCGCGCCG<br>CCGACCTCGTGCGCAAGCTCTTGGCTTTCTCGCGCAAGCAGACCGTGCAGCGCGAGGTGCTGGAT<br>CTGGGCGAGCTGATCAGCGAGTTCGAGGTCTTGCTGCGCCGCCTGCTGCGCGAAGACGTCAAGC<br>TGATCACCGACTATGGCCGCGACCTGCCGCAGGTGCGCGCCGACAAGAGCCAGCTCGAGACGGC<br>GGTCATGAACCTGGCCGTCAACGCCCGCGACGCCGTGCGCGCGGCCAAGGGCGGCGGCGTCGT<br>GCGCATCCGCACCGCGCGCCTGACCCGCGACGAGGCGATCCAGCTGGGTTTCCCGGCCGCCGAC<br>GGCGACACGGCCTTCATTGAGGTCAGTGACGATGGTCCGGGCATTCCGCCCGACGTCATGGGCAA<br>GATCTTCGACCCGTTCTTCACCACCAAGCCGGTGGGCGAGGGTACGGGCCTAGGCCTAGCCACGG<br>TCTATGGCATCGTTAAGCAGAGCGACGGCTGGATTCACGTCCACAGCCGTCCGAACGAAGGCGCG<br>GCCTTCCGCATCTTCCTGCCGGTCTATGAAGCGCCCGCCGGCGCGGTCGCCGTCCAGGCCGTCGC<br>CGAGCCCGCCAAGCCGCGCGCCGCTCGCGACCTGTCGGGCGCCGGCCGCATCCTGTTCGTCGAG<br>GACGAGGACGCCGTGCGCAGCGTCGCCGCCCGCCTGCTGCGCGCCCGTGGCTACGAGGTGCTTG<br>AGGCGGCCGACGGCGAAGAGGCCCTGATCATCGCCGAGGAAAACGCCGGCACGATCGACCTCTT<br>GATCAGCGACGTGATCATGCCCGGCATCGACGGCCCGACCTTGCTGAAGAAGGCGCGTGGCTATC<br>TGGGGACCGCGCCGGTGATGTTCATCTCCGGCTATGCCGAGGCCGAGTTCAGCGACCTCTTGGAA<br>GGCGAGACGGGCGTGACCTTCCTCCCCAAGCCGATCGACATCAAGACCCTGGCCGAGCGCGGTC<br>AAGCAGCAGCTGCAAGCAGCATAGgaattcctgc |
| **pGB1960** | pBXMCS2 | pBXMCS2-CckA H- | agacgaccatATGGCCGACTTGCAGCTCCAGGACAAGGTTTCGACCGGCGCCCCGCGTCGGCGGTTT<br>GATCCATGGCTGGTCGGCGCGGCGGTGTTTTTCGTGGCTGCGGCGGCCCTCTCGGCGGCGCCGG<br>CGCTCAAGGCCGGACCGACCACCCTGGCCGGCCTGCTGCTGCTTCTGGGCGTCGCGGGCGTGGC<br>TGTGTTGGGCCTTGTCGCCATTCGCGGCTCAGCGCTTTCCGGCGGCGACGCCGACCAGGCTGAG<br>GGGTTCATCGAGGCGCTGGCCGAGCCGGCCGCCCTGGCCGCCGCCGACGGTCGCGTCCTGGCCG<br>CCAACGGGCCCTGGCGCGAAGTCATGGGCGAACAGCGCCGCCTGCCCAAGGGCGTGGCGGGCT<br>CCAGCCTGTTTGCGGCCCTGGTCCAGGCGCGCCAGGGGCAGATGGCCGAGGGCATGCTGAGCGC<br>TGGAGGAACCGACTATACCGCCAAGGTCTCGCGCCTTGCGGGCGGACGGCTGATGATCCGGCTTG<br>CGCCCATCGTTGTCGCTGAGCCGGTTGTGGAAGACGCGTCGCCGGCCCCGGTCGCCGAGCGCGC<br>CGCCCCGCCACCCAGCTCGCTGGACGCCTTCGCCGGAGCCTCGCCGTTCGGCGCGGCCCTGCTGG<br>AAGGCCTGGAGCCGTTCACCTCGCGGGTGCTGGAGACCAATCCGGCGCTGACCACGATGACCGG<br>CGCCAAGGCCGGTGTGCTGTTCGGCGATCTGATCGACGCCGCCTCGCGCGCCGAGGCCGAGACG<br>CGCCTGAACGAAGGCCGCGCCGGTCCCTACGAGGTGCGGTTGGCGCGCGATCCGTCGCGCATCG<br>CTCACCTTTACCTCTATCGCGCCGAAGGCCGCCTGGTGGCCTACATGATCGACGTGTCCGAGCAGA<br>AGCAGATCGAGCTGCAGCTGTCCCAGGCCCAGAAGATGCAGGCCATCGGCCAGCTGGCCGGCGG<br>CGTCGCGGCGGACTTCAACAACCTCTTGACCGCCATCCAGCTGCGTCTGGACGAGTTGCTGCATC<br>GCCATCCCGTCGGTGATCCGTCGTACGAGGGTCTCAACGAGATCCGTCAGACGGGCGTGCGCGCC<br>GCCGACCTCGTGCGCAAGCTCTTGGCTTTCTCGCGCAAGCAGACCGTGCAGCGCGAGGTGCTGG<br>ATCTGGGCGAGCTGATCAGCGAGTTCGAGGTCTTGCTGCGCCGCCTGCTGCGCGAAGACGTCAA<br>GCTGATCACCGACTATGGCCGCGACCTGCCGCAGGTGCGCGCCGACAAGAGCCAGCTCGAGACG<br>GCGGTCATGAACCTGGCCGTCAACGCCCGCGACGCCGTGCGCGCGGCCAAGGGCGGCGGCGTC |

| | | | |
|---|---|---|---|
| | | | GTGCGCATCCGCACCGCGCGCCTGACCCGCGACGAGGCGATCCAGCTGGGTTTCCCGGCCGCCG<br>ACGGCGACACGGCCTTCATTGAGGTCAGTGACGATGGTCCGGGCATTCCGCCCGACGTCATGGGC<br>AAGATCTTCGACCCGTTCTTCACCACCAAGCCGGTGGGCGAGGGTACGGGCCTAGGCCTAGCCAC<br>GGTCTATGGCATCGTTAAGCAGAGCGACGGCTGGATTCACGTCCACAGCCGTCCGAACGAAGGCG<br>CGGCCTTCCGCATCTTCCTGCCGGTCTATGAAGCGCCCGCCGGCGCGGTCGCCGTCCAGGCCGTC<br>GCCGAGCCCGCCAAGCCGCGCGCCGCTCGCGACCTGTCGGGCGCCGGCCGCATCCTGTTCGTCG<br>AGGACGAGGACGCCGTGCGCAGCGTCGCCGCCCGCCTGCTGCGCGCCCGTGGCTACGAGGTGCT<br>TGAGGCGGCCGACGGCGAAGAGGCCCTGATCATCGCCGAGGAAAACGCCGGCACGATCGACCTC<br>TTGATCAGCGACGTGATCATGCCCGGCATCGACGGCCCGACCTTGCTGAAGAAGGCGCGTGGCTA<br>TCTGGGGACCGCGCCGGTGATGTTCATCTCCGGCTATGCCGAGGCCGAGTTCAGCGACCTCTTGG<br>AAGGCGAGACGGGCGTGACCTTCCTCCCCAAGCCGATCGACATCAAGACCCTGGCCGAGCGCGT<br>CAAGCAGCAGCTGCAAGCAGCATAGgaattcctgc |
| **pGB1961** | pXMCS5 (NdeI, KpnI) | pXMCS5-YFP-CtrA RD+15 | agacgaccatATGAGCAAGGGCGAGGAGCTGTTCACCGGGGTGGTGCCCATCCTGGTCGAGCTGGA<br>CGGCGACGTAAACGGCCACAAGTTCAGCGTGTCCGGCGAGGGCGAGGGCGATGCCACCTACGGC<br>AAGCTGACCCTGAAGTTCATCTGCACCACCGGCAAGCTGCCCGTGCCCTGGCCCACCCTCGTGAC<br>CACCTTCGGCTACGGCCTGCAGTGCTTCGCCCGCTACCCCGACCACATGAAGCAGCACGACTTCTT<br>CAAGTCCGCCATGCCCGAAGGCTACGTCCAGGAGCGCACCATCTTCTTCAAGGACGACGGCAACT<br>ACAAGACCCGCGCCGAGGTGAAGTTCGAGGGCGACACCCTGGTGAACCGCATCGAGCTGAAGG<br>GCATCGACTTCAAGGAGGACGGCAACATCCTGGGGCACAAGCTGGAGTACAACTACAACAGCCA<br>CAACGTCTATATCATGGCCGACAAGCAGAAGAACGGCATCAAGGTGAACTTCAAGATCCGCCACA<br>ACATCGAGGACGGCAGCGTGCAGCTCGCCGACCACTACCAGCAGAACACCCCCATCGGCGACGG<br>CCCCGTGCTGCTGCCCGACAACCACTACCTGAGCTACCAGTCCGCCCTGAGCAAAGACCCCAACG<br>AGAAGCGCGATCACATGGTCCTGCTGGAGTTCGTGACCGCCGCCGGGATCACTCTCGGCATGGAC<br>GAGCTGTACAAGggaagtagcggtGCgCGCGTACTGTTGATCGAGGATGACAGCGCGACGGCGCAGA<br>CCATCGAACTGATGCTGAAGTCTGAAGGCTTCAACGTCTATACGACGGATCTGGGTGAAGAAGGC<br>GTCGATCTGGGCAAGATCTACGACTACGATCTTATCCTGCTCGACCTCAATCTTCCGGACATGAGCG<br>GCATCGATGTTCTGCGCACCCTGCGGGTCGCGAAGATCAACACGCCCATCATGATCCTGTCGGGCT<br>CGTCGGAAATCGACACCAAGGTCAAGACCTTCGCCGGCGGCGCCGACGACTACATGACCAAGCC<br>GTTCCACAAGGACGAAATGATCGCCCGCATCCACGCGGTGGTCCGTGGCTATGTGCTGCGCGACC<br>CGAACGAGCAGGTTAACGCCGCCTGAggtaccacgt |
| **pGB1962** | pXMCS5 | pXMCS5-YFP-TacA | agacgaccatATGAGCAAGGGCGAGGAGCTGTTCACCGGGGTGGTGCCCATCCTGGTCGAGCTGGA<br>CGGCGACGTAAACGGCCACAAGTTCAGCGTGTCCGGCGAGGGCGAGGGCGATGCCACCTACGGC<br>AAGCTGACCCTGAAGTTCATCTGCACCACCGGCAAGCTGCCCGTGCCCTGGCCCACCCTCGTGAC<br>CACCTTCGGCTACGGCCTGCAGTGCTTCGCCCGCTACCCCGACCACATGAAGCAGCACGACTTCTT<br>CAAGTCCGCCATGCCCGAAGGCTACGTCCAGGAGCGCACCATCTTCTTCAAGGACGACGGCAACT<br>ACAAGACCCGCGCCGAGGTGAAGTTCGAGGGCGACACCCTGGTGAACCGCATCGAGCTGAAGG<br>GCATCGACTTCAAGGAGGACGGCAACATCCTGGGGCACAAGCTGGAGTACAACTACAACAGCCA<br>CAACGTCTATATCATGGCCGACAAGCAGAAGAACGGCATCAAGGTGAACTTCAAGATCCGCCACA<br>ACATCGAGGACGGCAGCGTGCAGCTCGCCGACCACTACCAGCAGAACACCCCCATCGGCGACGG<br>CCCCGTGCTGCTGCCCGACAACCACTACCTGAGCTACCAGTCCGCCCTGAGCAAAGACCCCAACG<br>AGAAGCGCGATCACATGGTCCTGCTGGAGTTCGTGACCGCCGCCGGGATCACTCTCGGCATGGAC<br>GAGCTGTACAAGGGAAGTAGCGGTGCGACCAAAACGGTCCTTGTCGTCGATGACGACCCGACAC<br>AACGTCGGTTGATTCAAGCGGTGCTCGAGCGCGACGGGTTCGCGGTCTCGCACGCGGAAGGCGG<br>CGACGCGGCGATCGCGCACCTGACGTCCGGCGCGCCCGCCGATGTCATCCTGCTGGATCTCGTCAT<br>GCCTGGCCTCAACGGTCAGGACGCCCTGAAGGAAATGCGCGCCCGCGGCTTCAACCAGCCCGTG<br>ATCGTGCTGACGGCCAGCGGTGGCGTCGACACCGTGGTCAAGGCCATGCAGGCCGGCGCCTGCG<br>ACTTCTTCATCAAGCCCGCCTCGCCCGAGCGGATCACCGTGTCGATCCGCAACGCCCTGTCGATGG<br>GCGACCTCAAGGGCGAGGTCGAGCGGCTGACCAAGCGCGCGGGCGGCAAGACCACCTTCGCGG<br>ACCTGATCGGCGCCTCGCCGGTCATGACCATGGTCAAGCGCATGGGCGAGCGGGCCGCCAAGAG<br>CGGTATCCCGGTGCTGATCACCGGCGAAAGCGGCGTCGGCAAGGAGCTGATCGCCCGCGCCGTC<br>CACGGCTCTTCCGACCGCGCCGGCAAGCCGTTTGTCGCGGTCAACTGCGGCGCGATCCCCGAGAA<br>CCTCGTCGAGTCGATCCTGTTCGGCCACGAGAAGGGCTCGTTCACCGGCGCCACCGACAAGCATC<br>TGGGCAAGTTCAAGGAGGCCGACGGCGGCACCCTGTTCCTCGATGAGGTCGGCGAGCTGCCGCT<br>CGACATGCAGGTCAAGCTGCTGCGCGCCCTGCAGGAGGGCGAGATCGACCCGATCGGCTCCAAG |

| | | | |
|---|---|---|---|
| | | | CGCTCGATCAAGGTCGATGTCCGGATCGTGTCGGCGACCAATCGCGATCTGCAGCAGGCCGTTTC<br>GGGCGGCCGGTTCCGCGAAGACCTGTTTTATCGCCTGAACGTGTTCCCGATCGAGGCGCCGTCCC<br>TGCGCGAGCGTCGCGAGGACATCCCGGCCCTCGTCGAGGCGTTCATCCGCCGGTTCAACGTCGAA<br>GAAGGCAAGCGCGTGATCGGCGCCTCGCCCGAGACGATGCAACTGCTGACCAGCTTCGACTGGC<br>CCGGCAATGTGCGCCAACTGGAAAACACCGTCTATCGCGCCATCGTGCTGGCCGATGCGCCCTATC<br>TGCAGCCGTTCGACTTCCCGGCGATCTCGGGCCTGGCCGCGCCGATCGAGGCCGTATCGATCTCGC<br>CCTCGCCGCCGCCTGCAGCCCTGTTGCAAGCCACGCATGCGGCGATGGCCGCCGCCGTCGCAGAG<br>GCGCCTGTGCGCATCCTCGACGATCGCGGTCACCTGCGGACCCTGGAGGAGATCGAGCGCGACCT<br>CATCCAGCATGCGATCGACGTCTATGCCGGCCACATGAGCGAGGTCGCGCGGCGTTTGGGCATCG<br>GTCGCTCGACCCTCTATCGCAAGGTTCGCGAGCAGGGCATCGAAGTCGACATGAAGGAAGCGGG<br>CTGAggtaccacgt |
| **pGB1963** | pXMCS5 | pXMCS5-<br>YFP-PdeA | agacgaccatATGAGCAAGGGCGAGGAGCTGTTCACCGGGGTGGTGCCCATCCTGGTCGAGCTGGA<br>CGGCGACGTAAACGGCCACAAGTTCAGCGTGTCCGGCGAGGGCGAGGGCGATGCCACCTACGGC<br>AAGCTGACCCTGAAGTTCATCTGCACCACCGGCAAGCTGCCCGTGCCCTGGCCCACCCTCGTGAC<br>CACCTTCGGCTACGGCCTGCAGTGCTTCGCCCGCTACCCCGACCACATGAAGCAGCACGACTTCTT<br>CAAGTCCGCCATGCCCGAAGGCTACGTCCAGGAGCGCACCATCTTCTTCAAGGACGACGGCAACT<br>ACAAGACCCGCGCCGAGGTGAAGTTCGAGGGCGACACCCTGGTGAACCGCATCGAGCTGAAGG<br>GCATCGACTTCAAGGAGGACGGCAACATCCTGGGGCACAAGCTGGAGTACAACTACAACAGCCA<br>CAACGTCTATATCATGGCCGACAAGCAGAAGAACGGCATCAAGGTGAACTTCAAGATCCGCCACA<br>ACATCGAGGACGGCAGCGTGCAGCTCGCCGACCACTACCAGCAGAACACCCCCATCGGCGACGG<br>CCCCGTGCTGCTGCCCGACAACCACTACCTGAGCTACCAGTCCGCCCTGAGCAAAGACCCCAACG<br>AGAAGCGCGATCACATGGTCCTGCTGGAGTTCGTGACCGCCGCCGGGATCACTCTCGGCATGGAC<br>GAGCTGTACAAGGGAAGTAGCGGTGCGTCTTTTCGGACAGGCGAACGACGAATCTGGGACGCGA<br>CGGCCACGCTTGAGGCGCTGGGCGCGGCGGACGTCGCCCTGTGGATCTGGGAGCCCGAAACCG<br>ACAGGTTGCGTCTGAACGGCGCGGCGCGCGCCTTGGGCCTTGGGCCGCTGGCGCCTGAATGCTC<br>GTCGGCCGCCTTCCGCGCCCTGGCCCTGCCGCAGGATCGCGCCCAGGCCGAAGAGGTCCTGAAG<br>CCGCGTGAACCGGGCAGCGAAGTCGTCGCCCGTTTCCGCGTGCGCGGCGGCGAGACCTGCCTCT<br>GGCGCGGCGTCTGGCTGGAAGAGGGCGTGCGCGCCGCCGGCGTCGTGGCGCCCGAAACGAAAT<br>TCTCCGCGTCCGAGCTTTGCGACCTGACCGGACTTCTGGACCGTCGCAGCTTCCTCGCCCGCGCCC<br>GCGAGCGCCTGGCGCAGGAGGGGACCCACCAGTTGGTCGTCGCCGACCTCGACCGACTGCGTCG<br>CCTGAACGAGGCGCTGGGTCACGAGCGCGCGGACCTAGTCCTGGCGGCGCTGGGCTCGCGCCTG<br>GCGGCGGCGTTCCCGGCCCAGTCGATCCTGGGCCGGATCGGCGAGGACGAGTTCGCCGTTCTCT<br>GCCAGCCGCTCGGCTACGAACCCTCCGATGTGCTGCGCAGCGCGCTGGAGCAGCCGCTGCGCGT<br>GGCCGGCTTTGATATTCACCCGACCCTGTCGATCGGCGCGGTCTCGGCCGAAGGCGGCCTGGACG<br>CGCCGGACGCCGCCGAACTGCTGCGCCGCGCCGAACTGGCCGTCGAGGCCGCCGCCGCCGCCG<br>GCCGAGGCGGCGCGGCCGCCTATGGCCGGGCGATGGAGACCGACGGCCTGTCGCGCCTGGCGC<br>TGGAGGCTGATCTGCGCGGCGCCATTGGTCGGGGCGAGATCACGCCCTACTTCCAGCCGATCGTG<br>CGGCTGTCGACCGGCGCCCTGTCCGGCTTCGAGGCCCTGGCCCGCTGGATCCATCCGCGCCGGG<br>GCATGCTGCCGCCGGACGAGTTCCTGCCGCTGATCGAAGAGATGGGGCTGATGAGCGAGCTCGG<br>CGCGCACATGATGCACGCCGCCGCCCAGCAGCTGTCGACCTGGCGCGCCGCTCACCCGGCGATGG<br>GGAACCTGACTGTCAGCGTCAACCTGTCGACCGGCGAGATCGACCGGCCCGGCCTGGTCGCCGA<br>CGTGGCCGAGACCCTGCGGGTCAATCGCCTGCCGCGCGGCGCCCTGAAGCTGGAAGTCACCGAA<br>AGCGACATCATGCGCGATCCCGAACGGGCCGCCGTGATCCTGAAGACGCTGCGCGACGCGGGCG<br>CAGGGCTTGCGCTGGACGACTTCGGCACCGGCTTCTCGTCGCTGTCGTACCTGACGCGCCTGCCG<br>TTCGACACGCTGAAGATCGACCGCTACTTCGTCCGCACCATGGGCAATAACGCCGGCTCGGCCAA<br>GATCGTCCGCTCGGTGGTCAAGCTGGGCCAGGATCTGGATCTGGAAGTCGTCGCCGAAGGGGTC<br>GAGAACGCCGAGATGGCGCATGCGCTGCAATCGCTGGGCTGTGACTATGGCCAAGGCTTTGGCTA<br>TGCGCCGGCCCTGTCGCCGCAGGAGGCCGAGGTCTATCTGAACGAGGCCTATGTCGACGGCGCC<br>GCGCCGGTGAAGGCGCGGGGTTAAggtaccacgt |
| **pGB1964** | pXMCS5 | pXMCS5-HA-<br>CtrA RD+15 | agacgaccatATGTACCCATACGATGTTCCAGATTACGCAGGAAGTCGCGTACTGTTGATCGAGGATG<br>ACAGCGCGACGGCGCAGACCATCGAACTGATGCTGAAGTCTGAAGGCTTCAACGTCTATACGACG<br>GATCTGGGTGAAGAAGGCGTCGATCTGGGCAAGATCTACGACTACGATCTTATCCTGCTCGACCTC |

| | | | |
|---|---|---|---|
| | | | AATCTTCCGGACATGAGCGGCATCGATGTTCTGCGCACCCTGCGGGTCGCGAAGATCAACACGCC CATCATGATCCTGTCGGGCTCGTCGGAAATCGACACCAAGGTCAAGACCTTCGCCGGCGGCGCCG ACGACTACATGACCAAGCCGTTCCACAAGGACGAAATGATCGCCCGCATCCACGCGGTGGTCCGT GGCTATGTGCTGCGCGACCCGAACGAGCAGGTTAACGCCGCCTGAggtaccacgt |
| **pGB1965** | pXMCS5 | pXMCS5-HA-TacA | agacgaccatATGTACCCATACGATGTTCCAGATTACGCAGGAAGTACCAAAACGGTCCTTGTCGTCG ATGACGACCCGACACAACGTCGGTTGATTCAAGCGGTGCTCGAGCGCGACGGGTTCGCGGTCTC GCACGCGGAAGGCGGCGACGCGGCGATCGCGCACCTGACGTCCGGCGCGCCCGCCGATGTCATC CTGCTGGATCTCGTCATGCCTGGCCTCAACGGTCAGGACGCCCTGAAGGAAATGCGCGCCCGCGG CTTCAACCAGCCCGTGATCGTGCTGACGGCCAGCGGTGGCGTCGACACCGTGGTCAAGGCCATGC AGGCCGGCGCCTGCGACTTCTTCATCAAGCCCGCCTCGCCCGAGCGGATCACCGTGTCGATCCGC AACGCCCTGTCGATGGGCGACCTCAAGGGCGAGGTCGAGCGGCTGACCAAGCGCGCGGGCGGC AAGACCACCTTCGCGGACCTGATCGGCGCCTCGCCGGTCATGACCATGGTCAAGCGCATGGGCGA GCGGGCCGCCAAGAGCGGTATCCCGGTGCTGATCACCGGCGAAAGCGGCGTCGGCAAGGAGCT GATCGCCCGCGCCGTCCACGGCTCTTCCGACCGCGCCGGCAAGCCGTTTGTCGCGGTCAACTGCG GCGCGATCCCCGAGAACCTCGTCGAGTCGATCCTGTTCGGCCACGAGAAGGGCTCGTTCACCGGC GCCACCGACAAGCATCTGGGCAAGTTCAAGGAGGCCGACGGCGGCACCCTGTTCCTCGATGAGG TCGGCGAGCTGCCGCTCGACATGCAGGTCAAGCTGCTGCGCGCCCTGCAGGAGGGCGAGATCGA CCCGATCGGCTCCAAGCGCTCGATCAAGGTCGATGTCCGGATCGTGTCGGCGACCAATCGCGATCT GCAGCAGGCCGTTTCGGGCGGCCGGTTCCGCGAAGACCTGTTTTATCGCCTGAACGTGTTCCCGA TCGAGGCGCCGTCCCTGCGCGAGCGTCGCGAGGACATCCCGGCCCTCGTCGAGGCGTTCATCCGC CGGTTCAACGTCGAAGAAGGCAAGCGCGTGATCGGCGCCTCGCCCGAGACGATGCAACTGCTGA CCAGCTTCGACTGGCCCGGCAATGTGCGCCAACTGGAAAACACCGTCTATCGCGCCATCGTGCTG GCCGATGCGCCCTATCTGCAGCCGTTCGACTTCCCGGCGATCTCGGGCCTGGCCGCGCCGATCGA GGCCGTATCGATCTCGCCCTCGCCGCCGCCTGCAGCCCTGTTGCAAGCCACGCATGCGGCGATGG CCGCCGCCGTCGCAGAGGCGCCTGTGCGCATCCTCGACGATCGCGGTCACCTGCGGACCCTGGAG GAGATCGAGCGCGACCTCATCCAGCATGCGATCGACGTCTATGCCGGCCACATGAGCGAGGTCGC GCGGCGTTTGGGCATCGGTCGCTCGACCCTCTATCGCAAGGTTCGCGAGCAGGGCATCGAAGTCG ACATGAAGGAAGCGGGCTGAggtaccacgt |
| **pGB1966** | pXMCS5 | pXMCS5-HA-PdeA | agacgaccatATGTACCCATACGATGTTCCAGATTACGCAGGAAGTTCTTTTCGGACAGGCGAACGAC GAATCTGGGACGCGACGGCCACGCTTGAGGCGCTGGGCGCGGCGGACGTCGCCCTGTGGATCTG GGAGCCCGAAACCGACAGGTTGCGTCTGAACGGCGCGGCGCGCGCCTTGGGCCTTGGGCCGCT GGCGCCTGAATGCTCGTCGGCCGCCTTCCGCGCCCTGGCCCTGCCGCAGGATCGCGCCCAGGCCG AAGAGGTCCTGAAGCCGCGTGAACCGGGCAGCGAAGTCGTCGCCCGTTTCCGCGTGCGCGGCG GCGAGACCTGCCTCTGGCGCGGCGTCTGGCTGGAAGAGGGCGTGCGCGCCGCCGGCGTCGTGG CGCCCGAAACGAAATTCTCCGCGTCCGAGCTTTGCGACCTGACCGGACTTCTGGACCGTCGCAGC TTCCTCGCCCGCGCCCGCGAGCGCCTGGCGCAGGAGGGGACCCACCAGTTGGTCGTCGCCGACC TCGACCGACTGCGTCGCCTGAACGAGGCGCTGGGTCACGAGCGCGCGGACCTAGTCCTGGCGGC GCTGGGCTCGCGCCTGGCGGCGGCGTTCCCGGCCCAGTCGATCCTGGGCCGGATCGGCGAGGAC GAGTTCGCCGTTCTCTGCCAGCCGCTCGGCTACGAACCCTCCGATGTGCTGCGCAGCGCGCTGGA GCAGCCGCTGCGCGTGGCCGGCTTTGATATTCACCCGACCCTGTCGATCGGCGCGGTCTCGGCCG AAGGCGGCCTGGACGCGCCGGACGCCGCCGAACTGCTGCGCCGCGCCGAACTGGCCGTCGAGG CCGCCGCCGCCGCCGGCCGAGGCGGCGCGGCCGCCTATGGCCGGGCGATGGAGACCGACGGCC TGTCGCGCCTGGCGCTGGAGGCTGATCTGCGCGGCGCCATTGGTCGGGGCGAGATCACGCCCTAC TTCCAGCCGATCGTGCGGCTGTCGACCGGCGCCCTGTCCGGCTTCGAGGCCCTGGCCCGCTGGAT CCATCCGCGCCGGGGCATGCTGCCGCCGGACGAGTTCCTGCCGCTGATCGAAGAGATGGGGCTG ATGAGCGAGCTCGGCGCGCACATGATGCACGCCGCCGCCCAGCAGCTGTCGACCTGGCGCGCCG CTCACCCGGCGATGGGGAACCTGACTGTCAGCGTCAACCTGTCGACCGGCGAGATCGACCGGCCC GGCCTGGTCGCCGACGTGGCCGAGACCCTGCGGGTCAATCGCCTGCCGCGCGGCGCCCTGAAGC TGGAAGTCACCGAAAGCGACATCATGCGCGATCCCGAACGGGCCGCCGTGATCCTGAAGACGCTG CGCGACGCGGGCGCAGGGCTTGCGCTGGACGACTTCGGCACCGGCTTCTCGTCGCTGTCGTACCT GACGCGCCTGCCGTTCGACACGCTGAAGATCGACCGCTACTTCGTCCGCACCATGGGCAATAACG CCGGCTCGGCCAAGATCGTCCGCTCGGTGGTCAAGCTGGGCCAGGATCTGGATCTGGAAGTCGTC GCCGAAGGGGTCGAGAACGCCGAGATGGCGCATGCGCTGCAATCGCTGGGCTGTGACTATGGCC |

| | | | |
|---|---|---|---|
| | | | AAGGCTTTGGCTATGCGCCGGCCCTGTCGCCGCAGGAGGCCGAGGTCTATCTGAACGAGGCCTAT<br>GTCGACGGCGCCGCGCCGGTGAAGGCGCGGGGTTAAggtaccacgt |
| **pGB1967** | pVMCS6<br>(NdeI,<br>KpnI) | pVMCS6-<br>ClpX-GFP | ggaaacgcatATGACGAAAGCCGCGAGCGGCGACACGAAAAGCACCCTGTACTGCTCTTTCTGCGG<br>AAAGAGCCAACATGAGGTGCGCAAGCTCATCGCGGGACCGACGGTGTTCATTTGCGATGAATGCG<br>TCGAGCTCTGCATGGACATCATCCGCGAAGAGCACAAGATCGCCTTCGTGAAGTCTAAGGACGGC<br>GTCCCGACGCCGCGCGAAATCTGCGAAGTCCTGGATGATTACGTGATCGGTCAAGGTCACGCCAA<br>GAAGGTCCTCGCGGTCGCAGTGCACAATCACTACAAGCGGCTGAACCACGCTTCGAAGAATAACG<br>ACGTCGAACTGGCCAAGTCGAACATCCTGCTGGTCGGTCCGACCGGTACGGGTAAGACCCTGCTG<br>GCGCAGACGCTGGCCCGAATCATCGACGTTCCGTTCACGATGGCGGACGCCACGACGCTGACCGA<br>AGCCGGTTACGTCGGCGAAGACGTCGAGAACATCGTGCTGAAGCTGCTGCAGGCCGCCGACTAC<br>AACGTCGAGCGCGCCCAGCGCGGCATCGTCTACATCGACGAAATCGACAAGATCAGCCGCAAGTC<br>CGACAACCCGTCGATCACTCGCGACGTGTCGGGCGAGGGCGTGCAGCAGGCTCTGCTGAAGATC<br>ATGGAAGGCACGGTCGCCTCCGTGCCGCCGCAGGGCGGGCGCAAGCATCCTCAGCAGGAGTTCC<br>TGCAGGTCGACACGACGAACATCCTGTTCATCTGTGGCGGCGCCTTCGCTGGCCTGGAGAAGATC<br>ATCTCGGCGCGCGGCGCGGCCAAGTCGATCGGCTTCGGCGCCAAGGTGACCGATCCCGAAGAGC<br>GCCGGACGGGCGAGATCCTTCGGAACGTCGAGCCCGACGACCTGCAGCGTTTCGGCCTGATCCC<br>GGAGTTCATCGGCCGTCTGCCGGTGGTCGCCACGCTGGAGGATCTGGACGAGGCCGCCCTGGTC<br>AAGATCCTGACCGAGCCGAAGAACGCCTTCGTCAAGCAGTATCAGCGCCTGTTCGAGATGGAGAA<br>CATCGGCCTGACCTTCACCGAAGACGCTCTGCATCAGGTGGCCAAGAAGGCTATCGCGCGCAAGA<br>CCGGCGCGCGCGGCCTGCGCTCGATCATGGAAGGCATCCTGCTGGAGACCATGTTCGAACTGCCG<br>ACCTACGAGGGCGTCGAGGAAGTGGTGGTCAACGCCGAGGTCGTCGAAGGCCGGGCTCAGCCG<br>CTGCTGATCTATGCCGAGAAAAAGGGTGGGGCGGCATCGGCCGGAAGTAGTAAAGGTGAAGAAC<br>TGTTCACCGGTGTTGTTCCGATCCTGGTTGAACTGGATGGTGATGTTAACGGCCACAAATTCTCTGT<br>TCGTGGTGAAGGTGAAGGTGATGCAACCAACGGTAAACTGACCCTGAAATTCATCTGCACTACCG<br>GTAAACTGCCGGTTCCATGGCCGACTCTGGTGACTACCCTGACCTATGGTGTTCAGTGTTTTTCTCG<br>TTACCCGGATCACATGAAGCAGCATGATTTCTTCAAATCTGCAATGCCGGAAGGTTATGTACAGGA<br>GCGCACCATTTCTTTCAAAGACGATGGCACCTACAAAACCCGTGCAGAGGTTAAATTTGAAGGTG<br>ATACTCTGGTGAACCGTATTGAACTGAAAGGCATTGATTTCAAAGAGGACGGCAACATCCTGGGC<br>CACAAACTGGAATATAACTTCAACTCCCATAACGTTTACATCACCGCAGACAAACAGAAGAACGGT<br>ATCAAAGCTAACTTCAAAATTCGCCATAACGTTGAAGACGGTAGCGTACAGCTGGCGGACCACTAC<br>CAGCAGAACACTCCGATCGGTGATGGTCCGGTTCTGCTGCCGGATAACCACTACCTGTCCACCCAG<br>TCTAAACTGTCCAAAGACCCGAACGAAAAGCGCGACCACATGGTGCTGCTGGAGTTCGTTACTGC<br>AGCAGGTATCACGCACGGCATGGATGAACTCTACAAATAGggtaccacgt |
| **pGB1968** | pVMCS6 | pVMCS6-<br>RcdA-GFP | ggaaacgcatATGACCGAAGTGAACGCGTTCGCGGACACGCCTTGGCGCGCTGGAGTGATCCAGGA<br>TTTCGCGCGATCGGAACTGTTCGACCGGACGTTCGAGGAAGGCATGCAACTGGTAGAAGAGACC<br>GCCGCCTATCTCGACGGGGCCGGACGCCATGACAGCAAGGTCCTCTCTCGCAACGCCGCCCTGGG<br>CTACGCCACCGAAAGCATGCGCCTGACCACGCGCCTGATGCAGGTCGCCTCCTGGCTTTTGGTGC<br>AGCGCGCCGTACGTGAAGGCGAGATGCCGCCGGAAGCCGCCTGCGCTGAAGCCTATCGCCTGGC<br>CGAAGAGGCCCCGGCCGATGGTCCGGCCGTCGAGGAACTGCCGTTTGGCCTGATGAACCTGCTG<br>CAGCGCTCCGAGCGCCTGTACGAGCGCGTCCGCCACCTGGACCGCCGCATGTATGTCGAGTCGCC<br>GAACGAAGAAGCGCCGCGTCCGGTTCAGAACCAGCTCGATCGCTTGACGGCGGCGTTCGGAGGC<br>AGCGGCGGCGGCGGCAGTAAAGGTGAAGAACTGTTCACCGGTGTTGTTCCGATCCTGGTTGAAC<br>TGGATGGTGATGTTAACGGCCACAAATTCTCTGTTCGTGGTGAAGGTGAAGGTGATGCAACCAAC<br>GGTAAACTGACCCTGAAATTCATCTGCACTACCGGTAAACTGCCGGTTCCATGGCCGACTCTGGTG<br>ACTACCCTGACCTATGGTGTTCAGTGTTTTTCTCGTTACCCGGATCACATGAAGCAGCATGATTTCTT<br>CAAATCTGCAATGCCGGAAGGTTATGTACAGGAGCGCACCATTTCTTTCAAAGACGATGGCACCTA |

| | | | |
|---|---|---|---|
| | | | CAAAACCCGTGCAGAGGTTAAATTTGAAGGTGATACTCTGGTGAACCGTATTGAACTGAAAGGCA<br>TTGATTTCAAAGAGGACGGCAACATCCTGGGCCACAAACTGGAATATAACTTCAACTCCCATAACG<br>TTTACATCACCGCAGACAAACAGAAGAACGGTATCAAAGCTAACTTCAAAATTCGCCATAACGTTG<br>AAGACGGTAGCGTACAGCTGGCGGACCACTACCAGCAGAACACTCCGATCGGTGATGGTCCGGTT<br>CTGCTGCCGGATAACCACTACCTGTCCACCCAGTCTAAACTGTCCAAAGACCCGAACGAAAAGCGC<br>GACCACATGGTGCTGCTGGAGTTCGTTACTGCAGCAGGTATCACGCACGGCATGGATGAACTCTAC<br>AAATAGggtaccacgt |
| **pGB1969** | pACYC-duet (NcoI, XhoI) | pACYC-CpdR-TEV-GFP-6XHis | gagatataccATGGCAAGAATTCTCCTCGCTGAAGATGATGATTCTCTGAGAGGATTTCTGGCTAGAG<br>CTCTGGAAAGAGCTGGATTTGAAGTCCAGGCTTGCGCTGATGGAGAAGAAGCTGTCCAGCACCT<br>GGATCATCCTTGGGATCTGCTGCTGACAGATATTGTCATGCCTGGAATGGATGGAATTGAAGTGGC<br>TAGACAGGCTGCTGCTAGAGATCCGAGTCTGAGAATTATGTTTATTACAGGATTTGCTGCTGTGGCT<br>CTCTCGGCTCAGGATAGAGCTCCTGCTGGAGCTAAGGTGCTGAGTAAGCCTGTGCACCTGAGAGA<br>TCTCGTCGCTGAAGTCGAAAAGATGATGGCTGCTGAAAACCTGTATTTTCAGGGTAGCAGTAAAG<br>GTGAAGAACTGTTCACCGGTGTTGTTCCGATCCTGGTTGAACTGGATGGTGATGTTAACGGCCACA<br>AATTCTCTGTTCGTGGTGAAGGTGAAGGTGATGCAACCAACGGTAAACTGACCCTGAAATTCATCT<br>GCACTACCGGTAAACTGCCGGTTCCATGGCCGACTCTGGTGACTACCCTGACCTATGGTGTTCAGT<br>GTTTTTCTCGTTACCCGGATCACATGAAGCAGCATGATTTCTTCAAATCTGCAATGCCGGAAGGTTA<br>TGTACAGGAGCGCACCATTTCTTTCAAAGACGATGGCACCTACAAAACCCGTGCAGAGGTTAAATT<br>TGAAGGTGATACTCTGGTGAACCGTATTGAACTGAAAGGCATTGATTTCAAAGAGGACGGCAACA<br>TCCTGGGCCACAAACTGGAATATAACTTCAACTCCCATAACGTTTACATCACCGCAGACAAACAGA<br>AGAACGGTATCAAAGCTAACTTCAAAATTCGCCATAACGTTGAAGACGGTAGCGTACAGCTGGCG<br>GACCACTACCAGCAGAACACTCCGATCGGTGATGGTCCGGTTCTGCTGCCGGATAACCACTACCTG<br>TCCACCCAGTCTaaaCTGTCCAAAGACCCGAACGAAAAGCGCGACCACATGGTGCTGCTGGAGTTC<br>GTTACTGCAGCAGGTATCACGCACGGCATGGATGAACTCTACAAACACCATCACCATCACCATTAGc<br>tcgagtctg |
| **pGB1970** | pACYC-duet | pACYC-CpdR$_{D51A}$-TEV-GFP-6XHis | gagatataccATGGCAAGAATTCTCCTCGCTGAAGATGATGATTCTCTGAGAGGATTTCTGGCTAGAG<br>CTCTGGAAAGAGCTGGATTTGAAGTCCAGGCTTGCGCTGATGGAGAAGAAGCTGTCCAGCACCT<br>GGATCATCCTTGGGATCTGCTGCTGACAGCCATTGTCATGCCTGGAATGGATGGAATTGAAGTGGC<br>TAGACAGGCTGCTGCTAGAGATCCGAGTCTGAGAATTATGTTTATTACAGGATTTGCTGCTGTGGCT<br>CTCTCGGCTCAGGATAGAGCTCCTGCTGGAGCTAAGGTGCTGAGTAAGCCTGTGCACCTGAGAGA<br>TCTCGTCGCTGAAGTCGAAAAGATGATGGCTGCTGAAAACCTGTATTTTCAGGGTAGCAGTAAAG<br>GTGAAGAACTGTTCACCGGTGTTGTTCCGATCCTGGTTGAACTGGATGGTGATGTTAACGGCCACA<br>AATTCTCTGTTCGTGGTGAAGGTGAAGGTGATGCAACCAACGGTAAACTGACCCTGAAATTCATCT<br>GCACTACCGGTAAACTGCCGGTTCCATGGCCGACTCTGGTGACTACCCTGACCTATGGTGTTCAGT<br>GTTTTTCTCGTTACCCGGATCACATGAAGCAGCATGATTTCTTCAAATCTGCAATGCCGGAAGGTTA<br>TGTACAGGAGCGCACCATTTCTTTCAAAGACGATGGCACCTACAAAACCCGTGCAGAGGTTAAATT<br>TGAAGGTGATACTCTGGTGAACCGTATTGAACTGAAAGGCATTGATTTCAAAGAGGACGGCAACA<br>TCCTGGGCCACAAACTGGAATATAACTTCAACTCCCATAACGTTTACATCACCGCAGACAAACAGA<br>AGAACGGTATCAAAGCTAACTTCAAAATTCGCCATAACGTTGAAGACGGTAGCGTACAGCTGGCG<br>GACCACTACCAGCAGAACACTCCGATCGGTGATGGTCCGGTTCTGCTGCCGGATAACCACTACCTG<br>TCCACCCAGTCTaaaCTGTCCAAAGACCCGAACGAAAAGCGCGACCACATGGTGCTGCTGGAGTTC<br>GTTACTGCAGCAGGTATCACGCACGGCATGGATGAACTCTACAAACACCATCACCATCACCATTAGc<br>tcgagtctg |
| **pGB2010** | | pBXMCS2-pCpdR CpdR | agacgaccatTCACGGTTCAGATCCACATAGGCGCGGGCGAAGCGGGCGCGCACCACGGCGGCCCC<br>CAGGGCGGGCGCGCCGCCGATGATGCGGTCGACAAACGCGTCCTCGGACGCGCGCAGGGTCTC<br>GAGCGGCAGGCGGACCGCGGCGACCATGTCCTCCGGATAGAGGTCGCCCGAGTGCGGCGAGGC<br>GAACACCAAGGCGGTCGGCGGCGGCGCCCCGGCCGGGACCGCGCGCAGCACCTCGAACGCTGG<br>CCCGCCAAAGGTCTCCAGCGGCGGGGACGAAAGCTCTTCGCTCGAAAGGGGCCGCCACGCGCTC<br>ATGCCGCATCATCGAAGCGTCGCGGCGCGGGGTCAAATCTCAATCGCCGCTGACGAAACATCCCC<br>AGCCGCGACGTTTAGGTTCATCCCCGATTTACGGACGGGGCGATAAGGTGGATCCTCTATCGACGA<br>TCTTAATCGGACACGTGACCCCatgGCCCGCATCCTCCTCGCCGAAGACGATGATTCCCTGCGCGGC<br>TTCCTGGCCCGCGCGCTGGAACGCGCCGGCTTCGAAGTCCAGGCCTGCGCCGACGGCGAAGAGG<br>CCGTCCAGCACCTGGACCATCCCTGGGACCTGCTGCTGACCGACATCGTCATGCCCGGCATGGACG |

| | | | |
|---|---|---|---|
| | | | GCATCGAGGTGGCCCGCCAGGCCGCCGCCCGCGACCCGTCCCTGCGCATCATGTTCATCACCGGC<br>TTCGCCGCCGTGGCCCTCTCGGCCCAGGACCGCGCGCCCGCCGGCGCCAAGGTGCTGTCCAAGC<br>CCGTGCACCTGCGCGACCTCGTCGCCGAGGTCGAAAAGATGATGGCGGCCtgagaattcctgc |
| **pGB2011** | | pBXMCS2-<br>pCpdR<br>$CpdR_{D51A}$ | agacgaccatTCACGGTTCAGATCCACATAGGCGCGGGCGAAGCGGGCGCGCACCACGGCGGCCCC<br>CAGGGCGGGCGCGCCGCCGATGATGCGGTCGACAAACGCGTCCTCGGACGCGCGCAGGGTCTC<br>GAGCGGCAGGCGGACCGCGGCGACCATGTCCTCCGGATAGAGGTCGCCCGAGTGCGGCGAGGC<br>GAACACCAAGGCGGTCGGCGGCGGCGCCCCGGCCGGGACCGCGCGCAGCACCTCGAACGCTGG<br>CCCGCCAAAGGTCTCCAGCGGCGGGGACGAAAGCTCTTCGCTCGAAAGGGGCCGCCACGCGCTC<br>ATGCCGCATCATCGAAGCGTCGCGGCGCGGGGTCAAATCTCAATCGCCGCTGACGAAACATCCCC<br>AGCCGCGACGTTTAGGTTCATCCCCGATTTACGGACGGGGCGATAAGGTGGATCCTCTATCGACGA<br>TCTTAATCGGACACGTGACCCCatgGCCCGCATCCTCCTCGCCGAAGACGATGATTCCCTGCGCGGC<br>TTCCTGGCCCGCGCGCTGGAACGCGCCGGCTTCGAAGTCCAGGCCTGCGCCGACGGCGAAGAGG<br>CCGTCCAGCACCTGGACCATCCCTGGGACCTGCTGCTGACCGCCATCGTCATGCCCGGCATGGACG<br>GCATCGAGGTGGCCCGCCAGGCCGCCGCCCGCGACCCGTCCCTGCGCATCATGTTCATCACCGGC<br>TTCGCCGCCGTGGCCCTCTCGGCCCAGGACCGCGCGCCCGCCGGCGCCAAGGTGCTGTCCAAGC<br>CCGTGCACCTGCGCGACCTCGTCGCCGAGGTCGAAAAGATGATGGCGGCCtgagaattcctgc |
| **pGB2012** | | pBXMCS2-<br>pXyl-CckA<br>K+-pCpdR<br>CpdR | agacgaccatATGGCCGACTTGCAGCTCCAGGACAAGGTTTCGACCGGCGCCCCGCGTCGGCGGTTT<br>GATCCATGGCTGGTCGGCGCGGCGGTGTTTTTCGTGGCTGCGGCGGCCCTCTCGGCGGCGCCGG<br>CGCTCAAGGCCGGACCGACCACCCTGGCCGGCCTGCTGCTGCTTCTGGGCGTCGCGGGCGTGGC<br>TGTGTTGGGCCTTGTCGCCATTCGCGGCTCAGCGCTTTCCGGCGGCGACGCCGACCAGGCTGAG<br>GGGTTCATCGAGGCGCTGGCCGAGCCGGCCGCCCTGGCCGCCGCCGACGGTCGCGTCCTGGCCG<br>CCAACGGGCCCTGGCGCGAAGTCATGGGCGAACAGCGCCGCCTGCCCAAGGGCGTGGCGGGCT<br>CCAGCCTGTTTGCGGCCCTGGTCCAGGCGCGCCAGGGGCAGATGGCCGAGGGCATGCTGAGCGC<br>TGGAGGAACCGACTATACCGCCAAGGTCTCGCGCCTTGCGGGCGGACGGCTGATGATCCGGCTTG<br>CGCCCATCGTTGTCGCTGAGCCGGTTGTGGAAGACGCGTCGCCGGCCCCGGTCGCCGAGCGCGC<br>CGCCCCGCCACCCAGCTCGCTGGACGCCTTCGCCGGAGCCTCGCCGTTCGGCGCGGCCCTGCTGG<br>AAGGCCTGGAGCCGTTCACCTCGCGGGTGCTGGAGACCAATCCGGCGCTGACCACGATGACCGG<br>CGCCAAGGCCGGTGTGCTGTTCGGCGATCTGATCGACGCCGCCTCGCGCGCCGAGGCCGAGACG<br>CGCCTGAACGAAGGCCGCGCCGGTCCCTACGAGGTGCGGTTGGCGCGCGATCCGTCGCGCATCG<br>CTCACCTTTACCTCTATCGCGCCGAAGGCCGCCTGGTGGCCTACATGATCGACGTGTCCGAGCAGA<br>AGCAGATCGAGCTGCAGCTGTCCCAGGCCCAGAAGATGCAGGCCATCGGCCAGCTGGCCGGCGG<br>CGTCGCGCACGACTTCAACAACCTCTTGACCGCCATCCAGCTGCGTCTGGACGAGTTGCTGCATCG<br>CCATCCCGTCGGTGATCCGTCGTACGAGGGTCTCAACGAGATCCGTCAGACGGGCGTGCGCGCCG<br>CCGACCTCGTGCGCAAGCTCTTGGCTTTCTCGCGCAAGCAGACCGTGCAGCGCGAGGTGCTGGAT<br>CTGGGCGAGCTGATCAGCGAGTTCGAGGTCTTGCTGCGCCGCCTGCTGCGCGAAGACGTCAAGC<br>TGATCACCGACTATGGCCGCGACCTGCCGCAGGTGCGCGCCGACAAGAGCCAGCTCGAGACGGC<br>GGTCATGAACCTGGCCGTCAACGCCCGCGACGCCGTGCGCGCGGCCAAGGGCGGCGGCGTCGT<br>GCGCATCCGCACCGCGCGCCTGACCCGCGACGAGGCGATCCAGCTGGGTTTCCCGGCCGCCGAC<br>GGCGACACGGCCTTCATTGAGGTCAGTGACGATGGTCCGGGCATTCCGCCCGACGTCATGGGCAA<br>GATCTTCGACCCGTTCTTCACCACCAAGCCGGTGGGCGAGGGTACGGGCCTAGGCCTAGCCACGG<br>TCTATGGCATCGTTAAGCAGAGCGACGGCTGGATTCACGTCCACAGCCGTCCGAACGAAGGCGCG<br>GCCTTCCGCATCTTCCTGCCGGTCTATGAAGCGCCCGCCGGCGCGGTCGCCGTCCAGGCCGTCGC<br>CGAGCCCGCCAAGCCGCGCGCCGCTCGCGACCTGTCGGGCGCCGGCCGCATCCTGTTCGTCGAG<br>GACGAGGACGCCGTGCGCAGCGTCGCCGCCCGCCTGCTGCGCGCCCGTGGCTACGAGGTGCTTG<br>AGGCGGCCGACGGCGAAGAGGCCCTGATCATCGCCGAGGAAAACGCCGGCACGATCGACCTCTT<br>GATCAGCGACGTGATCATGCCCGGCATCGACGGCCCGACCTTGCTGAAGAAGGCGCGTGGCTATC<br>TGGGGACCGCGCCGGTGATGTTCATCTCCGGCTATGCCGAGGCCGAGTTCAGCGACCTCTTGGAA<br>GGCGAGACGGGCGTGACCTTCCTCCCCAAGCCGATCGACATCAAGACCCTGGCCGAGCGCGTCA<br>AGCAGCAGCTGCAAGCAGCATAGtctgtgctattctaacgaatcttgattcgTCACGGTTCAGATCCACATAG<br>GCGCGGGCGAAGCGGGCGCGCACCACGGCGGCCCCCAGGGCGGGCGCGCCGCCGATGATGCG<br>GTCGACAAACGCGTCCTCGGACGCGCGCAGGGTCTCGAGCGGCAGGCGGACCGCGGCGACCAT<br>GTCCTCCGGATAGAGGTCGCCCGAGTGCGGCGAGGCGAACACCAAGGCGGTCGGCGGCGGCGC |

| | | | |
|---|---|---|---|
| | | | CCCGGCCGGGACCGCGCGCAGCACCTCGAACGCTGGCCCGCCAAAGGTCTCCAGCGGCGGGGA<br>CGAAAGCTCTTCGCTCGAAAGGGGCCGCCACGCGCTCATGCCGCATCATCGAAGCGTCGCGGCGC<br>GGGGTCAAATCTCAATCGCCGCTGACGAAACATCCCCAGCCGCGACGTTTAGGTTCATCCCCGATT<br>TACGGACGGGGCGATAAGGTGGATCCTCTATCGACGATCTTAATCGGACACGTGACCCCatgGCCC<br>GCATCCTCCTCGCCGAAGACGATGATTCCCTGCGCGGCTTCCTGGCCCGCGCGCTGGAACGCGCC<br>GGCTTCGAAGTCCAGGCCTGCGCCGACGGCGAAGAGGCCGTCCAGCACCTGGACCATCCCTGGG<br>ACCTGCTGCTGACCGACATCGTCATGCCCGGCATGGACGGCATCGAGGTGGCCCGCCAGGCCGCC<br>GCCCGCGACCCGTCCCTGCGCATCATGTTCATCACCGGCTTCGCCGCCGTGGCCCTCTCGGCCCAG<br>GACCGCGCGCCCGCCGGCGCCAAGGTGCTGTCCAAGCCCGTGCACCTGCGCGACCTCGTCGCCG<br>AGGTCGAAAAGATGATGGCGGCCtgagaattcctgc |
| **pGB2013** | | pBXMCS2-pXyl-CckAK+-pCpdR $CpdR_{D51A}$ | agacgaccatATGGCCGACTTGCAGCTCCAGGACAAGGTTTCGACCGGCGCCCCGCGTCGGCGGTTT<br>GATCCATGGCTGGTCGGCGCGGCGGTGTTTTTCGTGGCTGCGGCGGCCCTCTCGGCGGCGCCGG<br>CGCTCAAGGCCGGACCGACCACCCTGGCCGGCCTGCTGCTGCTTCTGGGCGTCGCGGGCGTGGC<br>TGTGTTGGGCCTTGTCGCCATTCGCGGCTCAGCGCTTTCCGGCGGCGACGCCGACCAGGCTGAG<br>GGGTTCATCGAGGCGCTGGCCGAGCCGGCCGCCCTGGCCGCCGCCGACGGTCGCGTCCTGGCCG<br>CCAACGGGCCCTGGCGCGAAGTCATGGGCGAACAGCGCCGCCTGCCCAAGGGCGTGGCGGGCT<br>CCAGCCTGTTTGCGGCCCTGGTCCAGGCGCGCCAGGGGCAGATGGCCGAGGGCATGCTGAGCGC<br>TGGAGGAACCGACTATACCGCCAAGGTCTCGCGCCTTGCGGGCGGACGGCTGATGATCCGGCTTG<br>CGCCCATCGTTGTCGCTGAGCCGGTTGTGGAAGACGCGTCGCCGGCCCCGGTCGCCGAGCGCGC<br>CGCCCCGCCACCCAGCTCGCTGGACGCCTTCGCCGGAGCCTCGCCGTTCGGCGCGGCCCTGCTGG<br>AAGGCCTGGAGCCGTTCACCTCGCGGGTGCTGGAGACCAATCCGGCGCTGACCACGATGACCGG<br>CGCCAAGGCCGGTGTGCTGTTCGGCGATCTGATCGACGCCGCCTCGCGCGCCGAGGCCGAGACG<br>CGCCTGAACGAAGGCCGCGCCGGTCCCTACGAGGTGCGGTTGGCGCGCGATCCGTCGCGCATCG<br>CTCACCTTTACCTCTATCGCGCCGAAGGCCGCCTGGTGGCCTACATGATCGACGTGTCCGAGCAGA<br>AGCAGATCGAGCTGCAGCTGTCCCAGGCCCAGAAGATGCAGGCCATCGGCCAGCTGGCCGGCGG<br>CGTCGCGCACGACTTCAACAACCTCTTGACCGCCATCCAGCTGCGTCTGGACGAGTTGCTGCATCG<br>CCATCCCGTCGGTGATCCGTCGTACGAGGGTCTCAACGAGATCCGTCAGACGGGCGTGCGCGCCG<br>CCGACCTCGTGCGCAAGCTCTTGGCTTTCTCGCGCAAGCAGACCGTGCAGCGCGAGGTGCTGGAT<br>CTGGGCGAGCTGATCAGCGAGTTCGAGGTCTTGCTGCGCCGCCTGCTGCGCGAAGACGTCAAGC<br>TGATCACCGACTATGGCCGCGACCTGCCGCAGGTGCGCGCCGACAAGAGCCAGCTCGAGACGGC<br>GGTCATGAACCTGGCCGTCAACGCCCGCGACGCCGTGCGCGCGGCCAAGGGCGGCGGCGTCGT<br>GCGCATCCGCACCGCGCGCCTGACCCGCGACGAGGCGATCCAGCTGGGTTTCCCGGCCGCCGAC<br>GGCGACACGGCCTTCATTGAGGTCAGTGACGATGGTCCGGGCATTCCGCCCGACGTCATGGGCAA<br>GATCTTCGACCCGTTCTTCACCACCAAGCCGGTGGGCGAGGGTACGGGCCTAGGCCTAGCCACGG<br>TCTATGGCATCGTTAAGCAGAGCGACGGCTGGATTCACGTCCACAGCCGTCCGAACGAAGGCGCG<br>GCCTTCCGCATCTTCCTGCCGGTCTATGAAGCGCCCGCCGGCGCGGTCGCCGTCCAGGCCGTCGC<br>CGAGCCCGCCAAGCCGCGCGCCGCTCGCGACCTGTCGGGCGCCGGCCGCATCCTGTTCGTCGAG<br>GACGAGGACGCCGTGCGCAGCGTCGCCGCCCGCCTGCTGCGCGCCCGTGGCTACGAGGTGCTTG<br>AGGCGGCCGACGGCGAAGAGGCCCTGATCATCGCCGAGGAAAACGCCGGCACGATCGACCTCTT<br>GATCAGCGACGTGATCATGCCCGGCATCGACGGCCCGACCTTGCTGAAGAAGGCGCGTGGCTATC<br>TGGGGACCGCGCCGGTGATGTTCATCTCCGGCTATGCCGAGGCCGAGTTCAGCGACCTCTTGGAA<br>GGCGAGACGGGCGTGACCTTCCTCCCCAAGCCGATCGACATCAAGACCCTGGCCGAGCGCGTCA<br>AGCAGCAGCTGCAAGCAGCATAGtctgtgctattctaacgaatcttgattcgTCACGGTTCAGATCCACATAG<br>GCGCGGGCGAAGCGGGCGCGCACCACGGCGGCCCCCAGGGCGGGCGCGCCGCCGATGATGCG<br>GTCGACAAACGCGTCCTCGGACGCGCGCAGGGTCTCGAGCGGCAGGCGGACCGCGGCGACCAT<br>GTCCTCCGGATAGAGGTCGCCCGAGTGCGGCGAGGCGAACACCAAGGCGGTCGGCGGCGGCGC<br>CCCGGCCGGGACCGCGCGCAGCACCTCGAACGCTGGCCCGCCAAAGGTCTCCAGCGGCGGGGA |

| | | | |
|---|---|---|---|
| | | | CGAAAGCTCTTCGCTCGAAAGGGGCCGCCACGCGCTCATGCCGCATCATCGAAGCGTCGCGGCGC<br>GGGGTCAAATCTCAATCGCCGCTGACGAAACATCCCCAGCCGCGACGTTTAGGTTCATCCCCGATT<br>TACGGACGGGGCGATAAGGTGGATCCTCTATCGACGATCTTAATCGGACACGTGACCCCatgGCCC<br>GCATCCTCCTCGCCGAAGACGATGATTCCCTGCGCGGCTTCCTGGCCCGCGCGCTGGAACGCGCC<br>GGCTTCGAAGTCCAGGCCTGCGCCGACGGCGAAGAGGCCGTCCAGCACCTGGACCATCCCTGGG<br>ACCTGCTGCTGACCGCCATCGTCATGCCCGGCATGGACGGCATCGAGGTGGCCCGCCAGGCCGCC<br>GCCCGCGACCCGTCCCTGCGCATCATGTTCATCACCGGCTTCGCCGCCGTGGCCCTCTCGGCCCAG<br>GACCGCGCGCCCGCCGGCGCCAAGGTGCTGTCCAAGCCCGTGCACCTGCGCGACCTCGTCGCCG<br>AGGTCGAAAAGATGATGGCGGCCtgagaattcctgc |

**Supplementary Table 3**

| | Diffusion Coefficients | | | | | | | | PopZ Microdomain Size | |
|---|---|---|---|---|---|---|---|---|---|---|
| | Top Cell | | | | Bottom Cell | | | | Top Cell | Bottom Cell |
| File # | Y_cyto | R_cyto | Y_pole | R_pole | Y_cyto | R_cyto | Y_pole | R_pole | | |
| 1 | 20 | 20 | 0.5 | 0.5 | 20 | 20 | 0.5 | 0.5 | 0.50% | 0.125% |
| 2 | 20 | 20 | 0.5 | 0.5 | 20 | 20 | 0.5 | 0.5 | 0.50% | 2.000% |
| 3 | 20 | 20 | 0.5 | 0.5 | 20 | 20 | 0.5 | 0.5 | 0.50% | 0.125% |
| 4 | 20 | 20 | 0.5 | 0.5 | 20 | 20 | 0.5 | 0.5 | 0.50% | 2.000% |
| 5 | 20 | 20 | 20 | 20 | 20 | 20 | 5 | 5 | 0.50% | 0.50% |
| 6 | 20 | 20 | 20 | 20 | 20 | 20 | 2 | 2 | 0.50% | 0.50% |
| 7 | 20 | 20 | 0.5 | 0.5 | 20 | 20 | 0.125 | 0.125 | 0.50% | 0.50% |
| 8 | 20 | 20 | 0.5 | 0.5 | 20 | 20 | 0.05 | 0.05 | 0.50% | 0.50% |
| 9 | 20 | 20 | 0.05 | 0.05 | 20 | 20 | 0.005 | 0.005 | 0.50% | 0.50% |
| 10 | 20 | 20 | 0.0005 | 0.0005 | 20 | 20 | 0.00005 | 0.00005 | 0.50% | 0.50% |
| 11 | 20 | 20 | 0.000005 | 0.000005 | 20 | 20 | 0 | 0 | 0.50% | 0.50% |
| 12 | 20 | 20 | 0.000005 | 0.000005 | 20 | 20 | 5E-07 | 5E-07 | 0.50% | 0.50% |
| 13 | 20 | 20 | 5E-08 | 5E-08 | 20 | 20 | 5E-09 | 5E-09 | 0.50% | 0.50% |
| 15 | 20 | 20 | 0.5 | 0.5 | 20 | 20 | 0.5 | 0.5 | 0.50% | 4.000% |
| 16 | 20 | 20 | 0.5 | 0.5 | 20 | 20 | 0.5 | 0.5 | 15.00% | 40.000% |
| 17 | 0.5 | 0.5 | 0.5 | 0.5 | 0.5 | 0.5 | 20 | 20 | 100.00% | 60.000% |
| 18 | 20 | 20 | 0.5 | 0.5 | 20 | 20 | 0.5 | 0.5 | 0.50% | 0.000% |
| 19 | 20 | 20 | 2 | 2 | 20 | 20 | 0.125 | 0.125 | 0.50% | 0.50% |

Table: Particle diffusion and PopZ microdomain size parameters for Smoldyn files. Each file includes simulations for two different cells, labeled top and bottom. Y and R denote yellow and red colored particles, respectively, in bulk cytoplasm (cyto) and in PopZ microdomains (pole). PopZ microdomain size is expressed as a percentage of total cell volume.
